# Quantifying transcriptome turnover on phylogenies by modeling gene expression as a binary trait

**DOI:** 10.1101/2024.10.03.616564

**Authors:** Ammon Thompson, Michael R. May, Ben Hopkins, Nerisa Riedl, Olga Barmina, Benjamin J. Liebeskind, Li Zhao, David Begun, Artyom Kopp

## Abstract

Changes in gene expression are a key driver of phenotypic evolution, leading to a persistent interest in the evolution of transcriptomes. Traditionally, gene expression is modeled as a continuous trait, leaving qualitative transitions largely unexplored. In this paper, we detail the development of new Bayesian inference techniques to study the evolutionary turnover of organ-specific transcriptomes, which we define as instances where orthologous genes gain or lose expression in a particular organ. To test these techniques, we analyze the transcriptomes of two male reproductive organs, testes and accessory glands, across 11 species of the Drosophila melanogaster species group. We first discretize gene expression states by estimating the probability that each gene is expressed in each organ and species. We then define a phylogenetic model of correlated transcriptome evolution in two or more organs and fit it to the expression state data. Inferences under this model show that many genes have gained and lost expression in each organ, and that the two organs experienced accelerated transcriptome turnover on different branches of the Drosophila phylogeny.

## Introduction

Phenotypic evolution, broadly speaking, can result from three general mechanisms: gains and losses of genes; mutations in the coding sequences of genes, leading to changes in protein functions; and regulatory mutations, which lead to changes in gene expression. While the exact balance between these modes of genetic change in driving the origin of new phenotypes is a debated topic, it is clear that regulatory evolution plays a prominent role (King and Wilson, 1975; Hoekstra and Coyne, 2007; Carroll, 2008; Wray, 2007; Stern and Orgogozo, 2008; Courtier-Orgogozo et al., 2020). Across a diverse range of taxa and traits, changes in the expression of orthologous genes are pivotal in creating major phenotypic differences within and between species (e.g., Rebeiz et al. 2009; Fraser et al. 2010; Loehlin et al. 2019; Kowalczyk et al. 2022; Marand et al. 2023).

RNA sequencing (RNA-seq) technology has enabled quantification of gene expression levels for the entire transcriptome across diverse biological conditions and developmental stages (Wang et al., 2009; Mortazavi et al., 2008; Hrdlickova et al., 2017; Marguerat and Bähler, 2010). As a result, RNA-seq has become essential for studying regulatory evolution, providing comprehensive and quantitative data for inferring large- and small-scale changes in transcriptomes associated with adaptive evolution (Wittkopp and Kalay, 2012; Harrison et al., 2012; Todd et al., 2016). Transcriptome-level studies of gene expression evolution have largely focused on quantitative differences between species, modeling a gene’s expression as a continuous trait. Indeed, most evolutionary changes in gene expression are of a quantitative nature (Rifkin et al., 2003; Brawand et al., 2011; Harrison et al., 2012; Romero et al., 2012; Coolon et al., 2014; Nourmohammad et al., 2017; Cardoso-Moreira et al., 2019). By contrast, qualitative changes, manifested as discrete gain and loss of gene expression in a particular tissue over evolutionary time (“transcriptome turnover”), have received comparatively little attention (Mika et al., 2021, 2022). Consequently, questions about the rates of transcriptome turnover, the consistency of these rates over time and across different evolutionary lineages, and the degree of correlation in turnover rates between different organs remain unexplored. This may be an important deficit in our understanding of phenotypic evolution given the many examples where gain or loss of gene expression in an organ has a profound impact on its structure or function (Glassford et al., 2015; Hu et al., 2018; Kellenberger et al., 2023; Molina-Gil et al., 2023).

Studying gene expression at a qualitative level brings with it two significant advantages. The first is biological. Relative to quantitative changes, deactivation of a gene, or activation of an ancestrally inactive gene, in a particular organ might be expected to have a more profound impact on phenotypic evolution (Harlin-Cognato et al., 2006; Carroll, 2008; Wray, 2007; Thompson et al., 2016, 2018; Tanaka et al., 2011; Glassford et al., 2015; Martinson et al., 2017; Hu et al., 2018). The generality of this idea, which is currently supported by anecdotal examples, needs to be tested systematically at the genome-wide level. The second advantage is technical. If inferred reliably, discrete expression states should be less sensitive to technical factors such as sequencing depth and library size, differences in cell type composition caused by dissection variability, and the well-documented problems of comparing compositional values such as TPM or FPKM between tissues and species (Brawand et al., 2011; Dillies et al., 2013; Li et al., 2014; Musser and Wagner, 2015). While the effect of some of these variables on cross-species and cross-tissue quantitative analyses can in principle be minimized through normalization approaches, selecting an appropriate evolutionary model for complex transformations and normalization schemes based on, for example, reference gene sets can be challenging to implement and interpret (Vandesompele et al., 2002; de Jonge et al., 2007; Brawand et al., 2011; Dillies et al., 2013; Chen et al., 2014; Cardoso-Moreira et al., 2019; Zhou et al., 2019; Quinn et al., 2018; Dimayacyac et al., 2023; Mantica et al., 2024).

The main obstacle to investigating transcriptome turnover on a genome-wide scale is the difficulty of translating the continuous RNA-seq data (TPM or FPKM) into discrete (ON/OFF) gene expression states. Many genes that play important roles in development can be expressed at low levels, making their expression difficult to detect reliably. Conversely, some genes that are not actively expressed in a particular tissue may nevertheless be detected at non-zero levels in a transcriptome sample due to “noisy” transcription that occurs throughout the genome or technical artifacts of the sequencing pipeline (Kin et al., 2015; Singh and Petrov, 2007; Wagner et al., 2013; Jensen et al., 2013; Struhl, 2007; Schwanhäusser et al., 2011; Artieri and Fraser, 2014; Lenive et al., 2016; Ham et al., 2020). Historically, gene expression states in transcriptomes have been discretized using hard expression thresholds; for example, genes present above 1 - 3 TPM may be considered actively expressed, while those below 1 TPM are assigned as inactive (Mortazavi et al., 2008; Hebenstreit et al., 2011; Wagner et al., 2013; Huang et al., 2015; Cridland et al., 2020; Mika et al., 2021, 2022). In some cases, specific expression cut-offs are well justified on mechanistic biological grounds implied by chromatin marks associated with active or repressed transcription (Ernst et al., 2011; Hart et al., 2013; Singh and Petrov, 2007; Wagner et al., 2013). However, fixed cut-offs are poorly suited to cross-species and cross-tissue comparisons, since it is far from clear that a single expression threshold can accurately distinguish between active expression and transcriptional noise in all genes, species, and tissues, especially when the species are distantly related and/or the tissues differ strongly in transcriptome complexity. In fact, our recent work has shown that the boundary between active expression and noise is different even among tissues that are physiologically similar (e.g., different primate brain regions; Thompson et al., 2020).

To develop a more objective and generally applicable approach to discretizing gene expression states, we recently created and validated a method, *zigzag* (Thompson et al., 2020), for inferring gene expression states from replicated RNA-seq datasets. *zigzag* uses Markov chain Monte Carlo to estimate the posterior probability of active expression (the “ON” state) for each gene in each tissue using a well-defined statistical model and testable prior assumptions, enabling easy validation. *zigzag* achieves this by learning universal landmarks in transcriptome datasets that distinguish between active and inactive genes (Hebenstreit et al., 2011; Wagner et al., 2013; Hart et al., 2013; Huang et al., 2015; Thompson et al., 2020; Costa et al., 2022). This method was shown to be sensitive enough to correctly classify expression states of transcription factors expressed in only a few cells in the drosophila testes while classifying smell and taste receptor genes as largely inactive in the same tissue despite detecting reads mapping to these genes (Thompson et al. 2020). The ability to classify gene expression states probabilistically using *zigzag* opens the way for investigating the evolutionary turnover of tissue-specific transcriptomes (i.e., the transition of conserved genes between OFF and ON expression states in each tissue) using well-established phylogenetic models of discrete-trait evolution.

In this study, we present a pipeline that integrates zigzag with phylogenetic comparative methods to infer the evolutionary dynamics of transcriptome turnover. To test this pipeline, we used RNA-seq datasets from two male reproductive organs, testes and seminal fluid-producing accessory glands (AGs), across 11 Drosophila species. We chose these organs as the test subject for our approach because previous studies have shown that the male reproductive system evolves faster than other tissues at the level of both protein sequence (Begun et al., 2000; Kern et al., 2004; Haerty et al., 2007; Patlar et al., 2021) and gene expression (Meiklejohn et al., 2003; Ranz et al., 2003; Rifkin et al., 2003; Zhang et al., 2007; Pal et al., 2023), suggesting that we may be able to estimate the rate of transcriptome turnover even with a limited number of taxa.

With this analysis pipeline we were able to estimate the rate of transriptome turnover for each organ, how much those rates correlate between the two organs, the degree of punctuated evolution or “burstiness” of turnover, and how much rates vary among different gene families and functional categories. Our results suggest that qualitative gains and losses of gene expression are fairly common in these organs, and that the rates of turnover vary over time. We discuss the possible implications of widespread activation and silencing of genes for organ evolution. We also discuss a number of important challenges and pitfalls associated with this approach.

## Results

### Transcriptomes of Testes and Accessory Glands in 11 *Drosophila* Species

We sequenced the transcriptomes of testes and accessory glands in 11 species of the *Drosophila melanogaster* species group (see Materials and Methods). We used *zigzag* (Thompson et al., 2020) to assign genes to active or inactive expression states in the testes and accessory glands of each species under multiple probability thresholds. *zigzag* jointly estimates a probability of active expression for all genes thus producing a distribution of gene-specific marginal posterior probabilities of active expression. All genes with probabilities above an upper threshold were classified as active while those below a lower threshold were considered inactive. As an example, Figure 1 (top) shows the estimation performed for *D. melanogaster* at P *<* 0.5 for the inactive and P *>* 0.5 for the active state. Inference under a *zigzag* model suggests that a larger proportion of the genome is expressed in the testes compared to the AG (Figure 2), which supports recent findings (Cridland et al., 2020). The inferred proportion of expressed genes is 46–64% in the AG and 65–75% in the testis (lowest 2.5% quantile to highest 97.5% quantile among species posterior distributions). In most cases, a higher proportion of single-copy genes (gene families containing a single gene in all species) are actively expressed, compared to the rest of the genome (Figure 2). The single-copy set is likely enriched for housekeeping genes, which are broadly expressed and tend to have less copy number variation in populations than other genes (Dopman and Hartl, 2007; Henrichsen et al., 2009).

**Figure 1:**
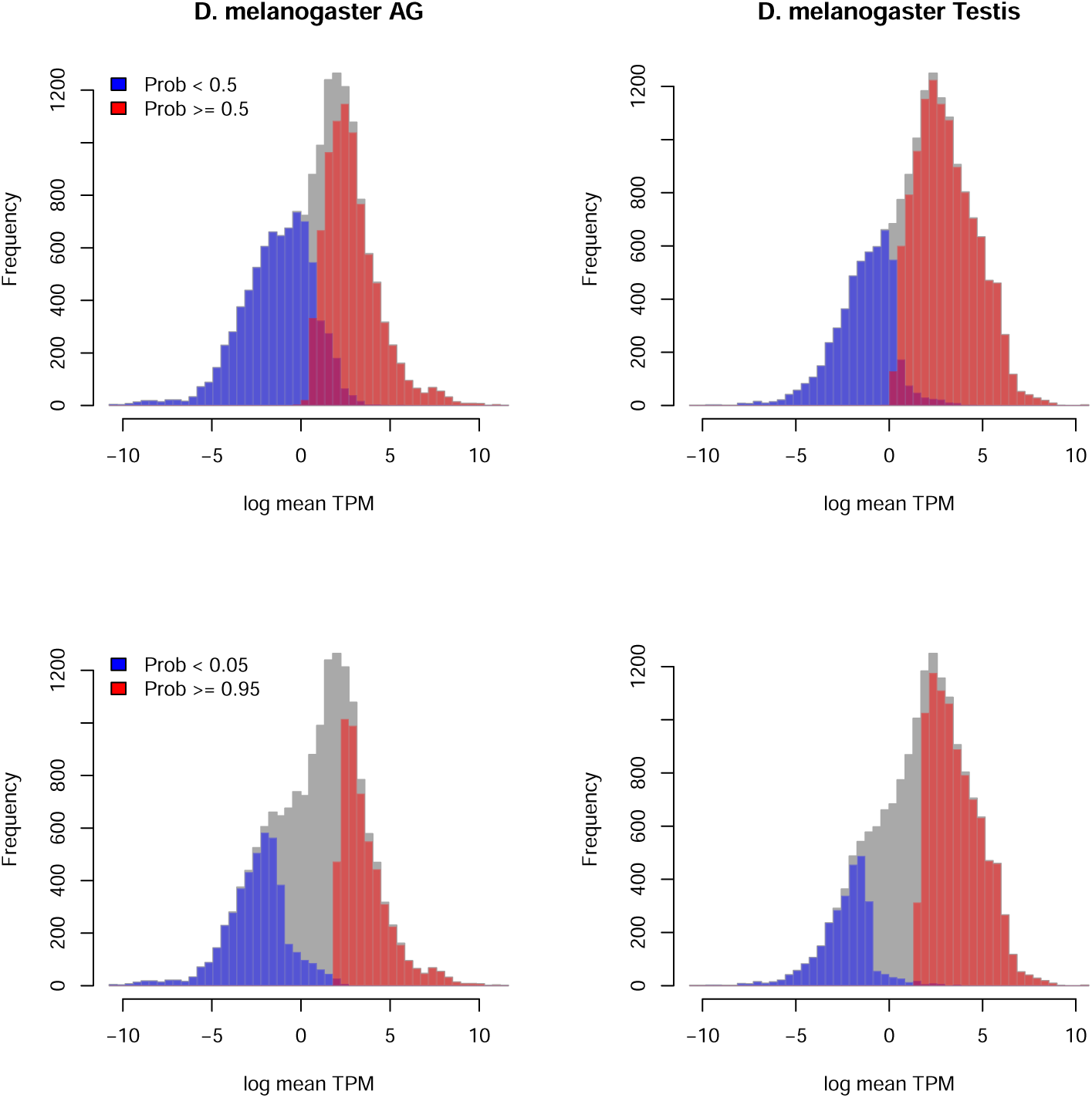
Probabilistic estimation of discrete expression states from continuous RNA-seq data using *zigzag* at two different probability cutoffs. Top row: genes with posterior probabilities of active expression *P <* 0.5 are assigned to the inactive state (blue), while those with *P >* 0.5 are assigned to the active state (red). Bottom row: *P <* 0.05 are inactive and *P >* 0.95 are active.; genes with intermediate probabilities are not assigned to either active or inactive states. Similar estimation was performed for all species, separately for each tissue, under a range of probability thresholds. Gray shows the combined frequency of genes classified as active, inactive or neither. As the probability cut-offs become more stringent, the overlap between active and inactive distributions is reduced, but a larger fraction of genes remain unclassified

**Figure 2:**
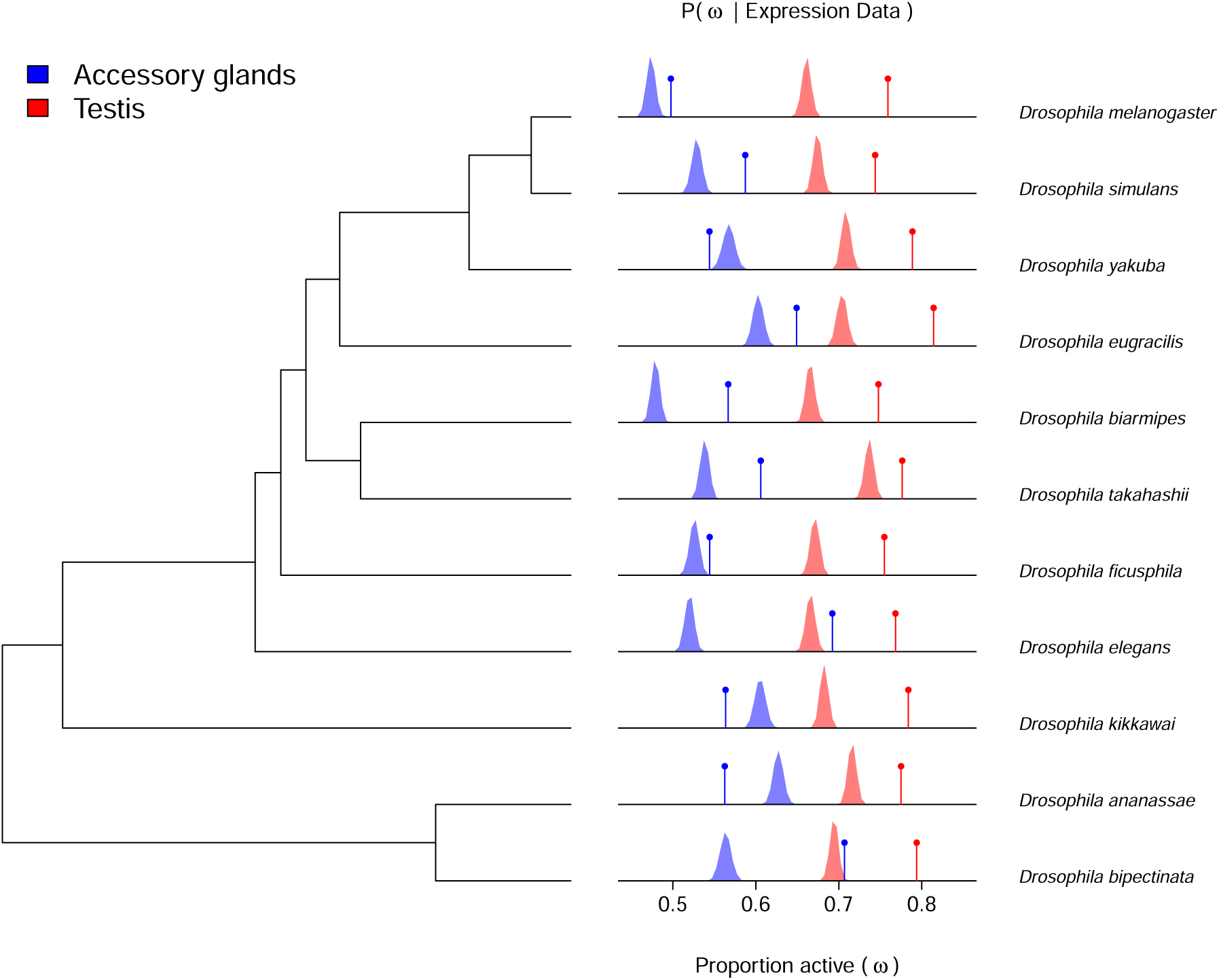
Posterior distributions of the weight active parameter in the *zigzag* model (*ω*), which measures the proportion of all protein-coding genes that are actively expressed in the AG (blue) and testes (red) of each species; species phylogeny is shown on the left. Vertical lines indicate the expected proportion of single-copy genes (out of n=8660) that are active in each organ, calculated from the posterior probabilities of active expression of each gene in each species. If the expression states of genes were, on average, the same for single-copy and non-single-copy genes, the peaks of the weight active posterior distributions would coincide with the vertical lines. In most cases, however, a higher proportion of single-copy genes is estimated to be active, compared to the total protein-coding genome.

The total abundance of transcripts coming from inactive genes is similar in the two organs. The combined expression of genes with less than 0.5 probability of being active contributed less than 1% of the total mRNA in each transcriptome. The model fit to the data suggests that transcripts from inactive genes comprised 0.1–0.6% of the AG transcriptome and 0.1–0.4% of the testis transcriptome. We estimated the threshold level of expression at which a gene is, on average, as likely to be active as inactive (probability of active expression P = 0.5). Depending on the species, this threshold ranges from 0.6 to 2.0 TPM in the AG and 0.8 to 1.6 TPM in the testis. These estimates, which are similar to those obtained by other methods in other tissues and organisms (Costa et al., 2022; Wagner et al., 2013), confirm that no single “hard” cut-off of TPM or FPKM values would be equally appropriate for all species and tissues (Thompson et al., 2020).

Inference under the *zigzag* model supports previous findings that the testis and AG transcriptomes are highly overlapping (Cridland et al., 2020). Under a more stringent threshold where genes with probability of expression between 0.1 and 0.9 were classified as unknown, we found that the AGs and testes share between 5,330 and 7,318 actively expressed (P *>* 0.9) genes, depending on the species. The two organs differ in the number of exclusive genes, i.e., those actively expressed in one but not the other organ. The AGs express between 68 and 200 exclusive genes per species, while the testes are more variable with 301–1,939 exclusive genes per species (Table 1). This supports previous studies in *D. melanogaster*, *D. yakuba*, and *D. simulans* showing that the testes express an elevated number of tissue-specific genes relative to other tissues, including the AGs (Takashima et al., 2023; Cridland et al., 2020). These are likely minimum estimates due to our conservative threshold for active and inactive expression calls. Our *zigzag* analyses confirm that the set of genes that are active in a transcriptome vary not just among organs but between species (Figure 3).

**Figure 3:**
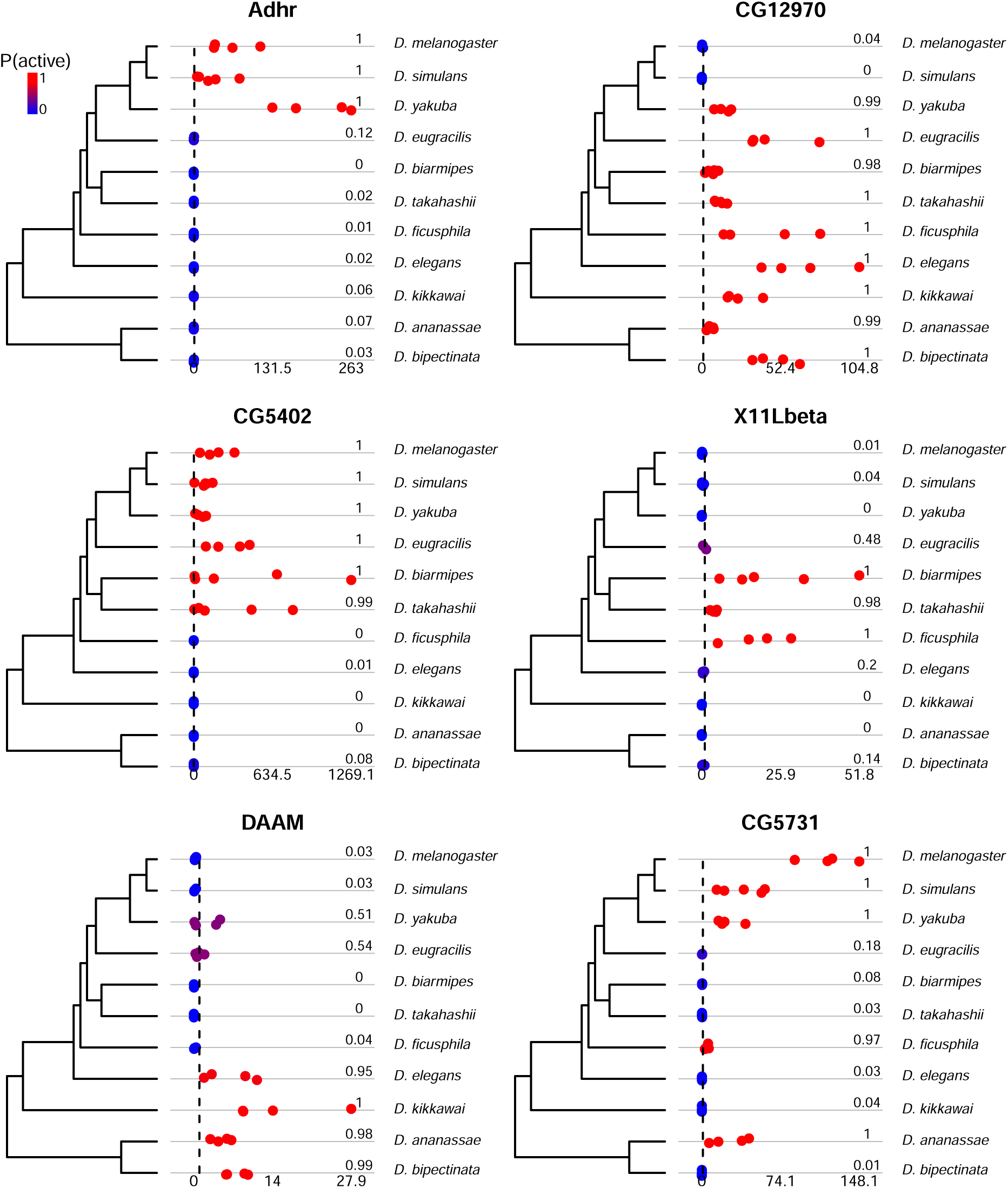
Comparative expression levels and expression state estimates for 6 genes in accessory glands that show one or more evolutionary changes in expression state among the 11 species. Dot plots show the estimated TPM values for each of the 4 to 5 biological replicate RNA-seq libraries in each species, with the range of TPM values indicated under the dot plots. TPM = 1 is indicated by a vertical dashed line. Hot and cold color gradient and numbers to the right of the dots show posterior probability of active expression under the *zigzag* model.

**Table 1:** The transcriptomes of testes and accessory glands.

|  | AG | Testis |
| --- | --- | --- |
| % active protein coding genes | 46–64% | 65–75% |
| % transcripts from inactive genes | 0.1–0.6% | 0.1–0.4% |
| TPM of genes with $P = 0.5$ active | 0.6–2.0 TPM | 0.8–1.6 TPM |
| Number of exclusive expressed genes | 68–200 | 301–1,939 |

### Expression of X-linked Genes in Male Reproductive Organs

Previous studies have found that the X chromosome is depleted for genes with male-biased expression (Parisi et al., 2003; Ranz et al., 2003; Mueller et al., 2005; Sturgill et al., 2007). In the genus *Drosophila*, genes with testis-biased expression show a tendency to move from the X to the autosomes (Vibranovski et al., 2009), while accessory gland proteins (Acps) are significantly underrepresented on the X chromosome (Ravi Ram and Wolfner, 2007) in *D. melanogaster*. In these studies, tissue-specific (or tissue-enriched) genes were identified on the basis of expression bias, i.e., quantitative difference in transcript abundance between different organs or between males and females. Interestingly, our approach, which classifies genes by presence or absence of expression in each tissue separately, highlights a different pattern. Our estimates of expression state indicate that X-linked genes are more likely to be expressed in both testes and AGs, compared to the genome average (Figure 4).

**Figure 4:**
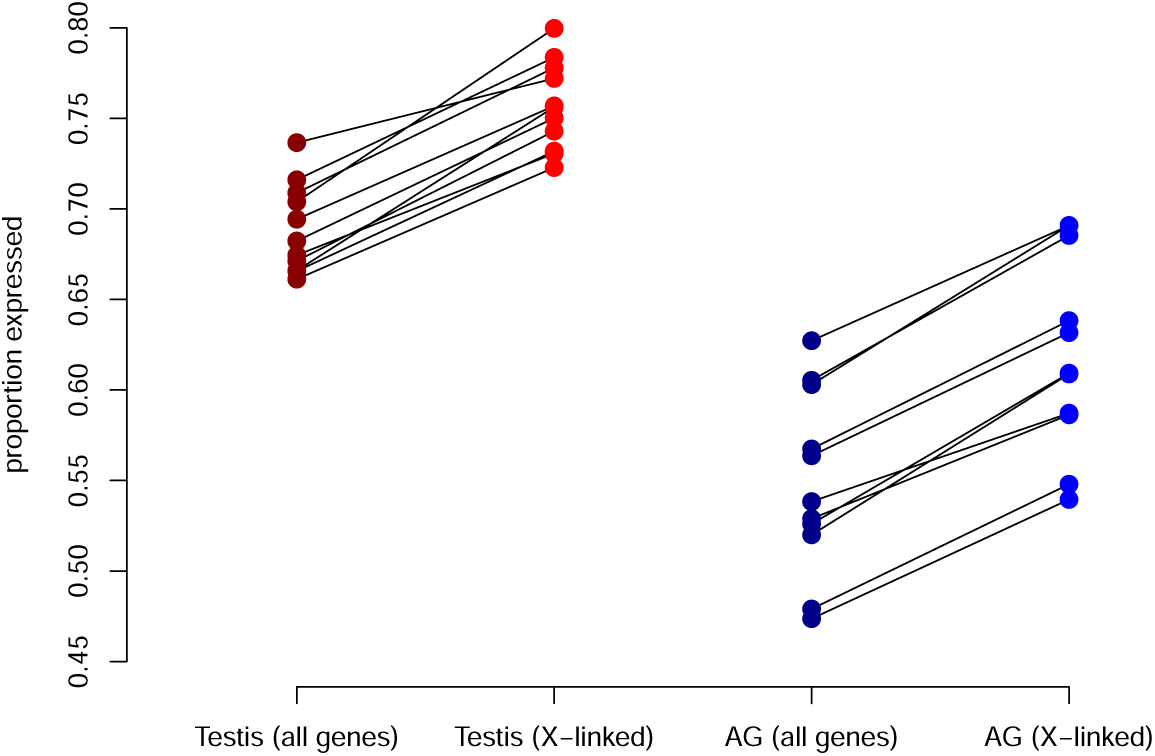
The proportion of genes expressed in the testis and accessory glands. Each colored dot shows the expected (averaged over posterior probability of active expression) proportion of genes expressed in each species and organ. Lines connect the “all genes” (left) and “X-linked” (right) categories within each species.

### Phylogenetic Model of Transcriptome Turnover

To investigate the evolutionary processes that contributed to the patterns we observe in our data, we fit a model of transcriptome evolution in the two organs to a time-calibrated phylogenetic tree and the expression states for single-copy genes in both organs (see Methods). In brief, we assumed that each gene’s expression state evolved as a two-state (active/inactive) continuous-time Markov chain (CTMC) with turnover rate (rate of activation/deactivation of expression) varying across branches according to an uncorrelated relaxed clock model where each branch of the tree was assumed to have a unique mean turnover rate that correlated between the two organs. We also included among-gene rate variation in our model to account for differences in turnover rate among genes.

Ideally, we would average over the uncertainty of the expression states of all genes while fitting evolutionary models to the *zigzag* posterior probabilities. Unfortunately, such phylogenetic analysis methods are not currently available for transcriptome-scale comparative data, which involve thousands of characters. We therefore set two probability thresholds to classify genes as active (high probability of active expression), inactive (low probability of active expression), or unknown/missing data (intermediate probability of active expression). Because, to our knowledge, this type of study has not been performed before, it was important to investigate the consistency, adequacy, and robustness of not just the *zigzag* predictions of expression state, but also of the evolutionary model fit to those inferred expression states. The probability cutoffs for active and inactive genes in particular can potentially have a large impact on inferences. If these cutoffs are too conservative, too much data will be thrown out and the prior will have a strong impact on our inferences. As the cutoffs approach 0.5, more of the data is used for inference but the number of misclassifications increases, which could overpower the true signal in the data.

To assess the overall adequacy of our evolutionary model and to select appropriate probability cutoffs for gene expression states, we conducted a series of cross-validation experiments and sensitivity tests (see Methods and Supplemental Text S1). The results of this cross-validation experiment revealed that the probability thresholds of *α* = 0.05, 0.1, and 0.25 work well for our data such that phylogenetic inferences of the expression states of the hold-out data agreed with the *zigzag* estimates with probability 0.88 on average. Accuracy was similar for each species (see Supplemental text S1, figures S5 and S6). We performed phylogenetic analyses under all cutoffs, and mostly report the estimates based on an *α* = 0.1 as our most confident estimates since that level had the highest average accuracy.

### History and Dynamics of Transcriptome Turnover

Inferences under our model suggest distinct modes of evolution for the two organ’s transcriptomes. Our model allowed us to directly infer the correlation of branch-specific rates of transcriptome turnover in the two organs. Despite there being a great deal of uncertainty about the strength and direction of this correlation (95% highest posterior density interval (HPD) for the correlation parameter is −0.23, 0.65), the correlation is likely no more than 0.65.

We estimated the rates at which genes were turned on and off in the AG and testis and explored how those rates varied among genes and organs as well as among the different branches of the species phylogeny. Assuming that the root age of the species tree is approximately 25 million years (Obbard et al., 2012), our posterior estimates of branch rates suggest that the accessory glands experienced a pulse of rapid transcriptome turnover around 10 million years ago, near the base of the Oriental lineage of the *melanogaster* species group (Figure 5). In contrast, we infer much more subdued rates of turnover in the testis on those branches. Also, in contrast to the AG, we infer accelerated transcriptome turnover in the testis on the *D. melanogaster* tip branch, compared to the rest of the tree (Figure 5). For probability cutoffs (*α*) of 0.05, 0.1, and 0.25, the relative rates of turnover were fairly robust to the choice of probability cutoff, with branches 15, 16 and 17 at the base of the Oriental lineage consistently showing the highest rates of turnover for accessory glands, while some terminal branches consistently show elevated turnover in testes, in particular in *D. melanogaster* (Supplemental text S1 Figures S1–S4).

**Figure 5:**
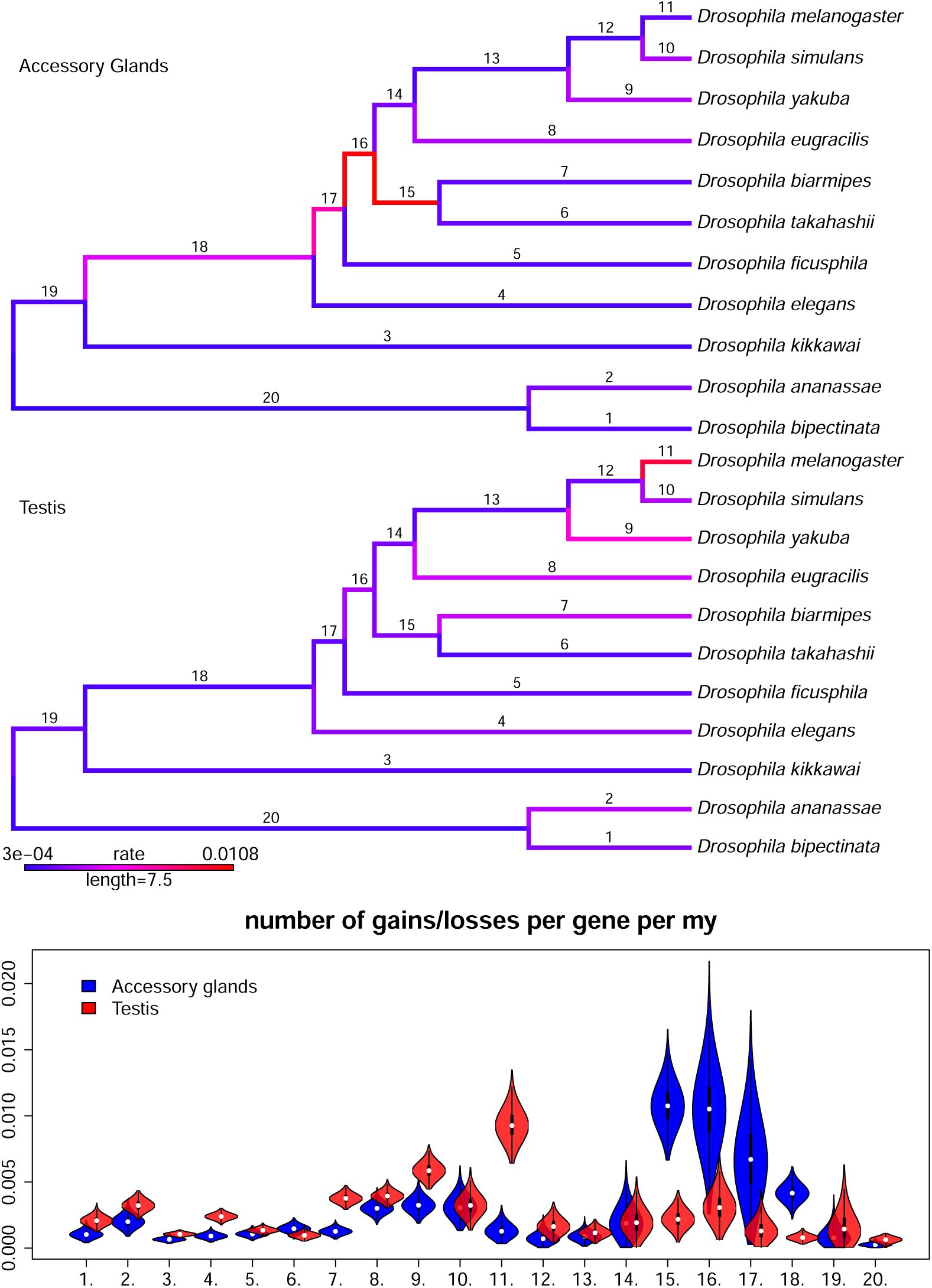
Posterior mean estimations of branch-specific relative turnover rates for single-copy genes in the AG (top phylogeny) and testis (bottom phylogeny). Branch color reflects the mean posterior estimates of the transcriptome turnover rate (changes per single-copy gene per MY), from low (blue) to high (red). The bottom violin plot shows the full posterior samples of the relative turnover rate for AG (blue) and testis (red) for each branch numbered on the phylogenetic trees. Note the accelerated turnover on branch 11 in the testis and branches 15-18 in the AG.

At the cutoff of *α* = 0.1, we estimate that the mean gene expression turnover rate is 1.8 *×* 10*^−^*^3^ per million years in the AG and 2.0 *×* 10*^−^*^3^ per million years in the testis (Table 2; Data file S2). Overall, the per-gene turnover rate appears to be on the order of 10*^−^*^3^ per MY (Table 2). For reference, this is on the same order of magnitude as the rate of nucleotide substitutions in *Drosophila*, where estimates are around 8 *×* 10*^−^*^3^ per MY (Obbard et al., 2012). Thus, for a gene of size 1 kb, we expect it to change expression state on the order of once every 1,000 base changes. For a genome of around 15,000 genes, these turnover rates imply that roughly 40 expression activation/deactivation events happen in each organ per million years. However, this does not imply a massive rewiring of the transcriptome. Most expression turnover is likely driven by the frequent change in the expression state of a relatively small subset of genes (Figure 6). In the accessory glands, we estimate that half of all transcriptome turnover among single-copy genes is being driven by about 11% of the genes, while about 15% of genes are driving half of all changes in the testis. Since these estimates come from single-copy genes, which appear to be more likely to be expressed in the AG and testis compared to the rest of the genome (Figure 2), these rates of turnover probably differ from the genome average and, we suspect, underestimate the average rate for all genes. On the other hand, transcriptome turnover may be slower in other tissues compared to the rapidly evolving male reproductive system. In summary, our analyses imply that the rate of transcriptome turnover changes over time independently in the two organs and is highly variable among genes.

**Table 2:**
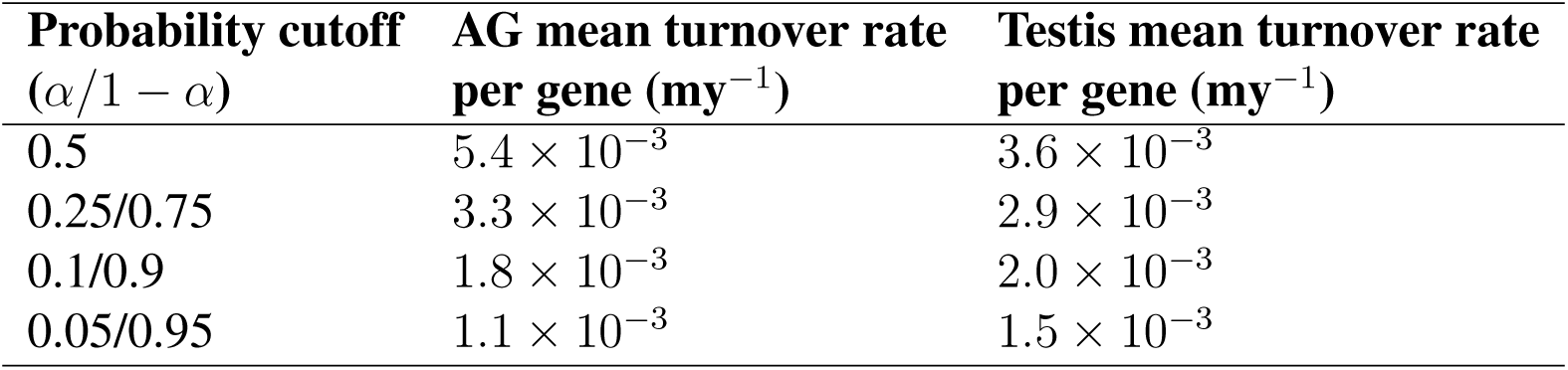
Mean turnover rates per gene for AG and testis at different probability cutoffs (*α*).

**Figure 6:**
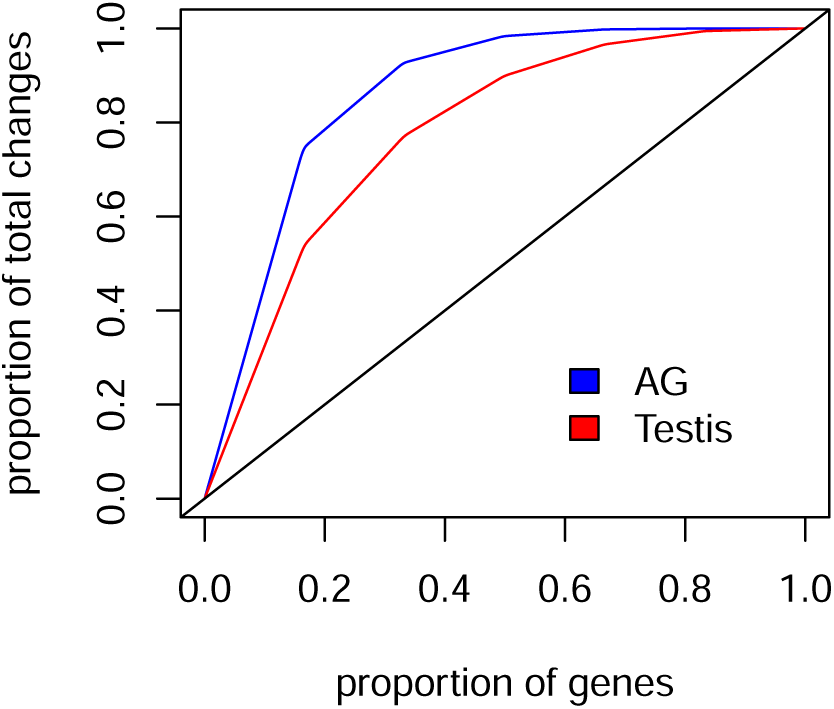
Transcriptome turnover is concentrated among a minority of genes. Proportion of single-copy genes (n = 8660) are ranked on the X axis by their rate of change, from fastest to slowest evolving. The red (testis) and blue (AG) curves show the cumulative proportion of transcriptome turnover events explained by each subset of genes. For example, 20% of genes account for nearly 80% of all turnover in the AG. The black line shows the expected curve if all genes turned over at the same rate.

We tested whether the genes that show elevated turnover rates in male reproductive organs are enriched for particular biological functions. To do this, we performed gene ontology (GO) analysis on the sets of genes that showed turnover rates at least two-fold higher than the mean. This analysis was done separately for AGs and testes, using all singleton genes expressed in the corresponding tissue as the background. 8 GO terms in the testis, and 19 in the AG, showed significant enrichment (Fig 7 A, B; and Figure S7 in Supplemental text S1). There was strong overlap in the significant terms between the two tissues, such that all gene categories that were enriched in the testis were also enriched in the AG. However, there was only a limited overlap between the individual genes driving the enrichment (Figure 7 C). The enriched GO terms were related to sensory perception (including olfactory and gustatory receptor genes and odorant-binding proteins), GPCR signaling (especially neuropeptide signaling), and cuticle development. The AG additionally showed enrichment for terms related to cilium organization and cell-cell adhesion.

**Figure 7:**
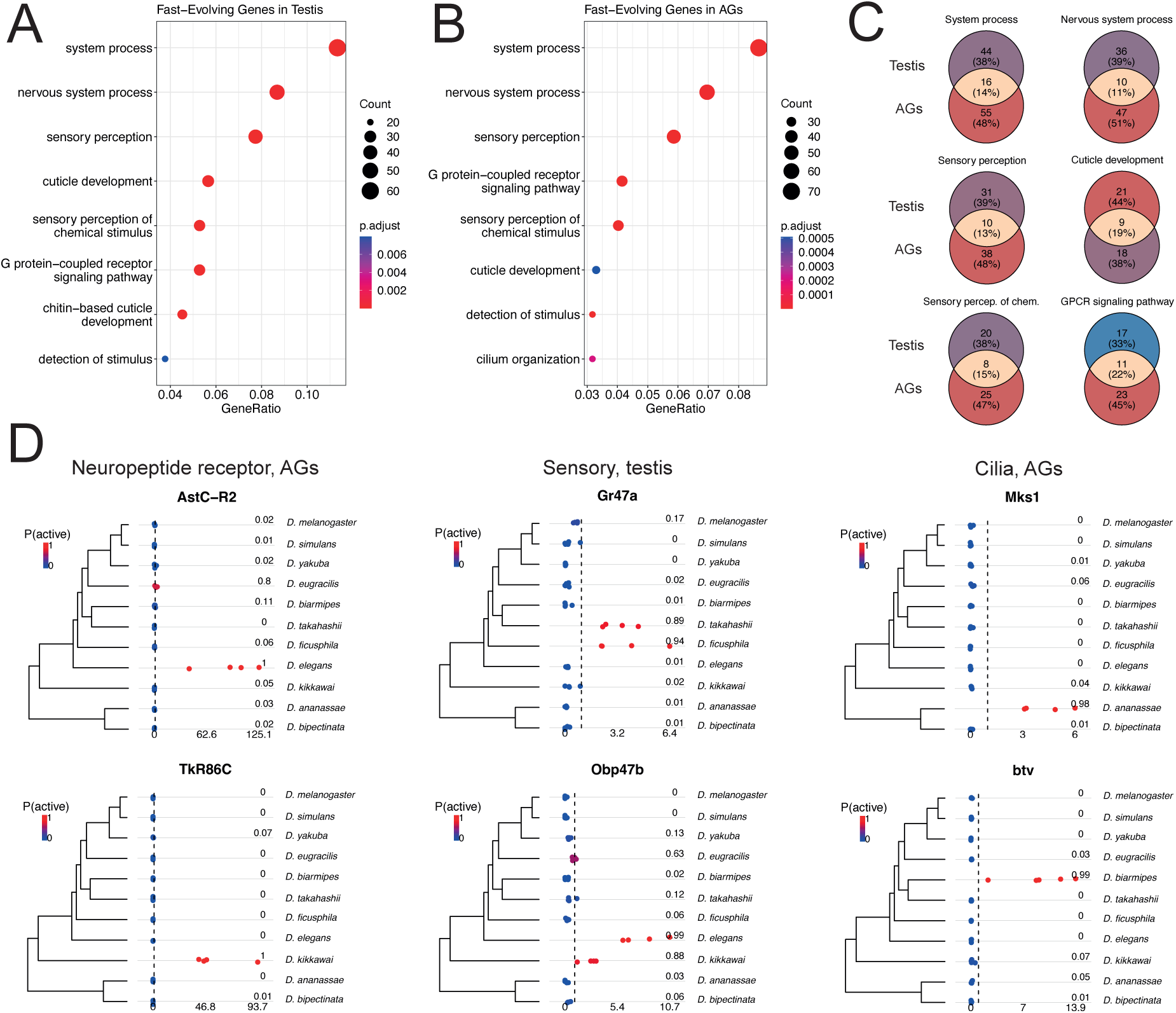
Genes with elevated turnover rates are enriched for similar functional categories in testes and AGs. (A) A dot plot from a gene ontology (GO) analysis showing all significantly enriched (q-value *<* 0.05) biological process terms among the genes showing rapid turnover in the testis. Genes are designated as showing rapid turnover if they have rates at least two-fold higher than the mean. The list of all singleton genes expressed in the testis is used as the background. (B) The top 8 significantly enriched terms in the AG; analysis performed the same way as in (A). The full list of significantly enriched AG terms can be found in Figure S7 of Supplemental text S1. (C) Venn diagrams illustrating the overlap between the genes showing rapid turnover in the testis and AG, in each of the shared enriched GO terms. Note that most individual genes are exclusive to one organ. (D) Dot plots showing the TPM values for selected genes showing high turnover rates, mapped onto the species phylogeny. Two genes per functional class (neuropeptide receptors, sensory perception, and cilium organization) are shown. For each pair, both genes are shown in the same organ. Dots are colored in accordance with the posterior probability of active expression inferred from the zigzag model. Separate dots within a species represent different biological replicates.

The pattern of unusually fast turnover could potentially be explained by contamination of some samples with RNA from non-target tissues. For example, the presence of genes related to cilium organization in AG samples could reflect contamination from sperm, transcripts related to cuticle development could be coming from epithelial tissues, and genes related to GPCR signaling and sensory perception may reflect contamination from neurons that innervate the reproductive organs. However, several lines of evidence argue against contamination being a major factor. First, other genes that are highly expressed in the potential sources of contamination (such as ion channels and sperm-specific genes) do not show up in the enrichment analysis. Second, the inferred gains of expression are generally consistent across replicates of the same species (Figure 7 D). Third, these gains are often associated with high TPM counts (Figure 7 D). Fourth and most important, different genes in the same GO category show expression gains in different species (Figure 7 D), whereas contamination would produce a highly correlated gain. For these reasons, contamination is unlikely to be a major contributor to the rapid transcriptome turnover we observe.

We do not know what roles, if any, the genes undergoing rapid turnover play in male reproductive organs, but some enriched terms are consistent with their physiology. Neuropeptides are known to regulate copulation, for example, by coupling mating duration to the transfer of sperm and seminal fluid (Taylor et al. 2012). Similarly, the ability of males to upregulate sperm production in response to the presence of females depends on the ability of somatic cyst stem cells of the testis to respond to neuronally secreted octopamine (Martin-Diaz and Herrera, 2024). It is conceivable, therefore, that co-option of genes involved in neuropeptide signaling could lead to lineage-specific changes in inter-organ communication or the regulation of secretory cell activity – changes that may, for example, have implications for the males’ ability to dynamically respond to the sociosexual environment (e.g., Hopkins et al. 2019).

We next investigated two plausible artifacts that could cause the inferred patterns of branch rate variation in these two organs. One obvious cause could be an artifact of our method of classifying genes as ON or OFF. Our inferences could be highly sensitive to misclassification, which could lead to higher estimates of turnover rates near the tips of the tree. The second scenario involves a subset of genes exhibiting a very high rate of turnover, such that phylogenetic information is quickly erased to different degrees in different parts of the tree, e.g., longer branches vs. shorter branches near the tips. To explore these potential sources of error, we simulated two datasets under these scenarios using *sim.char* in the R package *geiger* v 2.0.11 (Pennell et al., 2014), with the same number of genes and species and the same tree as the *Drosophila* dataset. In both simulations, activation rate equaled deactivation rate and branch rates were held constant in order to test whether these potential problems could lead our method to infer significant among-branch rate variation where none exists. In the first simulation, all genes evolved at the same rate, except 5% of genes evolved 100-fold faster. In the second simulation, there was no variation in rates among genes, but 1% of the genes in each species and organ had their expression state flipped to simulate misclassification. Figure 8 shows that the branch rates at shorter terminal branches in our model are sensitive to both these scenarios. Thus, the elevated rates observed on branches 1, 2, 9, 10, and 11 may be caused in part by state misclassification, a very high turnover rate in a subset of genes, or both.

**Figure 8:**
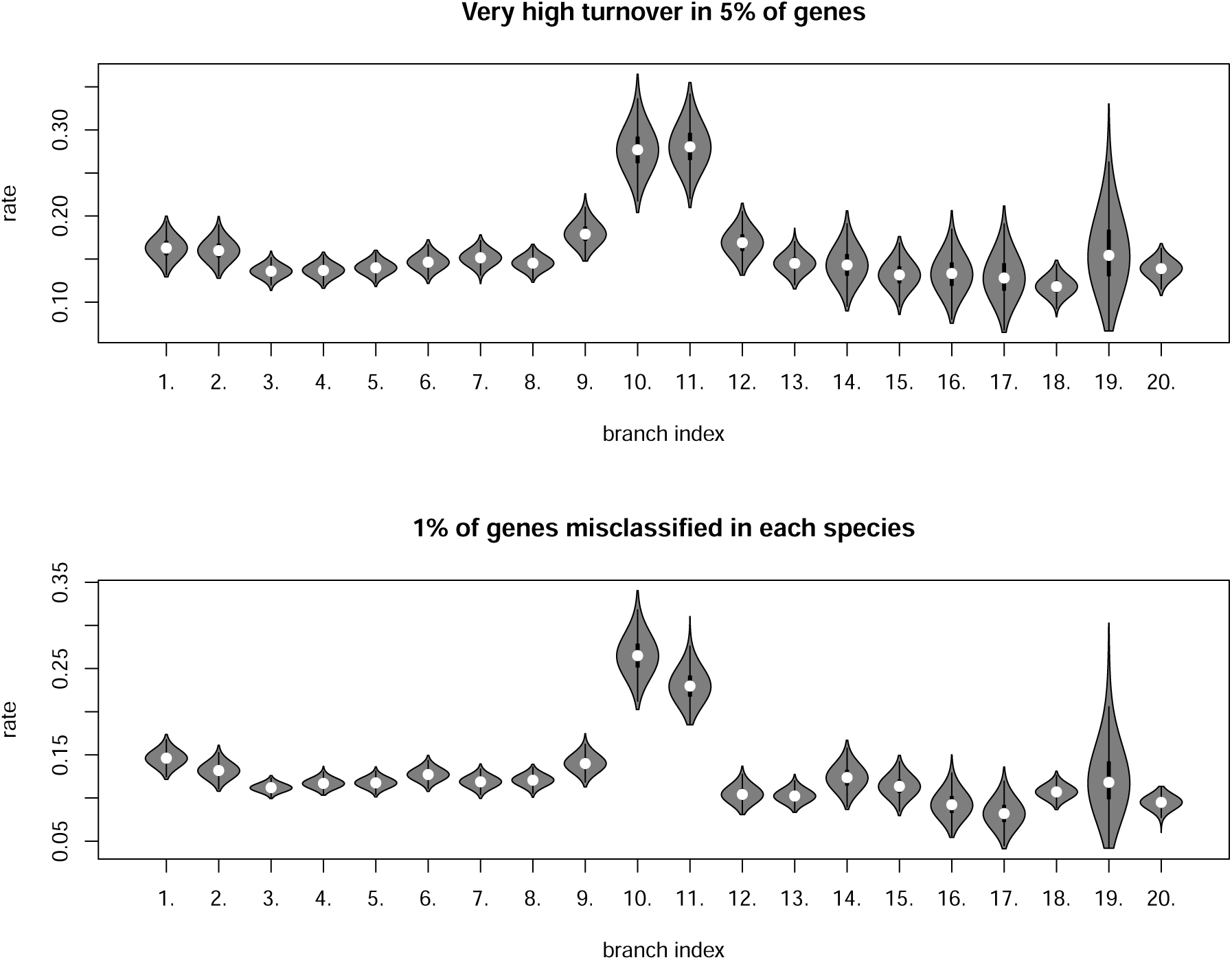
Posterior distributions of branch rates for data simulated under two scenarios where the true branch rates are constant. See Figure 5 for branch numbers. Top panel shows inferred branch rates when 5% of genes change expression states 100 times more rapidly than other genes. Bottom panel shows the inferred branch rates when 1% of genes have their expression states misclassified. Note that branches 10 and 11 are the shortest terminal branches in the tree.

### Turnover of Transcription Factors

Differences in the rate of expression evolution among classes of genes can suggest possible mechanisms for phenotypic evolution. In particular, gain or loss of a transcription factor expression can have large downstream consequences on the transcriptome and thus on the function of an organ. If a transcription factor is lost or gained by a transcriptome it may bring with it downstream targets, resulting in a pulse of transcriptome turnover in a lineage on the phylogeny. We examined the mean difference in the turnover rate between transcription factor genes and the rest of the single-copy gene families in our phylogenetic study. Our data and models imply that transcription factors change expression states at different rates in the accessory glands and testes, where they evolve at a relatively slow rate in the former and a relatively fast rate in the latter. We report the posterior credible intervals of the percent difference of the mean rate (PDMR) between each group of genes (i.e., TF and non-TF):

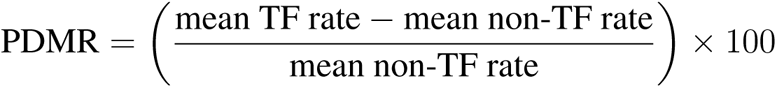

Our model and the RNA-seq data imply that expression states of single-copy transcription factors in AGs turn over on average 12–35% slower than those of non-TF single-copy genes. Conversely, TF expression states in testes turn over 0–17% faster compared to non-TF genes (colored lines and circles in each histogram in Figure 9; top row). To check how frequently random subsets of genes deviate from the genome average at least this strongly, we measured the mean PDMR (M-PDMR) for each comparison and created an empirical null distribution from random partitions of the data where we summarized each random partition with the same statistic, the M-PDMR (Figure 9; bottom row). Results show that for TF genes, this statistic is extreme relative to the reference set in both AGs and testis, with less than 0.1% of genes in the reference set having more extreme statistics for both organs. Thus, at the level of discrete ON/OFF transitions, our analysis implies that TF genes evolve slightly faster than non-TF genes in the testis, but significantly slower than non-TF genes in the AG.

**Figure 9:**
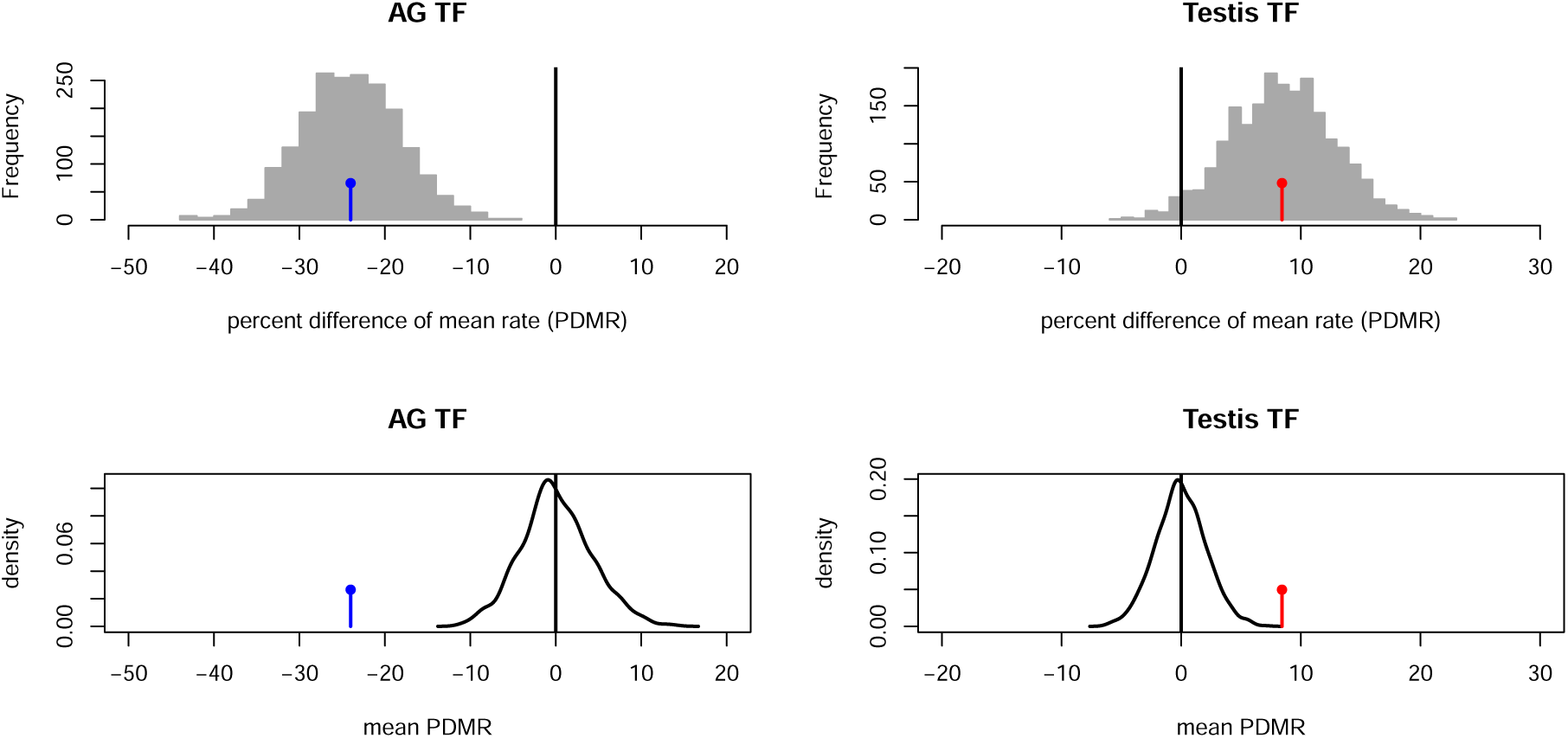
The posterior distributions of the percent difference of mean rate (PDMR) between transcription factors and non-transcription factors (top row). For example, the histogram in the top left shows the mean rate of the TF genes minus the mean rate of non-TF genes divided by the mean non-TF rate computed for each MCMC sample (n=2,000). The colored lines topped with a circle indicate the mean of the posterior distribution of PDRM. The bottom row shows reference distributions of mean PDMR (M-PDMR) of 1,000 random partitions of all genes into two groups equivalent in size to the TF/non-TF partition (black lines). The mean PDMRs of the TF groups (colored lines) are extreme relative to the control distributions.

### Turnover of X-linked Genes

X-linked genes may evolve differently from autosomal genes due to the differences in selective pressures and effective population size that the X chromosome experiences in males vs females (Parisi et al., 2003; Ravi Ram and Wolfner, 2007; Singh and Petrov, 2007; Sturgill et al., 2007; Vibranovski et al., 2009; Khodursky et al., 2020). We tested whether transcriptome turnover rate differs between X-linked and autosomal genes using the same approach as described above for transcription factors. We found that both classes of genes turn over at similar rates. The posterior distribution of PDMR overlaps zero and places little probability on differences greater than about 10% in either organ (Figure 10; top row). For X-linked genes, the reference distribution of mean PDMR suggests little surprise at seeing this pattern (Figure 10; bottom row) when comparing gene partitions that are random with respect to X-linkage. In summary, the two male reproductive organs show little evidence of either slower or faster evolutionary turnover in the expression of X-chromosomal genes compared to other single-copy gene families.

**Figure 10:**
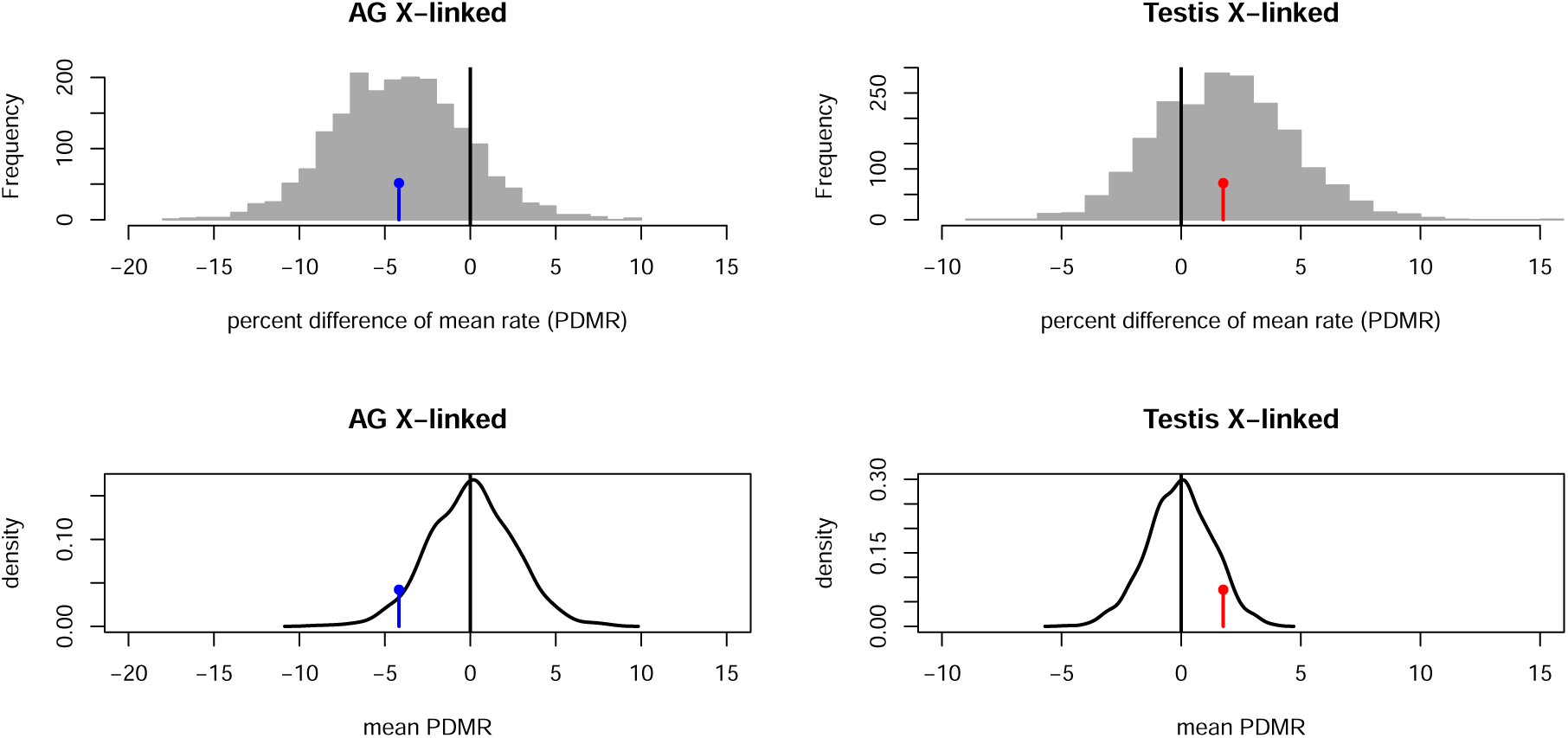
The posterior distributions of the percent difference of mean rate (PDMR) between X-linked and autosomal genes (top row). For example, the histogram in the top left shows the mean rate of the X-linked genes minus the mean rate of the autosomal genes divided by the mean autosomal rate computed for each MCMC sample (n=2,000). The colored lines topped with a circle indicate the mean of the posterior distribution of PDRM. The bottom row shows reference distributions of mean PDMR (M-PDMR) of 1,000 random partitions of all genes into two groups equivalent in size to the X-chromosome/autosome partition (black lines). The mean PDMRs of the X-linked genes (colored lines) are well within the control distributions.

### Ancestral Transcriptomes of Testes and Accessory Glands

We used our phylogenetic model to estimate the posterior probability that each single-copy gene was expressed in the testes and AGs of the most recent common ancestor of the 11 species in our analysis (i.e., the most recent common ancestor of the *melanogaster* species group). Our model and data imply, at a probability cutoff of 0.95, that 63% of the single-copy genes were expressed in the ancestral accessory gland, and 75% were expressed in the ancestral testis. We also estimated the posterior probability that each gene is actively expressed in *D. melanogaster* but was not expressed in the most recent common ancestor (i.e., that it gained expression in the testis or AG in the *D. melanogaster* lineage over the last *∼*25 MY) by analyzing the joint posterior distribution of the expression state at the root node of the tree and at the *D. melanogaster* tip (Data file S3). This analysis implies that at a probability *>* 0.95, *D. melanogaster* expresses 23 single-copy genes in the AG that were not expressed in that organ in the most recent common ancestor of the *melanogaster* species group, while 112 such genes are expressed in the *D. melanogaster* testis.

These results raise several questions, in particular – how big are the changes in transcript abundance associated with the gain of active expression? Do recently activated genes remain expressed at levels barely above our probability cut-offs, or does their expression approach the levels typical of other genes expressed in that tissue? To address this question, we compared the expression levels of newly active and ancestrally active genes in each species in both organs (Figure 11 A,B). Newly active genes were defined as those that have *>* 0.95 probability of being ON in the focal species and *>* 0.95 probability of being OFF in the most recent common ancestor of the 11 species in our study, and ancestrally active genes as those with *>* 0.95 probability of being ON both in the focal species and in the common ancestor. We estimated the posterior distribution of the difference in log TPM between these two groups using the R package BEST v 0.5.4 (Kruschke, 2013) setting default vague priors (Figure 11 C). Resulting inferences imply that the two organs differ in the expression levels of newly active genes. In testes, such genes have lower expression compared to ancestrally active genes, while in the AG both classes of genes are expressed at roughly similar levels (Figure 11). Thus, in at least some tissues newly activated genes approach typical expression levels. In fact, some recently activated genes are expressed at high TPM (Figure 3, Figure 7 D).

**Figure 11:**
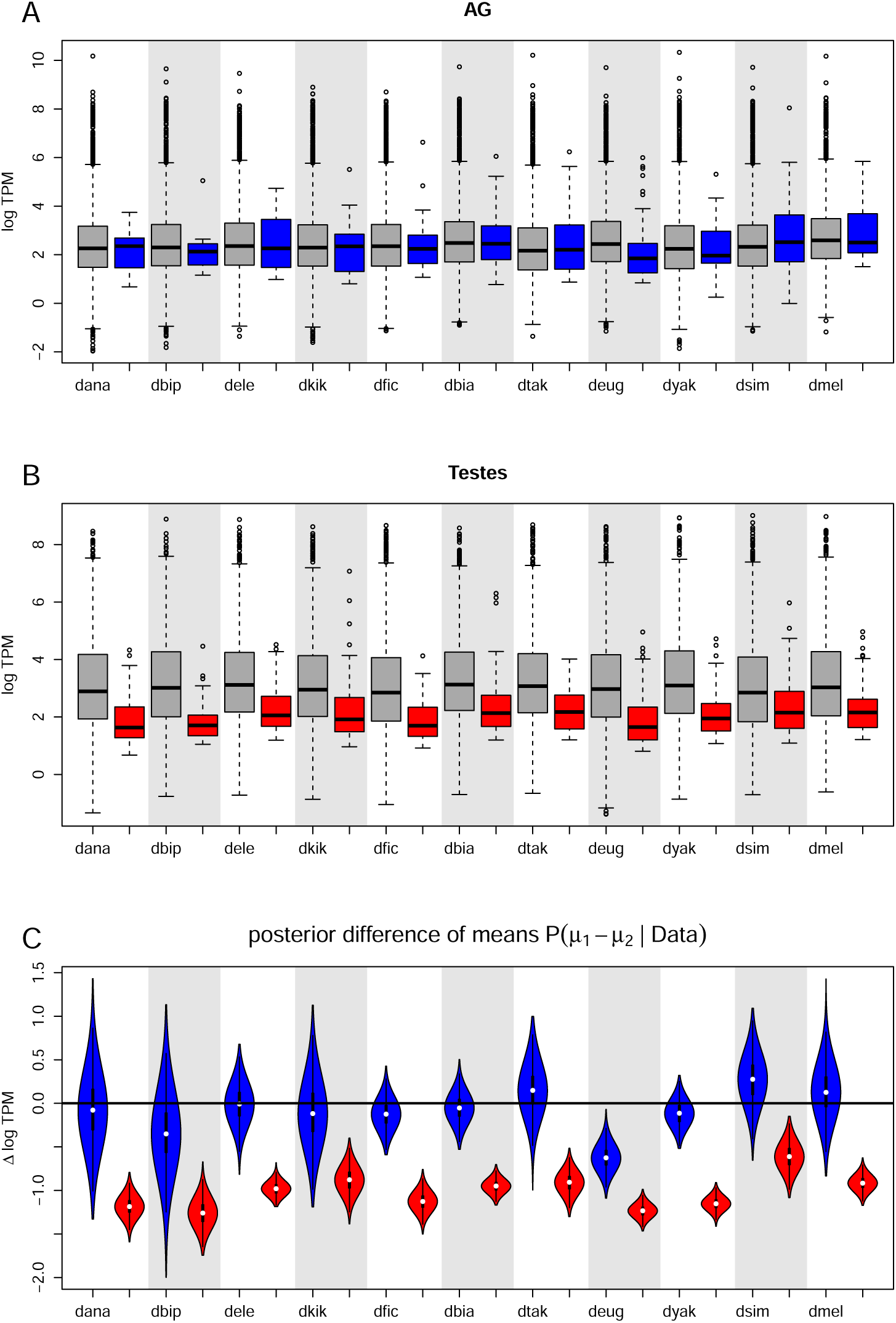
Expression of recently activated genes compared to genes with conserved expression. A,B) “Recently activated” is defined as having *>* 0.95 probability of being OFF in the common ancestor and ON in species X (A; blue boxes for AG, B; red for testes). “conserved expression” is defined as having *>* 0.95 probability of being ON in the common ancestor and ON in species X (gray boxes). C) The posterior distribution of the difference in log TPM between the two groups (*µ*_1_ is the mean of the recently activated group and *µ*_2_ is the mean of the conserved expression group).

## Discussion

In this report, we explored the feasibility of using discretized RNA-seq data to study the evolutionary turnover of transcriptomes—that is, the gain and loss of tissue-specific expression by conserved genes. We created an analysis pipeline that fits discrete trait evolution models to comparative RNA-seq datasets. This pipeline translates continuous RNA-seq data to discrete trait data using a Bayesian method for expression state inference (*zigzag*; Thompson et al., 2020). To investigate the co-evolution of transcriptomes across multiple organs, we defined a phylogenetic model of correlated gene expression evolution in two or more organs. We demonstrated the power of this research pipeline by examining two rapidly evolving organs in the male reproductive system, namely the accessory glands and the testes.

We first characterized attributes of the two organ transcriptomes in each species individually. We estimated that testes express a greater proportion of the genome compared to accessory glands, with a proportion of actively expressed genes falling in the 65–75% range in the testis and 46–64% in the AG across the 11 species (95% HPD). The level of background expression noise (i.e., the fraction of the transcriptome that does not correspond to actively expressed genes) is similar between the two organs and is below 1% in all species. In other words, more than 99% of protein-coding transcripts in cells appear to be from actively expressed genes.

With our model of correlated transcriptome evolution, we inferred turnover rates and historical expression states of 8,660 single-copy genes. This inference suggests that the transcriptomes of the two male reproductive organs evolve at similar rates but follow distinct evolutionary paths. The accessory glands and testes show bursts of rapid transcriptome turnover at different times and in different parts of the tree. Among genes, there is greater variation in turnover rates in the AG than in the testis, indicating a lower baseline rate of turnover in the AG with a subset of genes showing higher rates of evolution.

We explored the evolutionary patterns of two important classes of genes, transcription factors and X-linked genes, to test whether they follow distinct evolutionary dynamics compared to the rest of the genome. Transcription factor expression appears to evolve faster than the non-TF single-copy genes in the testis, but much slower in the AG. It is interesting to note that our estimates of the overall transcriptome turnover rates are similar for the two organs. Given that changes in TF expression are expected to have a large impact on the transcriptome, this is a curious result. One possibility is that the gain and loss of TF expression may modulate the expression levels of genes that are already expressed in the organ, rather than changing the expression states of genes from off to on or vice versa. Alternatively, downstream expression changes that result from TF turnover could be more pronounced in non-single-copy gene families that undergo duplication and deletion more frequently. Finally, we note that the posterior distributions of the mean AG and testis turnover rate are both fairly wide, and thus moderate to large differences in rates are plausible. Analysis among more species could help provide more precise estimates.

We find that X-linked genes turn over at rates not significantly different from autosomal genes. Several studies report that genes with testis-biased and AG-biased expression tend to be under-represented on the X chromosome in *D. melanogaster* and other *Drosophila* species, suggesting that X-linked genes with male-specific functions may be disfavored by selection (Parisi et al., 2004; Mikhaylova and Nurminsky, 2011; Assis et al., 2012; Meisel et al., 2012). Our estimates of expression state indicate X-linked genes are actually more likely to be expressed in the male reproductive organs than the average gene. We do not observe an evolutionary trend toward regulatory silencing of X-linked genes in the testis or AG, suggesting that such silencing is unlikely to play a central role in the demasculinization of the X chromosome.

### Challenges and Pitfalls

Different dimensions of the dataset are informative on different model parameters. An obvious question is how this type of analysis is influenced by the number of species and the number of transcriptome samples per species. The first step in our analysis pipeline involves inferring the expression states of genes from replicated continuous (TPM) data. Four samples per species appear to be sufficient for *zigzag*, as we find that increasing the number of samples further provides only marginally more precise estimates (Thompson et al., 2020). As for the number of taxa, we found that with just 11 species, two organs, and 8,660 genes, we were able to detect differences in the mode and tempo of transcriptome turnover both between tree branches and between organs. However, many model parameters of interest, such as the correlation of turnover rates between the two organs, have posterior distributions that are quite wide. This is also true of the posterior distributions for the turnover rates of individual genes. Datasets containing more species should better resolve these parameters as well as many other important questions relating to transcriptome evolution while allowing for richer and biologically realistic models of regulatory evolution. In summary, gene-level parameter estimates should benefit primarily from more species, while transcriptome-level parameters should benefit from more species as well as from including more genes in the analysis.

The first point of failure in any comparative transcriptome study is dissection of biological samples. Because “comparative” assumes homology, it is crucial that RNA samples are isolated from truly homologous tissues. This is not a trivial problem if the organ system under study changes structure frequently and/or is not defined in a consistent way among species. It is easy to imagine scenarios where it is difficult to distinguish the evolutionary co-option of new genes in a transcriptome from contamination by other tissues and cells. We took many precautions with the accessory glands and testis at several steps in our analysis pipeline to detect and mitigate potential artifacts emanating from dissection error. Our analysis of individual gene expression patterns suggests that cross-tissue contamination is not a major source of artifacts in this study. Nevertheless, it is important to consider this potential source of erroneous inference in all comparative transcriptome analyses, whether qualitative or quantitative.

The next set of challenges emerge from how expression is defined and quantified. Similar to comparative analyses that treat gene expression as a continuous character, evolutionary inference of discrete expression states is fraught with potential artifacts emanating from the data processing pipeline (Khaitovich et al., 2005; Harrison et al., 2012; Romero et al., 2012; Rohlfs et al., 2014; Dimayacyac et al., 2023; Pal et al., 2023). The most obvious factors to influence results would be genome assembly and annotation quality. If some species have few reads mapping to an ortholog because the gene was not correctly assembled or given an incorrect annotation, that could obviously influence the inferred expression state and comparative patterns for that gene.

An essential assumption for conducting a macroevolutionary analysis is that expression states of genes don’t vary among populations in each species, which may not be the case for some genes (Cridland et al., 2020). Our approach to this problem was to sample two distinct populations in each species and then run a leave-one-library-out *zigzag* analysis to see which genes are sensitive to the exclusion of individual libraries (see Methods).

Our use of a new method for classifying gene expression states, *zigzag*, requires careful model checking, sensitivity tests, and validation; indeed, posterior predictive checks and cross-validations are essential for any study seeking to combine numerous complex models in an inference pipeline. Our sensitivity and posterior predictive checks and cross-validation experiments suggest some robustness. However, it can be difficult to know how robust the inferences are because when it comes to expression states of genes, ground truths are themselves inferences under models. The fact that we observe an 88% probability of agreement between evolutionary predictions of the expression states of holdout genes and the *zigzag* predictions of their expression states (Supplemental Text S1) appears to be a good indication that our analysis is providing a fair approximation of the expression states and evolutionary processes behind the comparative expression patterns in our data. However, interpreting this result requires care, and we encourage others to critically explore potential weaknesses that may exist in this approach. As in any comparative study, our results depend on the accuracy with which species relationships are inferred. We treat the species tree and root age as known, when in fact these are estimates and thus have some unmodeled level of uncertainty around them. We have high confidence in the tree topology, since it is similar to the phylogeny based on hundreds of loci distributed across the entire genome (Suvorov et al., 2021). On the other hand, the estimates of node ages in *Drosophila* are notoriously uncertain due to the dearth of fossil calibration points (Suvorov et al., 2021). Including this uncertainty may decrease the precision of posterior estimates of branch rates.

Another critical assumption in our methods (as in other analyses of gene expression evolution) is that each gene is evolving independently. This is a convenient simplifying assumption of the phylogenetic inference model. However, it is unknown how much bias emanates from this simplification for transcriptome evolution, where this is obviously not true given the regulatory interactions among genes. If the expression state of the average gene is governed by a complex interaction of numerous genes, then this assumption of independence may be a reasonable approximation. However, if many genes are highly sensitive to the state of a small number of genes, this assumption may result in misleading inferences.

### Future Directions

The first step toward identifying causal mechanisms is to find patterns of correlation. While quantitative changes in gene expression are both highly prevalent and affect adaptive phenotypes, qualitative gain or loss of gene expression can, like gene deletion and duplication, have especially profound consequences for organ structure and function (Gompel et al., 2005; Rebeiz et al., 2009; Chan et al., 2010; Fraser et al., 2010; Arnoult et al., 2013; Loehlin et al., 2019; Kowalczyk et al., 2022; Marand et al., 2023). Potential cases of association between genetic and phenotypic change can be identified by phylogenetic analysis: if a gene was activated or inactivated on the branch where a new phenotype evolved, that gene may be part of a tissuespecific regulatory pathway mediating the evolutionary change. In this study, we took a step toward a systematic comparative analysis of gene expression states by developing a method for quantifying the rate of transcriptome turnover on phylogenies. In the future, we can build upon this foundation to map the transitions between active and inactive gene expression states to specific branches of species trees. This approach is likely to provide new insights into the regulatory mechanisms driving phenotypic evolution.

This study focuses only on single-copy gene families, which comprise slightly over half of the genome and may not be representative of the transcriptome as a whole. Gene families with unstable sizes (those undergoing frequent gene duplications and losses) may also be subject to faster regulatory turnover (Lynch and Force, 2000; Lynch and Conery, 2000; Makino and McLysaght, 2010; Thompson et al., 2016; Ebadi et al., 2023). To fully understand how organ function evolves, it is essential to characterize the role of genome evolution in transcriptome turnover. This involves developing models that integrate evolutionary processes that add and remove genes from the genome with regulatory processes that turn genes on and off. Though challenging, solving this problem will uncover complex relationships, such as those between gene duplication and expression changes or regulatory inactivation and gene loss. As genome annotations improve, expression estimation becomes more precise, and new evolutionary models are developed, we anticipate many novel and intriguing questions will arise from this research.

## Methods

### RNA Sequencing and Expression Estimation

*D. melanogaster* testis expression estimates were obtained from Thompson et al. (2020). All other RNA-seq libraries from testes and accessory glands were created and analyzed using the same methods as in Thompson et al. (2020). In brief, mRNA from each organ was extracted from approximately 25 mixed-stage adult male flies. At least two biological replicates each from the genome reference strain (used for genome assembly and annotation) and from a second, non-reference strain were obtained. Sequencing was performed on either a HiSeq2500, 3000, or 4000 (see the supplemental methods S2.3.2 of Thompson et al., 2020). We mapped paired-end reads to the genome annotations (see Table 3 below) using STAR v. 2.5.3 (Dobin et al., 2013). We assembled reference-only transcripts using Stringtie v. 1.3.4 (Pertea et al., 2015) and estimated transcript abundance in transcripts per million (TPM). Suppl. Table S1 provides more details about samples.

**Table 3:** Genome and annotation information.

| Species | Genome release |
| --- | --- |
| *D. melanogaster* | r6.12 |
| *D. simulans* | r2.02 |
| *D. yakuba* | r1.05 |
| *D. eugracilis* | Deug_2.0 |
| *D. biarmipes* | Dbia_2.0 |
| *D. takahashii* | Dtak_2.0 |
| *D. ficusphila* | Dfic_2.0 |
| *D. kikkawai* | Dkik_2.0 |
| *D. elegans* | Dele_2.0 |
| *D. ananassae* | r1.05 |
| *D. bipectinata* | Dbip_2.0 |

### Expression State Inference

We used the R package *zigzag* (Thompson et al., 2020) v.1.0.0 to infer gene expression states from TPM values from multiple biological replicates (libraries) of each tissue and species. *zigzag* assumes that inactive and active genes are drawn from distinct distributions in a mixture model, which are approximately normal and overlap to varying degrees depending on the tissue and species. The model also assumes that individual libraries (biological replicates) are noisy samples from the combined inactive and active distributions of the mixture model. Importantly, *zigzag* assumes that the latent true expression state of each gene is shared among all biological replicates. Increasing numbers of libraries therefore increases the certainty about the expression states of genes. Table 4 shows the prior settings assumed for the accessory glands and the testis.

**Table 4:** Summary of zigzag prior distributions.

| Parameter | Distribution Type | Distribution Parameters | Description |
| --- | --- | --- | --- |
| $s_0$ | Normal | $\mu = -1, \sigma = 2$ | baseline variance in expression |
| $-s_1$ | Exponential | $\lambda = 2$ | influence of expression level on variance |
| $\tau$ | Exponential | $\lambda = 1$ | variation in gene-specific variance |
| $\alpha_r$ | Exponential | $\lambda = 1/10$ | Proportional to library dropout rate |
| $\omega_a$ | Beta | $\alpha = 2, \beta = 2$ | Proportion active |
| $\omega_a^k$ | Beta | $\alpha = 1, \beta = 1$ | Proportional within active subcomponent $k$ |
| $\rho$ | Beta | $\alpha = 1, \beta = 1$ | Dropout parameter |
| $\mu_{a1} - 1$ | Exponential | $\lambda = 1/3$ | Mean of active component $a1$ |
| $\mu_{a2} - 4$ | Exponential | $\lambda = 1/3$ | Mean of active component $a2$ |
| $\mu_{a3} - 6$ | Exponential | $\lambda = 1/3$ | Mean of active component $a3$ (testis only) |
| $-\mu_i$ | Exponential | $\lambda = 1/3$ | Mean of inactive component |
| $\sigma^2$ | logUniform | $\log(\min) = \log(0.01),$<br>$\log(\max) = \log(5)$ | Variance parameter shared by all mixture components |

We updated *zigzag* (v1.0.0) so that all mixture components could share a single variance parameter. This significantly improves MCMC convergence and efficiency with apparently little impact on inference for our samples. Additionally, the previous default model (v0.1.0) allowing the variances to vary between inactive and active components puts positive prior probability on unrealistic models where, for example, if the variance of the inactive component is much higher than the active component(s), genes with very high expression could be assigned to the inactive component. Two independent MCMC chains were run for 25,000 cycles, sampled every 5 cycles, and compared for convergence by measuring PSRF (Brooks and Gelman, 1998). A PSRF close to 1 and less than 1.2 is considered a good convergence. All parameters for both organs had PSRF *<* 1.2.

#### Model Adequacy and Validation

We used posterior predictive simulation to select a set of hyperparameters and number of mixture components that do not inappropriately influence the posterior. We focused on re-simulating library expression distributions and comparing them to the data distributions through plots to select appropriate mixture distributions and prior thresholds (Data File S1, upper-level plots). We found that two active mixture components for the accessory glands and three for the testis worked well. We visiually analyzed posterior predictive plots output by *zigzag* and found that the simulated library distributions from testes matched very closely with the data, while the accessory glands showed a much weaker match in many of the species (Data File S1 lower-level (library-specific) plots).

We checked the sensitivity of all genes to this mismatch to see which genes may violate the assumption of the same expression state in all replicate samples by performing leave-onelibrary-out analysis. Each library was removed from the set and expression state probabilities were reestimated. Thus, for a set of 4 replicate libraries, we obtained 4 expression state probabilities for each gene. Any gene where the difference between the maximum and minimum probability of active expression was greater than 0.25 was treated as unknown for downstream analyses. The accessory glands contained many more such genes (60–755) than the testis (29– 122). This could be due to the greater challenge of dissecting the AG cleanly and the presence of contaminants from neighboring tissues, which we expect to be more variable among replicates.

### Orthology table and Species tree inference

An orthology table was estimated using OrthoFinder v. 2.4.0 with default settings (Emms and Kelly, 2019). This produced 15,322 gene families including 8,660 conserved single-copy families (i.e., one gene per family in each of the 11 species), constituting 56% of all genes. A comprehensive list of *D. melanogaster* transcription factors was downloaded from FlyMine.org. Of the 757 TFs, 75% were in single-copy gene families.

The phylogenetic tree of the 11 *Drosophila* species was inferred using RevBayes (Höhna et al., 2016). The goal was to create a species tree with accurate relative node ages (branch lengths in time units rather than expected number of nucleotide substitutions) that could then be treated as fixed in the phylogenetic analysis of gene expression states. To construct this tree, we used 12 conserved single-copy genes from the dataset that was used to infer a *Drosophila* chronogram by (Turelli et al., 2018). In each species, we used the isoform with the longest protein sequence of the following genes: *Ald1*, *bcd*, *Eno*, *esc*, *Glyp*, *Glys*, *ninaE*, *Pepck1*, *Pgi*, *pic*, *Tpi*, and *Taldo*. Mafft v.7.471 on auto settings was used to align each set of 11 homologous proteins, followed by pal2nal v.14 (Suyama et al., 2006) to convert protein alignments into codon alignments. All codon alignments were then partitioned by codon position for phylogenetic analysis, yielding a dataset with 36 partitions.

All phylogenetic analyses were conducted in RevBayes v. 1.1.0. Posterior samples of relative chronograms were simulated under three clock models: a global clock (one rate for the whole tree), uncorrelated relaxed clock (UCLN) with one rate for each branch which is uncorrelated with neighboring branches, and an autocorrelated relaxed clock model (ACLN) where neighboring branch rates were correlated (Drummond et al., 2006). All partitions were assumed to evolve at the same clock rate on each branch, with root age set to 1. All three models assumed the same substitution model for each individual partition (3 partitions per gene for 12 genes): GTR + G + I, with G having four rates that are distributed according to a discretized Gamma distribution with mean set to 1 and I being the proportion of invariant sites.

For the tree prior in all analyses, we assumed the sampled birth-death model of (Yang and Rannala, 1997). For the strict and uncorrelated analyses, the number of species in the clade that spans the sampled species was assumed to be around 220–240, which is slightly larger than the number of described species in the *melanogaster* species group (spanning the clade from *D. ananassae* to *D. melanogaster*). We averaged over this uncertainty by specifying an informative prior of Gamma (529, 2.3). MCMCs were run for 75,000 generations with 10% burnin. ESS was checked so that all parameters had ESS *>* 150. Multiple independent MCMCs converged on identical posterior distributions for parameters and trees in each model.

We inferred the topology of the tree under an uncorrelated relaxed clock model which supported the phylogeny used with at least 0.92 posterior probability at each branch except for the placement of *D. kikkawai*, which was equally likely placed with the (*D. anannassae* + *D. bipectinata*) clade as with the other clade at the first post-root split. To estimate branch lengths under a more realistic autocorrelated relaxed clock model, we selected the topology that placed *D. kikkawai* basal to the other 9 species (Suvorov et al., 2021) and treated that as fixed. The tree topology matched almost exactly the topology that was recovered in a larger analysis that included *>*100 species and several hundred conserved dipteran BUSCO genes (Suvorov et al., 2021). The one difference was the locations of *D. elegans* and *D. ficusphila*, which are reversed in our UCLN analysis with posterior probability = 0.92 on the branch separating the two. The maximum a posteriori (MAP) tree with UCLN MAP estimate of topology and ACLN MAP estimate of branch lengths was used for subsequent analyses of transcriptome evolution.

### Transcriptome evolution inference

We modeled the evolution of gene expression states as a continuous-time Markov chain (CTMC) with two states, *{*OFF, ON*}*. The transition rates between these states vary across genes and branches of the phylogenetic tree. Additionally, the branch-specific rates are correlated between the two organs.

#### Relative rates model

We assume that the process is at stationarity over the time scale of the whole tree, with genes at the root being active (ON) in organ *i* with frequency *π_i_*, where *i* indexes the organs *{*AG, Testis*}*. The relative transition rate matrix, **Q***_i_*, is parameterized by *π_i_* such that the mean transition rate across all genes is one (Yang, 1994). Specifically,

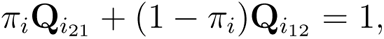

where 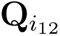 is proportional to the rate at which genes are activated (turned ON), and 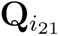 is proportional to the rate at which genes are deactivated (turned OFF). The relative rate matrix is therefore:

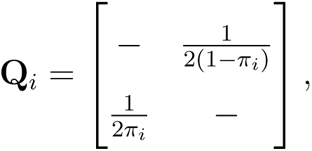

where dashes along the diagonal are set so that rows sum to zero.

#### Among-gene rate heterogeneity model

To model rate variability among genes, we specified a scale factor, *a_i,g_*, to scale the relative rate matrix, **Q***_i_* for each gene, *g*. This variable is drawn from a discretized gamma distribution with mean fixed to 1 (Yang, 1994):

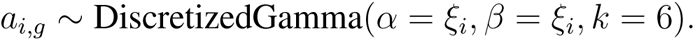

Here, the distribution is a mixture model where a gamma distribution, with mean *α/β* = 1 and variance *α/β*^2^ = 1*/ξ_i_*, and is divided into *k* = 6 intervals with equal probability. The mixture weights are thus 1/6. The rate category at each interval is the mean rate for that interval. *ξ_i_* is the precision of the distribution.

#### Among-branch rate heterogeneity model

In our model of turnover rate heterogeneity across branches of the tree, the evolutionary rates for both organs on each branch follow a bivariate log-normal distribution. The bivariate log-transformed rates, log **r***_b_*, for the two organs on branch *b* are given by:

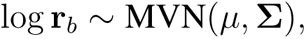

The transition rate matrix for gene, *g* in organ, *i* on branch, *b* is thus equal to

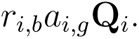

We summarize the prior distributions for the paramaters of the above model in Table 5 below:

**Table 5:** Summary of the probabilistic model parameters and their distributions or definitions. The LKJ distribution is the Lewandowski-Kurowicka-Joe distribution (Lewandowski et al., 2009).

| Parameter | Distribution Type | Distribution Parameters | Description |
| --- | --- | --- | --- |
| $\pi_i$ | Beta | $\alpha = \beta = 1$ | Frequency genes at the root are active |
| $\xi_i$ | LogNormal | $\log\text{mean} = \ln(5),$<br>$\log\text{sd} = 0.587$ | Precision of the discretized gamma distribution of $a_{i,g}$ |
| $\phi$ | Uniform | $\min = -10,$<br>$\max = 1$ | Log mean branch scale parameter |
| $\alpha_i$ | Beta | $\alpha = 2, \beta = 2$ | Proportional mean rates of 2 organs |
| $\mu_i$ | Defined | $\phi + \ln(2\alpha_i)$ | Log mean branch rate of organ $i$ |
| $\theta$ | LogUniform | $\min = 0.0001,$<br>$\max = 5$ | Branch rate variance scale parameter |
| $\beta_i$ | Beta | $\alpha = 10, \beta = 10$ | Proportional branch rate variance |
| $\sigma_i^2$ | Defined | $\theta + 2\beta_i$ | Variance of branch rates of organ $i$ |
| $C$ | LKJ | $\eta = 1$ | Correlation matrix |
| $S$ | Defined | $\begin{pmatrix} \sigma_1 & 0 \\ 0 & \sigma_2 \end{pmatrix}$ | Organ branch rate standard deviation |
| $\Sigma$ | Defined | $SCS$ | Covariance matrix |

#### Markov Chain Monte Carlo (MCMC)

We sampled from the model posterior using RevBayes (Hohna et al. 2016). After an initial burnin period, we ran the MCMC for 10,000 generations sampling every 5 generations. We confirmed that two independent runs converged by measuring the PSRF for all parameters and checking that it was less than 1.2. We also checked that ESS was at least 200.

### GO analysis

GO analyses were performed in R v4.1.1 using clusterProfiler (v4.0.5; Yu et al. 2012). Outputs were plotted using clusterProfiler, ggplot2 (v3.4.4), and ggVennDiagram (v1.5.2; Gao et al. 2021). Ontogeny was limited to ‘Biological Process’ and a Benjamini-Hochberg procedure was used to correct for the false discovery rate. A q-value cutoff of 0.05 was used. The org.Dm.eg.db (v3.13.0) package was used for gene annotation data.

### BEST analysis

The prior distributions and data distribution assumed in *BEST* (Kruschke, 2013) are listed below in Table 6.

**Table 6:**
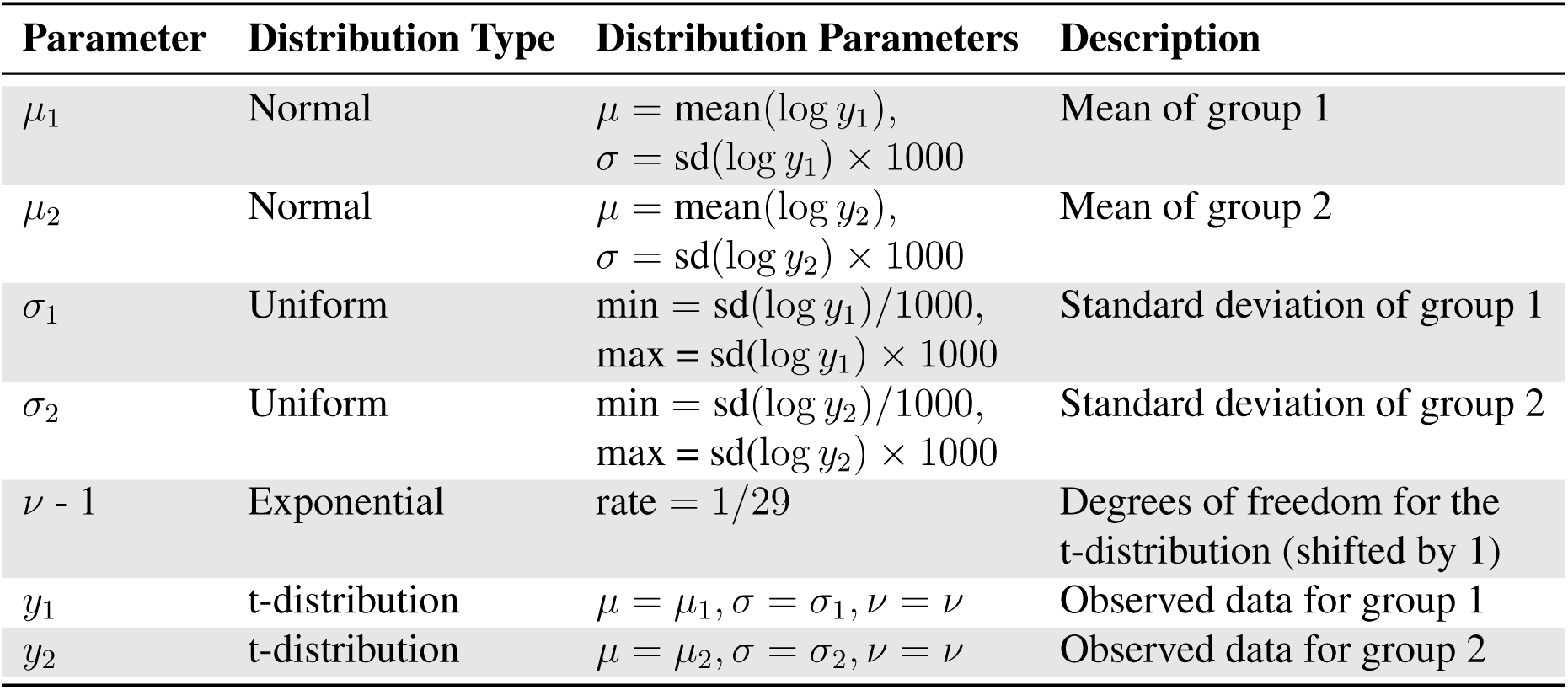
Summary of the probabilistic model parameters and distributions with specified priors.

| Parameter | Distribution Type | Distribution Parameters | Description |
| --- | --- | --- | --- |
| $\mu_1$ | Normal | $\mu = \text{mean}(\log y_1),$<br>$\sigma = \text{sd}(\log y_1) \times 1000$ | Mean of group 1 |
| $\mu_2$ | Normal | $\mu = \text{mean}(\log y_2),$<br>$\sigma = \text{sd}(\log y_2) \times 1000$ | Mean of group 2 |
| $\sigma_1$ | Uniform | $\text{min} = \text{sd}(\log y_1)/1000,$<br>$\text{max} = \text{sd}(\log y_1) \times 1000$ | Standard deviation of group 1 |
| $\sigma_2$ | Uniform | $\text{min} = \text{sd}(\log y_2)/1000,$<br>$\text{max} = \text{sd}(\log y_2) \times 1000$ | Standard deviation of group 2 |
| $\nu - 1$ | Exponential | $\text{rate} = 1/29$ | Degrees of freedom for the t-distribution (shifted by 1) |
| $y_1$ | t-distribution | $\mu = \mu_1, \sigma = \sigma_1, \nu = \nu$ | Observed data for group 1 |
| $y_2$ | t-distribution | $\mu = \mu_2, \sigma = \sigma_2, \nu = \nu$ | Observed data for group 2 |

## Acknowledgements

This work was supported by NIH grants 5F32GM125107-02 to A.T. and R35 GM122592 to A.K. and a long-term fellowship from the Human Frontier Science Program Organization (LT000123/2020-L to B.R.H.)

## Author Contributions

## Data Availability

## Conflicts of Interest

## Notes

### Competing Interest Statement

The authors have declared no competing interest.

## References

L. Arnoult, K. F. Y. Su, D. Manoel, C. Minervino, J. Magriña, N. Gompel, and B. Prud’homme. Emergence and diversification of fly pigmentation through evolution of a gene regulatory module. Science (New York, N.Y.), 339(6126):1423–1426, Mar. 2013. ISSN 1095-9203. doi: 10.1126/science.1233749.

C. G. Artieri and H. B. Fraser. Evolution at two levels of gene expression in yeast. Genome Research, 24(3):411–421, Mar. 2014. ISSN 1088-9051, 1549-5469. doi: 10.1101/gr.165522.113.

R. Assis, Q. Zhou, and D. Bachtrog. Sex-Biased Transcriptome Evolution in Drosophila. Genome Biology and Evolution, 4(11):1189–1200, Jan. 2012. ISSN 1759-6653. doi: 10.1093/gbe/evs093.

D. J. Begun, P. Whitley, B. L. Todd, H. M. Waldrip-Dail, and A. G. Clark. Molecular population genetics of male accessory gland proteins in Drosophila. Genetics, 156(4):1879–1888, Dec. 2000. ISSN 0016-6731. doi: 10.1093/genetics/156.4.1879.

D. Brawand, M. Soumillon, A. Necsulea, P. Julien, G. Csárdi, P. Harrigan, M. Weier, A. Liechti, A. Aximu-Petri, M. Kircher, F. W. Albert, U. Zeller, P. Khaitovich, F. Grützner, S. Bergmann, R. Nielsen, S. Pääbo, and H. Kaessmann. The evolution of gene expression levels in mammalian organs. Nature, 478(7369):343–348, Oct. 2011. ISSN 1476-4687. doi: 10.1038/na-ture10532.

S. P. Brooks and A. Gelman. General Methods for Monitoring Convergence of Iterative Simulations. Journal of Computational and Graphical Statistics, 7(4):434–455, Dec. 1998. ISSN 1061-8600, 1537-2715. doi: 10.1080/10618600.1998.10474787.

M. Cardoso-Moreira, J. Halbert, D. Valloton, B. Velten, C. Chen, Y. Shao, A. Liechti, K. Ascenção, C. Rummel, S. Ovchinnikova, P. V. Mazin, I. Xenarios, K. Harshman, M. Mort, D. N. Cooper, C. Sandi, M. J. Soares, P. G. Ferreira, S. Afonso, M. Carneiro, J. M. A. Turner, J. L. VandeBerg, A. Fallahshahroudi, P. Jensen, R. Behr, S. Lisgo, S. Lindsay, P. Khaitovich, W. Huber, J. Baker, S. Anders, Y. E. Zhang, and H. Kaessmann. Gene expression across mammalian organ development. Nature, 571(7766):505–509, July 2019. ISSN 0028-0836, 1476-4687. doi: 10.1038/s41586-019-1338-5.

S. B. Carroll. Evo-devo and an expanding evolutionary synthesis: A genetic theory of morphological evolution. Cell, 134(1):25–36, July 2008. ISSN 1097-4172. doi: 10.1016/j.cell.2008.06.030.

Y. F. Chan, M. E. Marks, F. C. Jones, G. Villarreal, M. D. Shapiro, S. D. Brady, A. M. Southwick, D. M. Absher, J. Grimwood, J. Schmutz, R. M. Myers, D. Petrov, B. Jónsson, D. Schluter, M. A. Bell, and D. M. Kingsley. Adaptive evolution of pelvic reduction in sticklebacks by recurrent deletion of a Pitx1 enhancer. Science (New York, N.Y.), 327(5963): 302–305, Jan. 2010. ISSN 1095-9203. doi: 10.1126/science.1182213.

C.-M. Chen, Y.-L. Lu, C.-P. Sio, G.-C. Wu, W.-S. Tzou, and T.-W. Pai. Gene Ontology based housekeeping gene selection for RNA-seq normalization. Methods, 67(3):354–363, June 2014. ISSN 1046-2023. doi: 10.1016/j.ymeth.2014.01.019.

J. D. Coolon, C. J. McManus, K. R. Stevenson, B. R. Graveley, and P. J. Wittkopp. Tempo and mode of regulatory evolution in Drosophila. Genome Research, 24(5):797–808, May 2014. ISSN 1549-5469. doi: 10.1101/gr.163014.113.

S. S. F. Costa, M. Rosikiewicz, J. Roux, J. Wollbrett, F. B. Bastian, and M. Robinson-Rechavi. Robust inference of expression state in bulk and single-cell RNA-Seq using curated intergenic regions, Apr. 2022.

V. Courtier-Orgogozo, L. Arnoult, S. R. Prigent, S. Wiltgen, and A. Martin. Gephebase, a database of genotype-phenotype relationships for natural and domesticated variation in Eukaryotes. Nucleic Acids Research, 48(D1):D696–D703, Jan. 2020. ISSN 1362-4962. doi: 10.1093/nar/gkz796.

J. M. Cridland, A. C. Majane, H. K. Sheehy, and D. J. Begun. Polymorphism and Divergence of Novel Gene Expression Patterns in Drosophila melanogaster. Genetics, 216(1):79–93, Sept. 2020. ISSN 1943–2631. doi: 10.1534/genetics.120.303515.

H. J. M. de Jonge, R. S. N. Fehrmann, E. S. J. M. de Bont, R. M. W. Hofstra, F. Gerbens, W. A. Kamps, E. G. E. de Vries, A. G. J. van der Zee, G. J. te Meerman, and A. ter Elst. Evidence Based Selection of Housekeeping Genes. PLOS ONE, 2(9):e898, Sept. 2007. ISSN 1932-6203. doi: 10.1371/journal.pone.0000898.

M.-A. Dillies, A. Rau, J. Aubert, C. Hennequet-Antier, M. Jeanmougin, N. Servant, C. Keime, G. Marot, D. Castel, J. Estelle, G. Guernec, B. Jagla, L. Jouneau, D. Laloë, C. Le Gall, B. Schaëffer, S. Le Crom, M. Guedj, F. Jaffrézic, and French StatOmique Consortium. A comprehensive evaluation of normalization methods for Illumina high-throughput RNA sequencing data analysis. Briefings in Bioinformatics, 14(6):671–683, Nov. 2013. ISSN 1477-4054. doi: 10.1093/bib/bbs046.

J. R. Dimayacyac, S. Wu, D. Jiang, and M. Pennell. Evaluating the Performance of Widely Used Phylogenetic Models for Gene Expression Evolution. Genome Biology and Evolution, 15(12):evad211, Dec. 2023. ISSN 1759-6653. doi: 10.1093/gbe/evad211.

A. Dobin, C. A. Davis, F. Schlesinger, J. Drenkow, C. Zaleski, S. Jha, P. Batut, M. Chaisson, and T. R. Gingeras. STAR: Ultrafast universal RNA-seq aligner. Bioinformatics (Oxford, England), 29(1):15–21, Jan. 2013. ISSN 1367-4811. doi: 10.1093/bioinformatics/bts635.

E. B. Dopman and D. L. Hartl. A portrait of copy-number polymorphism in Drosophila melanogaster. Proceedings of the National Academy of Sciences, 104(50):19920–19925, Dec. 2007. doi: 10.1073/pnas.0709888104.

A. J. Drummond, S. Y. W. Ho, M. J. Phillips, and A. Rambaut. Relaxed Phylogenetics and Dating with Confidence. PLOS Biology, 4(5):e88, Mar. 2006. ISSN 1545-7885. doi: 10.1371/journal.pbio.0040088.

M. Ebadi, Q. Bafort, E. Mizrachi, P. Audenaert, P. Simoens, M. V. Montagu, D. Bonte, and Y. V. de Peer. The duplication of genomes and gene regulatory networks and its potential for evolutionary adaptation and survival, Apr. 2023.

D. M. Emms and S. Kelly. OrthoFinder: Phylogenetic orthology inference for comparative genomics. Genome Biology, 20(1), Dec. 2019. ISSN 1474-760X. doi: 10.1186/s13059-019-1832-y.

J. Ernst, P. Kheradpour, T. S. Mikkelsen, N. Shoresh, L. D. Ward, C. B. Epstein, X. Zhang, L. Wang, R. Issner, M. Coyne, M. Ku, T. Durham, M. Kellis, and B. E. Bernstein. Mapping and analysis of chromatin state dynamics in nine human cell types. Nature, 473(7345): 43–49, May 2011. ISSN 1476-4687. doi: 10.1038/nature09906.

H. B. Fraser, A. M. Moses, and E. E. Schadt. Evidence for widespread adaptive evolution of gene expression in budding yeast. Proceedings of the National Academy of Sciences of the United States of America, 107(7):2977–2982, Feb. 2010. ISSN 1091-6490. doi: 10.1073/pnas.0912245107.

C.-H. Gao, G. Yu, and P. Cai. ggVennDiagram: An Intuitive, Easy-to-Use, and Highly Customizable R Package to Generate Venn Diagram. Frontiers in Genetics, 12:706907, 2021. ISSN 1664-8021. doi: 10.3389/fgene.2021.706907.

W. J. Glassford, W. C. Johnson, N. R. Dall, S. J. Smith, Y. Liu, W. Boll, M. Noll, and M. Rebeiz. Co-option of an Ancestral Hox-Regulated Network Underlies a Recently Evolved Morphological Novelty. Developmental Cell, 34(5):520–531, Sept. 2015. ISSN 1878-1551. doi: 10.1016/j.devcel.2015.08.005.

N. Gompel, B. Prud’homme, P. J. Wittkopp, V. A. Kassner, and S. B. Carroll. Chance caught on the wing: Cis-regulatory evolution and the origin of pigment patterns in Drosophila. Nature, 433(7025):481–487, Feb. 2005. ISSN 1476-4687. doi: 10.1038/nature03235.

W. Haerty, S. Jagadeeshan, R. J. Kulathinal, A. Wong, K. Ravi Ram, L. K. Sirot, L. Levesque, C. G. Artieri, M. F. Wolfner, A. Civetta, and R. S. Singh. Evolution in the Fast Lane: Rapidly Evolving Sex-Related Genes in Drosophila. Genetics, 177(3):1321–1335, Nov. 2007. ISSN 1943-2631. doi: 10.1534/genetics.107.078865.

L. Ham, D. Schnoerr, R. D. Brackston, and M. P. H. Stumpf. Exactly solvable models of stochastic gene expression. The Journal of Chemical Physics, 152(14):144106, Apr. 2020. ISSN 0021-9606. doi: 10.1063/1.5143540.

A. Harlin-Cognato, E. A. Hoffman, and A. G. Jones. Gene cooption without duplication during the evolution of a male-pregnancy gene in pipefish. Proceedings of the National Academy of Sciences of the United States of America, 103(51):19407–19412, Dec. 2006. ISSN 0027-8424. doi: 10.1073/pnas.0603000103.

P. W. Harrison, A. E. Wright, and J. E. Mank. The evolution of gene expression and the transcriptome-phenotype relationship. Seminars in Cell & Developmental Biology, 23(2): 222–229, Apr. 2012. ISSN 1096-3634. doi: 10.1016/j.semcdb.2011.12.004.

T. Hart, H. Komori, S. LaMere, K. Podshivalova, and D. R. Salomon. Finding the active genes in deep RNA-seq gene expression studies. BMC Genomics, 14(1):778, 2013. ISSN 1471-2164. doi: 10.1186/1471-2164-14-778.

D. Hebenstreit, M. Fang, M. Gu, V. Charoensawan, A. van Oudenaarden, and S. A. Teichmann. RNA sequencing reveals two major classes of gene expression levels in metazoan cells. Molecular Systems Biology, 7(1):497, Jan. 2011. ISSN 1744-4292, 1744-4292. doi: 10.1038/msb.2011.28.

C. N. Henrichsen, N. Vinckenbosch, S. Zöllner, E. Chaignat, S. Pradervand, F. Schütz, M. Ruedi, H. Kaessmann, and A. Reymond. Segmental copy number variation shapes tissue transcriptomes. Nature Genetics, 41(4):424–429, Apr. 2009. ISSN 1546-1718. doi: 10.1038/ng.345.

H. E. Hoekstra and J. A. Coyne. The Locus of Evolution: Evo Devo and the Genetics of Adaptation. Evolution, 61(5):995–1016, May 2007. ISSN 1558-5646. doi: 10.1111/j.1558-5646.2007.00105.x.

S. Höhna, M. J. Landis, T. A. Heath, B. Boussau, N. Lartillot, B. R. Moore, J. P. Huelsenbeck, and F. Ronquist. RevBayes: Bayesian Phylogenetic Inference Using Graphical Models and an Interactive Model-Specification Language. Systematic Biology, 65(4):726–736, July 2016. ISSN 1063-5157, 1076-836X. doi: 10.1093/sysbio/syw021.

B. R. Hopkins, I. Sepil, M.-L. Thézénas, J. F. Craig, T. Miller, P. D. Charles, R. Fischer, B. M. Kessler, A. Bretman, T. Pizzari, and S. Wigby. Divergent allocation of sperm and the seminal proteome along a competition gradient in Drosophila melanogaster. Proceedings of the National Academy of Sciences of the United States of America, 116(36):17925–17933, Sept. 2019. ISSN 1091-6490. doi: 10.1073/pnas.1906149116.

R. Hrdlickova, M. Toloue, and B. Tian. RNA-Seq methods for transcriptome analysis. Wiley interdisciplinary reviews. RNA, 8(1), Jan. 2017. ISSN 1757-7012. doi: 10.1002/wrna.1364.

Y. Hu, C. Schmitt-Engel, J. Schwirz, N. Stroehlein, T. Richter, U. Majumdar, and G. Bucher. A morphological novelty evolved by co-option of a reduced gene regulatory network and gene recruitment in a beetle. *Proceedings*. Biological Sciences, 285(1885):20181373, Aug. 2018. ISSN 1471-2954. doi: 10.1098/rspb.2018.1373.

L. Huang, Z. Yuan, P. Liu, and T. Zhou. Effects of promoter leakage on dynamics of gene expression. BMC systems biology, 9:16, Mar. 2015. ISSN 1752-0509. doi: 10.1186/s12918-015-0157-z.

T. H. Jensen, A. Jacquier, and D. Libri. Dealing with Pervasive Transcription. Molecular Cell, 52(4):473–484, Nov. 2013. ISSN 1097-2765. doi: 10.1016/j.molcel.2013.10.032.

R. T. Kellenberger, U. Ponraj, B. Delahaie, R. Fattorini, J. Balk, S. Lopez-Gomollon, K. H. Müller, A. G. Ellis, and B. J. Glover. Multiple gene co-options underlie the rapid evolution of sexually deceptive flowers in Gorteria diffusa. Current Biology, 33(8):1502–1512.e8, Apr. 2023. ISSN 0960-9822. doi: 10.1016/j.cub.2023.03.003.

A. D. Kern, C. D. Jones, and D. J. Begun. Molecular population genetics of male accessory gland proteins in the Drosophila simulans complex. Genetics, 167(2):725–735, June 2004. ISSN 0016-6731. doi: 10.1534/genetics.103.020883.

P. Khaitovich, S. Pääbo, and G. Weiss. Toward a Neutral Evolutionary Model of Gene Expression. Genetics, 170(2):929–939, June 2005. ISSN 0016-6731, 1943-2631. doi: 10.1534/genetics.104.037135.

S. Khodursky, N. Svetec, S. M. Durkin, and L. Zhao. The evolution of sex-biased gene expression in the Drosophila brain. Genome Research, 30(6):874–884, June 2020. ISSN 1088-9051, 1549-5469. doi: 10.1101/gr.259069.119.

K. Kin, M. C. Nnamani, V. J. Lynch, E. Michaelides, and G. P. Wagner. Cell-type Phylogenetics and the Origin of Endometrial Stromal Cells. Cell Reports, 10(8):1398–1409, Mar. 2015. ISSN 2211-1247. doi: 10.1016/j.celrep.2015.01.062.

M. C. King and A. C. Wilson. Evolution at two levels in humans and chimpanzees. Science (New York, N.Y.), 188(4184):107–116, Apr. 1975. ISSN 0036-8075. doi: 10.1126/sci-ence.1090005.

A. Kowalczyk, M. Chikina, and N. Clark. Complementary evolution of coding and noncoding sequence underlies mammalian hairlessness. eLife, 11:e76911, Nov. 2022. ISSN 2050-084X. doi: 10.7554/eLife.76911.

J. K. Kruschke. Bayesian estimation supersedes the t test. Journal of Experimental Psychology: General, 142(2):573–603, 2013. ISSN 1939-2222. doi: 10.1037/a0029146.

O. Lenive, P. D. W. Kirk, and M. P. H. Stumpf. Inferring extrinsic noise from single-cell gene expression data using approximate Bayesian computation. BMC Systems Biology, 10(1):81, Aug. 2016. ISSN 1752-0509. doi: 10.1186/s12918-016-0324-x.

D. Lewandowski, D. Kurowicka, and H. Joe. Generating random correlation matrices based on vines and extended onion method. Journal of Multivariate Analysis, 100(9):1989–2001, Oct. 2009. ISSN 0047-259X. doi: 10.1016/j.jmva.2009.04.008.

J. J. Li, P. J. Bickel, and M. D. Biggin. System wide analyses have underestimated protein abundances and the importance of transcription in mammals. PeerJ, 2:e270, Feb. 2014. ISSN 2167-8359. doi: 10.7717/peerj.270.

D. W. Loehlin, J. R. Ames, K. Vaccaro, and S. B. Carroll. A major role for noncoding regulatory mutations in the evolution of enzyme activity. Proceedings of the National Academy of Sciences of the United States of America, 116(25):12383–12389, June 2019. ISSN 1091-6490. doi: 10.1073/pnas.1904071116.

M. Lynch and J. S. Conery. The Evolutionary Fate and Consequences of Duplicate Genes. Science, 290(5494):1151–1155, Nov. 2000. ISSN 0036-8075, 1095-9203. doi: 10.1126/sci-ence.290.5494.1151.

M. Lynch and A. Force. The probability of duplicate gene preservation by subfunctionalization. Genetics, 154(1):459–473, Jan. 2000. ISSN 0016-6731.

T. Makino and A. McLysaght. Ohnologs in the human genome are dosage balanced and frequently associated with disease. Proceedings of the National Academy of Sciences of the United States of America, 107(20):9270–9274, May 2010. ISSN 1091-6490. doi: 10.1073/pnas.0914697107.

F. Mantica, L. P. Iñiguez, Y. Marquez, J. Permanyer, A. Torres-Mendez, J. Cruz, X. Franch-Marro, F. Tulenko, D. Burguera, S. Bertrand, T. Doyle, M. Nouzova, P. D. Currie, F. G. Noriega, H. Escriva, M. I. Arnone, C. B. Albertin, K. R. Wotton, I. Almudi, D. Martin, and M. Irimia. Evolution of tissue-specific expression of ancestral genes across vertebrates and insects. Nature Ecology & Evolution, pages 1–14, Apr. 2024. ISSN 2397-334X. doi: 10.1038/s41559-024-02398-5.

A. P. Marand, A. L. Eveland, K. Kaufmann, and N. M. Springer. Cis-Regulatory Elements in Plant Development, Adaptation, and Evolution. Annual Review of Plant Biology, 74: 111–137, May 2023. ISSN 1545-2123. doi: 10.1146/annurev-arplant-070122-030236.

S. Marguerat and J. Bähler. RNA-seq: From technology to biology. Cellular and molecular life sciences: CMLS, 67(4):569–579, Feb. 2010. ISSN 1420-9071. doi: 10.1007/s00018-009-0180-6.

J. Martin-Diaz and S. C. Herrera. A stem cell activation state coupling spermatogenesis with social interactions in Drosophila males. Cell Reports, 43(8):114647, Aug. 2024. ISSN 2211-1247. doi: 10.1016/j.celrep.2024.114647.

E. O. Martinson, n. Mrinalini, Y. D. Kelkar, C.-H. Chang, and J. H. Werren. The Evolution of Venom by Co-option of Single-Copy Genes. Current biology: CB, 27(13):2007–2013.e8, July 2017. ISSN 1879-0445. doi: 10.1016/j.cub.2017.05.032.

C. D. Meiklejohn, J. Parsch, J. M. Ranz, and D. L. Hartl. Rapid evolution of male-biased gene expression in Drosophila. Proceedings of the National Academy of Sciences of the United States of America, 100(17):9894–9899, Aug. 2003. ISSN 0027-8424. doi: 10.1073/pnas.1630690100.

R. P. Meisel, J. H. Malone, and A. G. Clark. Disentangling the relationship between sex-biased gene expression and X-linkage. Genome Research, 22(7):1255–1265, July 2012. ISSN 1088-9051. doi: 10.1101/gr.132100.111.

K. Mika, M. Marinić, M. Singh, J. Muter, J. J. Brosens, and V. J. Lynch. Evolutionary transcriptomics implicates new genes and pathways in human pregnancy and adverse pregnancy outcomes. eLife, 10:e69584, Oct. 2021. ISSN 2050-084X. doi: 10.7554/eLife.69584.

K. Mika, C. M. Whittington, B. M. McAllan, and V. J. Lynch. Gene expression phylogenies and ancestral transcriptome reconstruction resolves major transitions in the origins of pregnancy. eLife, 11:e74297, June 2022. ISSN 2050-084X. doi: 10.7554/eLife.74297.

L. M. Mikhaylova and D. I. Nurminsky. Lack of global meiotic sex chromosome inactivation, and paucity of tissue-specific gene expression on the Drosophila X chromosome. BMC Biology, 9(1):29, May 2011. ISSN 1741-7007. doi: 10.1186/1741-7007-9-29.

S. Molina-Gil, S. Sotillos, J. M. Espinosa-Vázquez, I. Almudi, and J. C.-G. Hombíıa. Interlocking of co-opted developmental gene networks in Drosophila and the evolution of preadaptive novelty. Nature Communications, 14(1):5730, Sept. 2023. ISSN 2041-1723. doi: 10.1038/s41467-023-41414-3.

A. Mortazavi, B. A. Williams, K. McCue, L. Schaeffer, and B. Wold. Mapping and quantifying mammalian transcriptomes by RNA-Seq. Nature Methods, 5(7):621–628, July 2008. ISSN 1548-7105. doi: 10.1038/nmeth.1226.

J. L. Mueller, K. Ravi Ram, L. A. McGraw, M. C. Bloch Qazi, E. D. Siggia, A. G. Clark, C. F. Aquadro, and M. F. Wolfner. Cross-species comparison of Drosophila male accessory gland protein genes. Genetics, 171(1):131–143, Sept. 2005. ISSN 0016-6731. doi: 10.1534/ge-netics.105.043844.

J. M. Musser and G. P. Wagner. Character trees from transcriptome data: Origin and individuation of morphological characters and the so-called “species signal”. Journal of Experimental Zoology Part B: Molecular and Developmental Evolution, 324(7):588–604, 2015. ISSN 1552-5015. doi: 10.1002/jez.b.22636.

A. Nourmohammad, J. Rambeau, T. Held, V. Kovacova, J. Berg, and M. Lässig. Adaptive Evolution of Gene Expression in Drosophila. Cell Reports, 20(6):1385–1395, Aug. 2017. ISSN 2211-1247. doi: 10.1016/j.celrep.2017.07.033.

D. J. Obbard, J. Maclennan, K.-W. Kim, A. Rambaut, P. M. O’Grady, and F. M. Jiggins. Estimating Divergence Dates and Substitution Rates in the Drosophila Phylogeny. Molecular Biology and Evolution, 29(11):3459–3473, Nov. 2012. ISSN 0737-4038. doi: 10.1093/mol-bev/mss150.

S. Pal, B. Oliver, and T. M. Przytycka. Stochastic Modeling of Gene Expression Evolution Uncovers Tissueand Sex-Specific Properties of Expression Evolution in the Drosophila Genus. Journal of Computational Biology: A Journal of Computational Molecular Cell Biology, 30(1):21–40, Jan. 2023. ISSN 1557-8666. doi: 10.1089/cmb.2022.0121.

M. Parisi, R. Nuttall, D. Naiman, G. Bouffard, J. Malley, J. Andrews, S. Eastman, and B. Oliver. Paucity of genes on the Drosophila X chromosome showing male-biased expression. *Science (New York*, N.Y.), 299(5607):697–700, Jan. 2003. ISSN 1095-9203. doi: 10.1126/sci-ence.1079190.

M. Parisi, R. Nuttall, P. Edwards, J. Minor, D. Naiman, J. Lü, M. Doctolero, M. Vainer, C. Chan, J. Malley, S. Eastman, and B. Oliver. A survey of ovary-, testis-, and soma-biased gene expression in Drosophila melanogasteradults. Genome Biology, 5(6):R40, June 2004. ISSN 1474-760X. doi: 10.1186/gb-2004-5-6-r40.

B. Patlar, V. Jayaswal, J. M. Ranz, and A. Civetta. Nonadaptive molecular evolution of seminal fluid proteins in Drosophila. Evolution; International Journal of Organic Evolution, 75(8): 2102–2113, Aug. 2021. ISSN 1558-5646. doi: 10.1111/evo.14297.

M. W. Pennell, J. M. Eastman, G. J. Slater, J. W. Brown, J. C. Uyeda, R. G. FitzJohn, M. E. Alfaro, and L. J. Harmon. Geiger v2.0: An expanded suite of methods for fitting macroevolutionary models to phylogenetic trees. Bioinformatics, 30(15):2216–2218, Aug. 2014. ISSN 1367-4803. doi: 10.1093/bioinformatics/btu181.

M. Pertea, G. M. Pertea, C. M. Antonescu, T.-C. Chang, J. T. Mendell, and S. L. Salzberg. StringTie enables improved reconstruction of a transcriptome from RNA-seq reads. Nature Biotechnology, 33(3):290–295, Mar. 2015. ISSN 1546-1696. doi: 10.1038/nbt.3122.

T. P. Quinn, T. M. Crowley, and M. F. Richardson. Benchmarking differential expression analysis tools for RNA-Seq: Normalization-based vs. log-ratio transformation-based methods. BMC Bioinformatics, 19(1):274, July 2018. ISSN 1471-2105. doi: 10.1186/s12859-018-2261-8.

J. M. Ranz, C. I. Castillo-Davis, C. D. Meiklejohn, and D. L. Hartl. Sex-dependent gene expression and evolution of the Drosophila transcriptome. *Science (New York*, N.Y.), 300 (5626):1742–1745, June 2003. ISSN 1095-9203. doi: 10.1126/science.1085881.

K. Ravi Ram and M. F. Wolfner. Seminal influences: Drosophila Acps and the molecular interplay between males and females during reproduction. Integrative and Comparative Biology, 47(3):427–445, Sept. 2007. ISSN 1557-7023. doi: 10.1093/icb/icm046.

M. Rebeiz, J. E. Pool, V. A. Kassner, C. F. Aquadro, and S. B. Carroll. Stepwise modification of a modular enhancer underlies adaptation in a Drosophila population. Science (New York, N.Y.), 326(5960):1663–1667, Dec. 2009. ISSN 1095-9203. doi: 10.1126/science.1178357.

S. A. Rifkin, J. Kim, and K. P. White. Evolution of gene expression in the Drosophila melanogaster subgroup. Nature Genetics, 33(2):138–144, Feb. 2003. ISSN 1061-4036. doi: 10.1038/ng1086.

R. V. Rohlfs, P. Harrigan, and R. Nielsen. Modeling Gene Expression Evolution with an Extended Ornstein–Uhlenbeck Process Accounting for Within-Species Variation. Molecular Biology and Evolution, 31(1):201–211, Jan. 2014. ISSN 0737-4038. doi: 10.1093/molbev/mst190.

I. G. Romero, I. Ruvinsky, and Y. Gilad. Comparative studies of gene expression and the evolution of gene regulation. Nature Reviews Genetics, 13(7):505–516, July 2012. ISSN 1471-0056. doi: 10.1038/nrg3229.

B. Schwanhäusser, D. Busse, N. Li, G. Dittmar, J. Schuchhardt, J. Wolf, W. Chen, and M. Selbach. Global quantification of mammalian gene expression control. Nature, 473(7347): 337–342, May 2011. ISSN 1476-4687. doi: 10.1038/nature10098.

N. D. Singh and D. A. Petrov. Evolution of gene function on the X chromosome versus the autosomes. Genome Dynamics, 3:101–118, 2007. ISSN 1660-9263. doi: 10.1159/000107606.

D. L. Stern and V. Orgogozo. The loci of evolution: How predictable is genetic evolution? Evolution; International Journal of Organic Evolution, 62(9):2155–2177, Sept. 2008. ISSN 0014-3820. doi: 10.1111/j.1558-5646.2008.00450.x.

K. Struhl. Transcriptional noise and the fidelity of initiation by RNA polymerase II. Nature Structural & Molecular Biology, 14(2):103–105, Feb. 2007. ISSN 1545-9993. doi: 10.1038/nsmb0207-103.

D. Sturgill, Y. Zhang, M. Parisi, and B. Oliver. Demasculinization of X chromosomes in the Drosophila genus. Nature, 450(7167):238–241, Nov. 2007. ISSN 1476-4687. doi: 10.1038/nature06330.

A. Suvorov, B. Y. Kim, J. Wang, E. E. Armstrong, D. Peede, E. R. D’Agostino, D. K. Price, P. Waddell, M. Lang, V. Courtier-Orgogozo, J. R. David, D. Petrov, D. R. Matute, D. R. Schrider, and A. A. Comeault. Widespread introgression across a phylogeny of 155 Drosophila genomes. Current Biology, page S0960982221014962, Nov. 2021. ISSN 09609822. doi: 10.1016/j.cub.2021.10.052.

M. Suyama, D. Torrents, and P. Bork. PAL2NAL: Robust conversion of protein sequence alignments into the corresponding codon alignments. Nucleic Acids Research, 34(suppl 2): W609–W612, July 2006. ISSN 0305-1048. doi: 10.1093/nar/gkl315.

Y. A. Takashima, A. C. Majane, and D. J. Begun. Evolution of secondary cell number and position in the Drosophila accessory gland. PLOS ONE, 18(10):e0278811, Oct. 2023. ISSN 1932-6203. doi: 10.1371/journal.pone.0278811.

K. Tanaka, O. Barmina, L. E. Sanders, M. N. Arbeitman, and A. Kopp. Evolution of sex-specific traits through changes in HOX-dependent doublesex expression. PLoS biology, 9 (8):e1001131, Aug. 2011. ISSN 1545-7885. doi: 10.1371/journal.pbio.1001131.

A. Thompson, H. H. Zakon, and M. Kirkpatrick. Compensatory Drift and the Evolutionary Dynamics of Dosage-Sensitive Duplicate Genes. Genetics, 202(2):765–774, Feb. 2016. ISSN 1943-2631. doi: 10.1534/genetics.115.178137.

A. Thompson, D. T. Infield, A. R. Smith, G. T. Smith, C. A. Ahern, and H. H. Zakon. Rapid evolution of a voltage-gated sodium channel gene in a lineage of electric fish leads to a persistent sodium current. PLOS Biology, 16(3):e2004892, Mar. 2018. ISSN 1545-7885. doi: 10.1371/journal.pbio.2004892.

A. Thompson, M. R. May, B. R. Moore, and A. Kopp. A hierarchical Bayesian mixture model for inferring the expression state of genes in transcriptomes. Proceedings of the National Academy of Sciences, 117(32):19339–19346, Aug. 2020. ISSN 0027-8424, 1091-6490. doi: 10.1073/pnas.1919748117.

E. V. Todd, M. A. Black, and N. J. Gemmell. The power and promise of RNA-seq in ecology and evolution. Molecular Ecology, 25(6):1224–1241, Mar. 2016. ISSN 1365-294X. doi: 10.1111/mec.13526.

M. Turelli, B. S. Cooper, K. M. Richardson, P. S. Ginsberg, B. Peckenpaugh, C. X. Antelope, K. J. Kim, M. R. May, A. Abrieux, D. A. Wilson, M. J. Bronski, B. R. Moore, J.-J. Gao, M. B. Eisen, J. C. Chiu, W. R. Conner, and A. A. Hoffmann. Rapid Global Spread of wRilike Wolbachia across Multiple Drosophila. Current biology: CB, 28(6):963–971.e8, Mar. 2018. ISSN 1879-0445. doi: 10.1016/j.cub.2018.02.015.

J. Vandesompele, K. De Preter, F. Pattyn, B. Poppe, N. Van Roy, A. De Paepe, and F. Speleman. Accurate normalization of real-time quantitative RT-PCR data by geometric averaging of multiple internal control genes. Genome Biology, 3(7):research0034.1, June 2002. ISSN 1474-760X. doi: 10.1186/gb-2002-3-7-research0034.

M. D. Vibranovski, Y. Zhang, and M. Long. General gene movement off the X chromosome in the Drosophila genus. Genome Research, 19(5):897–903, May 2009. ISSN 1088-9051. doi: 10.1101/gr.088609.108.

G. P. Wagner, K. Kin, and V. J. Lynch. A model based criterion for gene expression calls using RNA-seq data. Theory in Biosciences, 132(3):159–164, Sept. 2013. ISSN 1611-7530. doi: 10.1007/s12064-013-0178-3.

Z. Wang, M. Gerstein, and M. Snyder. RNA-Seq: A revolutionary tool for transcriptomics. Nature Reviews. Genetics, 10(1):57–63, Jan. 2009. ISSN 1471-0064. doi: 10.1038/nrg2484.

P. J. Wittkopp and G. Kalay. Cis-regulatory elements: Molecular mechanisms and evolutionary processes underlying divergence. Nature Reviews Genetics, 13(1):59–69, Jan. 2012. ISSN 1471-0064. doi: 10.1038/nrg3095.

G. A. Wray. The evolutionary significance of cis-regulatory mutations. Nature Reviews. Genetics, 8(3):206–216, Mar. 2007. ISSN 1471-0056. doi: 10.1038/nrg2063.

Z. Yang. Maximum likelihood phylogenetic estimation from DNA sequences with variable rates over sites: Approximate methods. Journal of Molecular Evolution, 39(3):306–314, Sept. 1994. ISSN 0022-2844, 1432-1432. doi: 10.1007/BF00160154.

Z. Yang and B. Rannala. Bayesian phylogenetic inference using DNA sequences: A Markov Chain Monte Carlo Method. Molecular Biology and Evolution, 14(7):717–724, July 1997. ISSN 0737-4038. doi: 10.1093/oxfordjournals.molbev.a025811.

G. Yu, L.-G. Wang, Y. Han, and Q.-Y. He. clusterProfiler: An R package for comparing biological themes among gene clusters. Omics: A Journal of Integrative Biology, 16(5):284–287, May 2012. ISSN 1557-8100. doi: 10.1089/omi.2011.0118.

Y. Zhang, D. Sturgill, M. Parisi, S. Kumar, and B. Oliver. Constraint and turnover in sex-biased gene expression in the genus Drosophila. Nature, 450(7167):233–237, Nov. 2007. ISSN 1476-4687. doi: 10.1038/nature06323.

Y. Zhou, J. Zhu, T. Tong, J. Wang, B. Lin, and J. Zhang. A statistical normalization method and differential expression analysis for RNA-seq data between different species. BMC Bioinformatics, 20(1):163, Mar. 2019. ISSN 1471-2105. doi: 10.1186/s12859-019-2745-1.

