## Supplemental_text_S1 for "Quantifying transcriptome turnover on phylogenies by modeling gene expression as a binary trait"

### Supplemental text S1: Quantifying transcriptome turnover on phylogenies by modeling gene expression as a binary trait

### Branch rate estimates

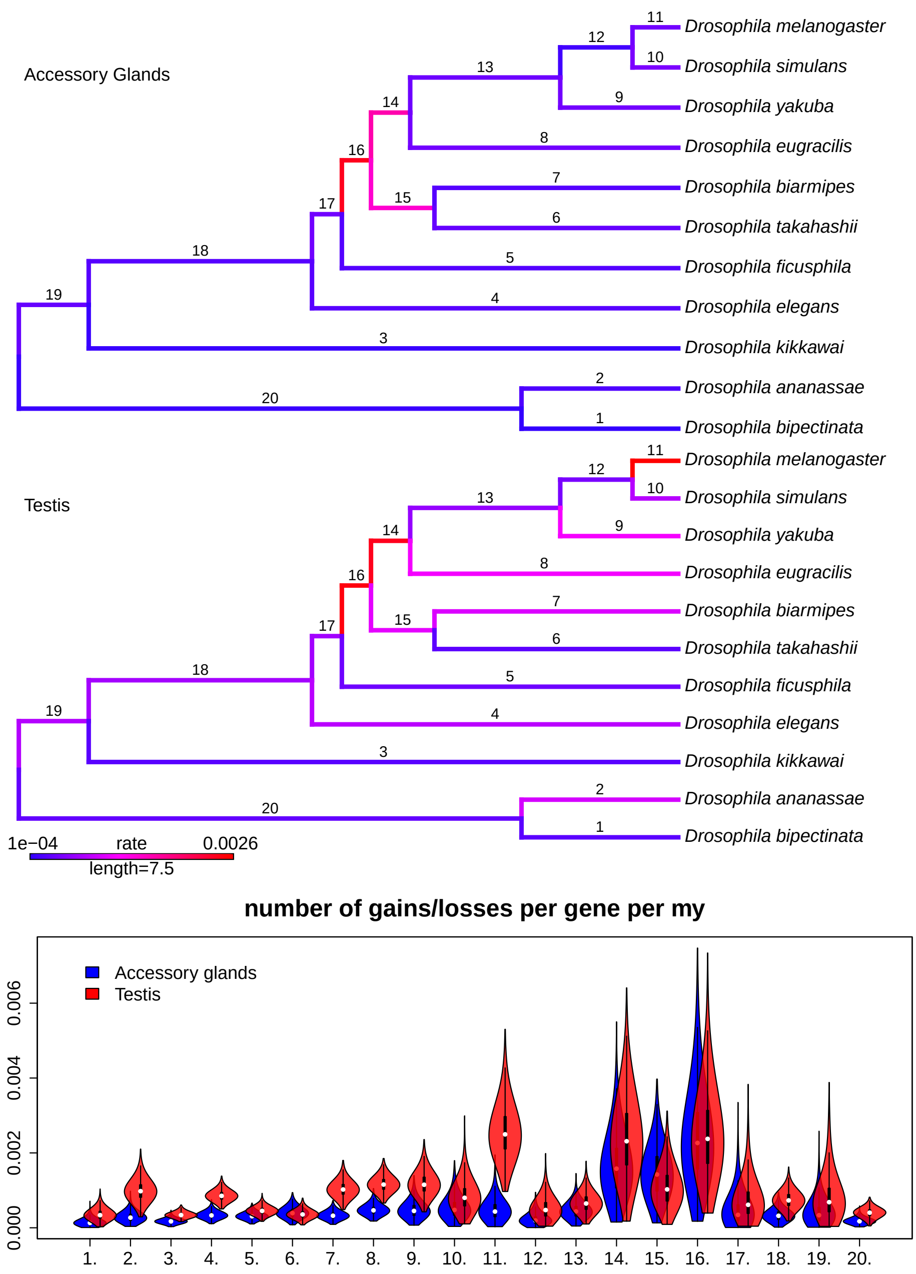


Figure S1: Posterior mean estimations of branch-specific relative turnover rates for single-copy genes in the AG (top phylogeny) and testis (bottom phylogeny) with threshold at 0.01 and 0.99. Branch color reflects the mean posterior estimates of the transcriptome turnover rate (changes per single-copy gene per MY), from low (blue) to high (red). The bottom violin plot shows the full posterior samples of the relative turnover rate for AG (blue) and testis (red) for each branch numbered on the phylogenetic trees.

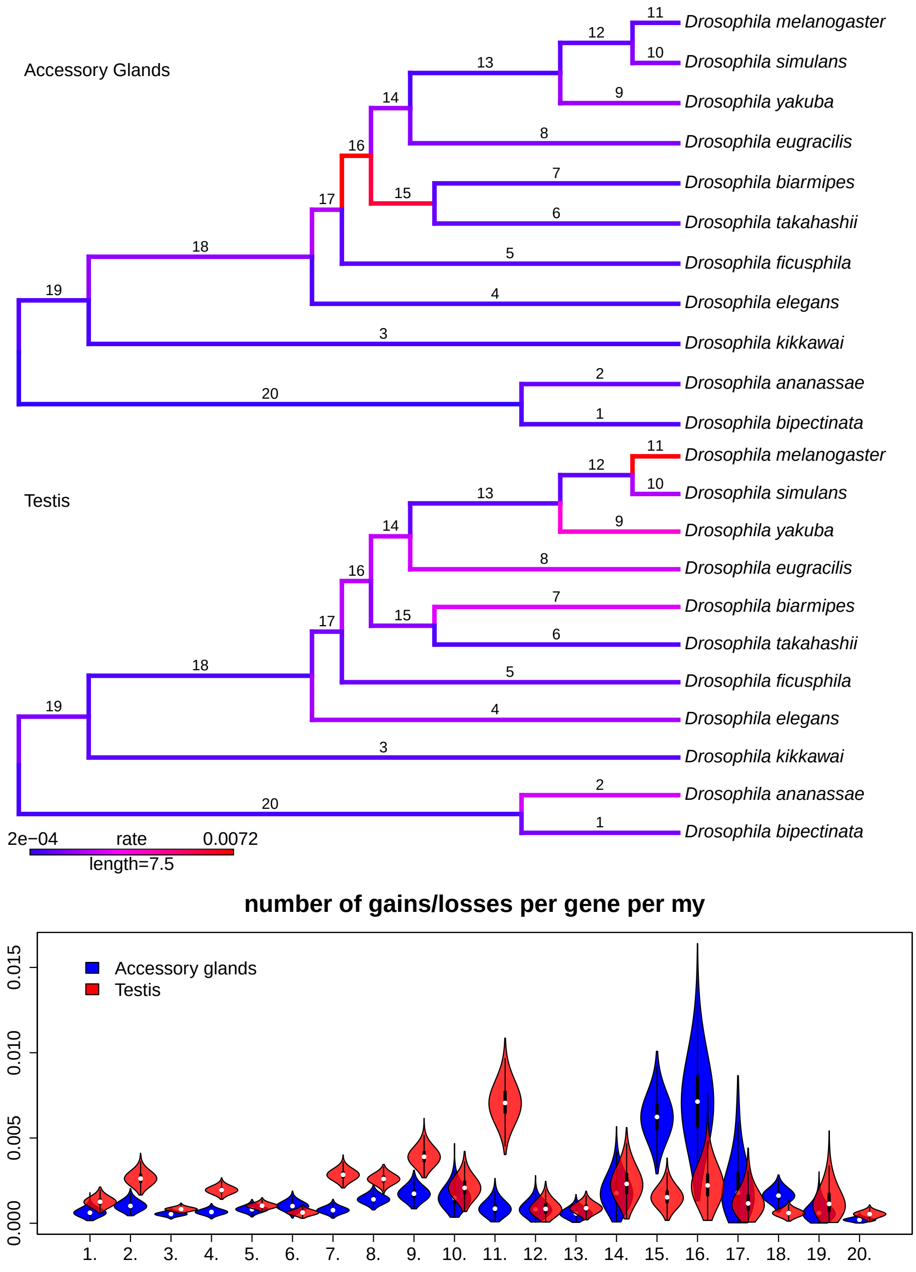


Figure S2: Threshold at 0.05 and 0.95.

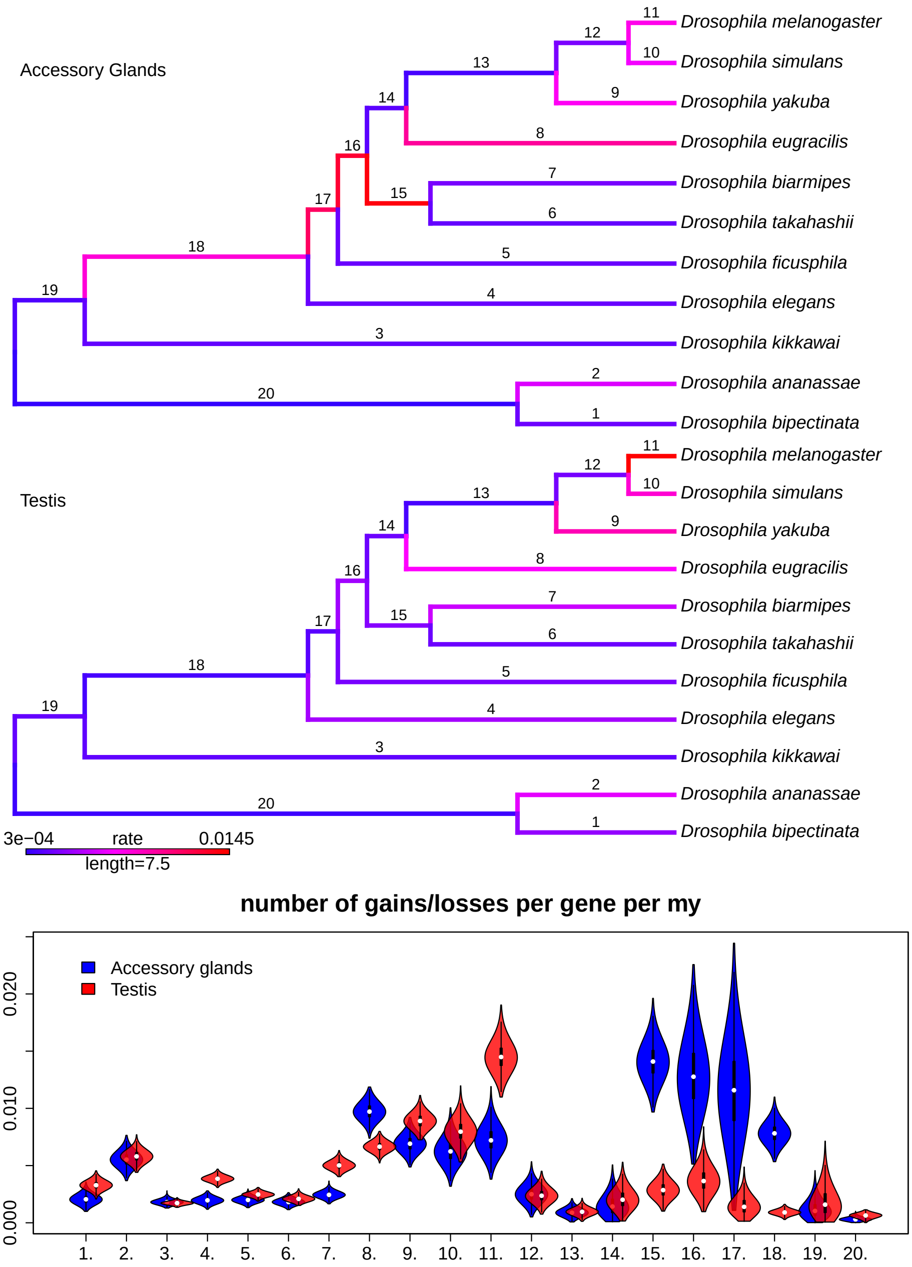


Figure S3: Threshold at 0.25 and 0.75.

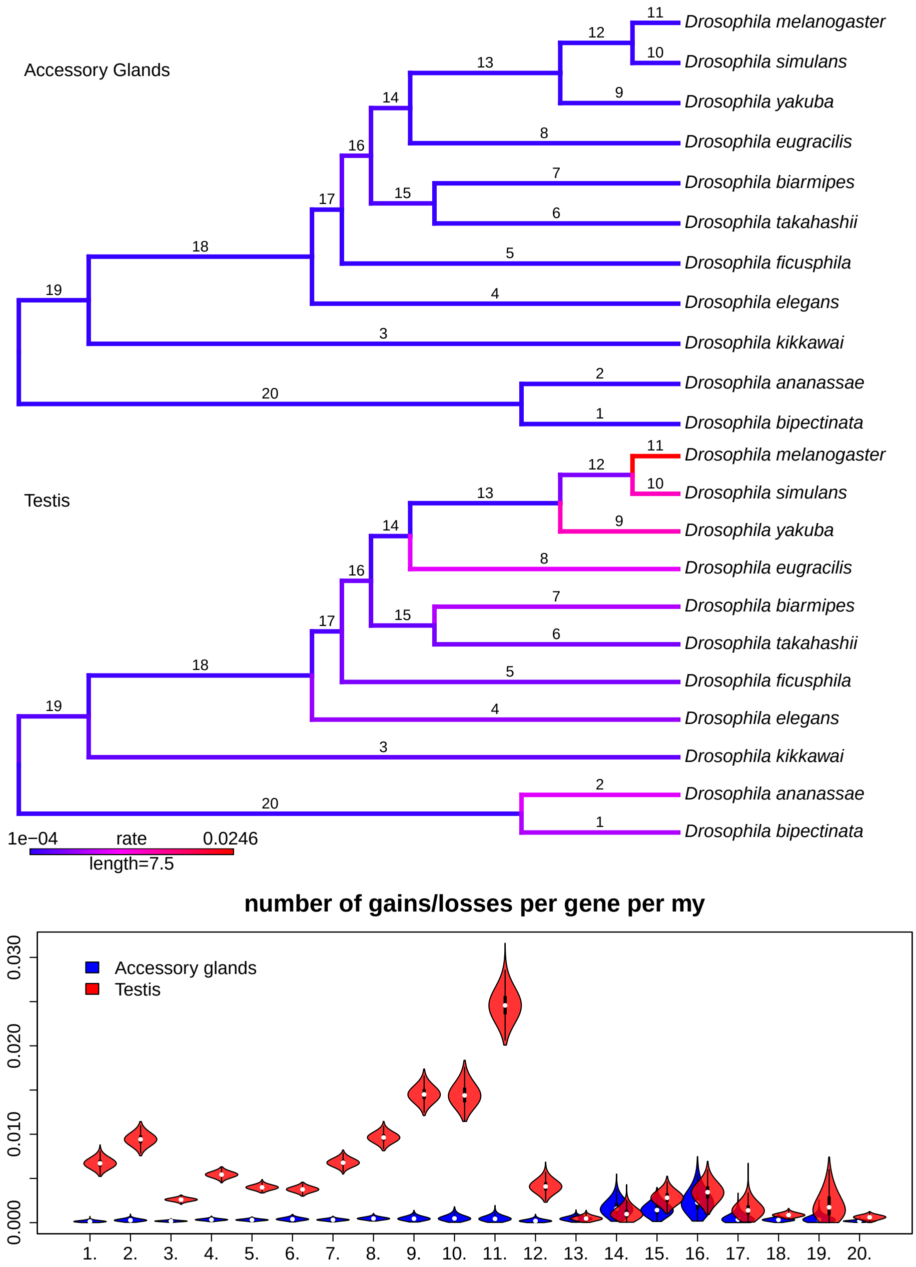


Figure S4: Threshold at 0.5.

### Cross-validation

#### Methods

If *zigzag* provides an adequate model of the relationship between RNA-seq data and expression state (Thompson et al., 2020), and the phylogenetic model adequately approximates the process that gave rise to the comparative patterns of these states among the eleven species, we should be able to split the data and fit the model to a subset of the data and use that model to predict the states of the holdout data. We expect that for genes withheld from phylogenetic inference in some species, the marginal probability of their inferred states under the phylogenetic model should agree with the probabilities assigned by *zigzag*. To test whether this agreement holds, we randomly selected 100 genes from each species and each organ and removed them from the phylogenetic inference by coding their expression states as missing data. This amounted to 2,200 total genes, or roughly 1% of the data. We then fit the evolutionary model to the remaining 99% of the data and sampled the marginal posterior distribution of the expression states of the missing genes given the model and the remaining data. This was computed in the same way as the ancestral character states.

We fit the phylogenetic model to the expression data using five different probability thresholds for classifying gene expression states; we describe these thresholds as $\alpha$ values of 0.5, 0.25, 0.1, 0.05, 0.01. Genes with probabilities of being active that are greater than (1 - $\alpha$) are coded as active, those below $\alpha$ are coded as inactive, and genes that fall between $\alpha$ and (1 - $\alpha$) are coded as missing data. For example, for an $\alpha$ = 0.25, *zigzag* probabilities less than 0.25 were coded as inactive, probabilities between 0.25 and 0.75 were considered unknown (missing data), and genes with P  0.75 were coded as active.

To quantify the level of agreement between *zigzag* and the phylogenetic inference of the holdout set of genes, we computed the probability that the two methods predict the same expression state across this set of hold-out genes for each organ in each species, which we call the accuracy:

$$\text{Accuracy} = P(z_i = \text{active})P(\psi_i = \text{active}) + P(z_i = \text{inactive})P(\psi_i = \text{inactive}),$$

where $z_i$ denotes the *zigzag* state and $\psi_i$ the inferred phylogenetic state of gene $i$.

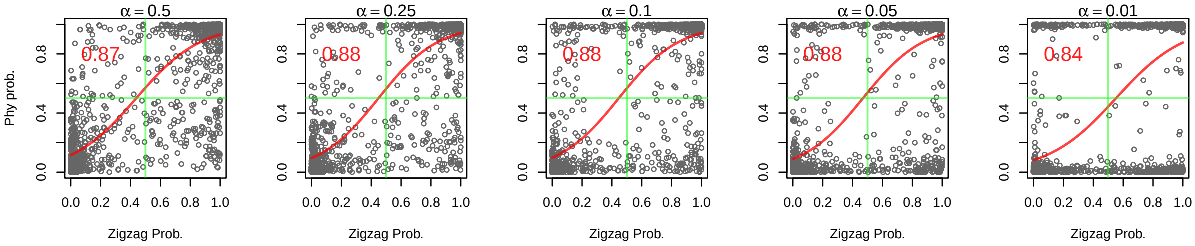


Figure S5: Cross-validation on hold-out gene set. Expression state of 100 random genes in each organ in each species were held-out before phylogenetic analysis under five different $\alpha$'s for classifying expression state. For example, $\alpha = 0.1$ indicates that genes with the posterior probability of expression $< 0.1$ are classified as inactive and those with $\alpha > 0.9$ as active, while the genes with intermediate probabilities are considered to have unknown expression states. At $\alpha = 0.5$, all genes are classified as either active or inactive. The red numbers in the upper left of each panel is the mean accuracy across the hold-out genes in both organs across all species (2,200 total genes).

#### Results

The phylogenetic inference of the hold-out gene set states correlated very well with the *zigzag* probabilities (Figures S5 and S6). All $\alpha$ values except 0.01 and 0.5 had similar mean accuracies with 0.1 having the highest. On average, the probability that the phylogenetic inference of expression state matched the *zigzag* inference was 0.88 for this value of $\alpha$. Main results presented in the main text are from analyses with $\alpha$ = 0.1.

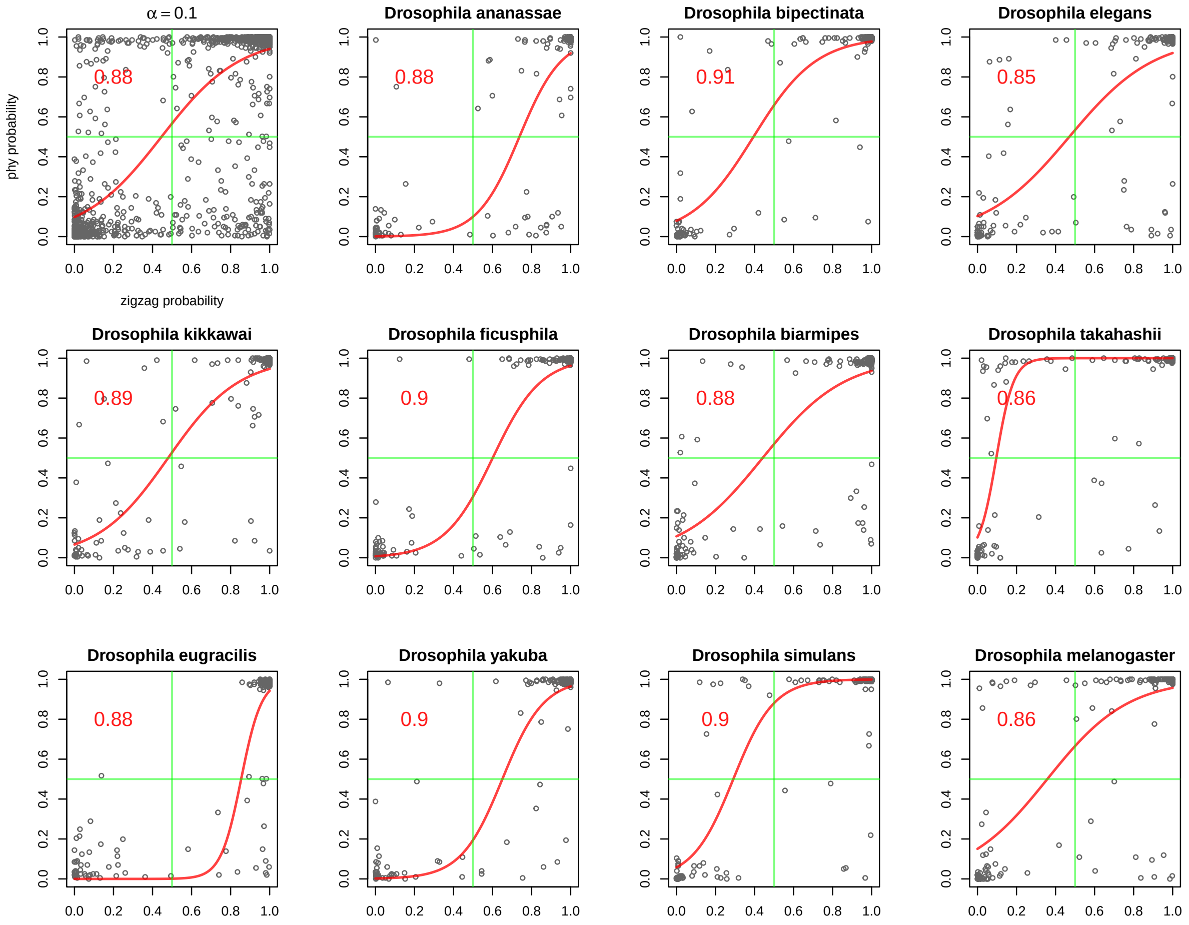


Figure S6: Comparison of inferred gene expression states given the phylogeny and evolution model (Y axis) and the probabilities of gene expression states predicted by *zigzag* given the expression data (X axis) for hold-out genes treated as missing data in the phylogenetic analysis in the accessory glands. The top left corner panel shows least-squares logistic regression fits (red curve) for all eleven species pooled. Each scatter plot shows the *zigzag* probability of active expression for genes treated as missing data (x-axis) and the ancestral state reconstruction of those genes under the phylogenetic model (y-axis), n = 2200 (100 per organ in each species). Overlaid are the fit logistic regressions for each species (red lines).

### GO Analysis

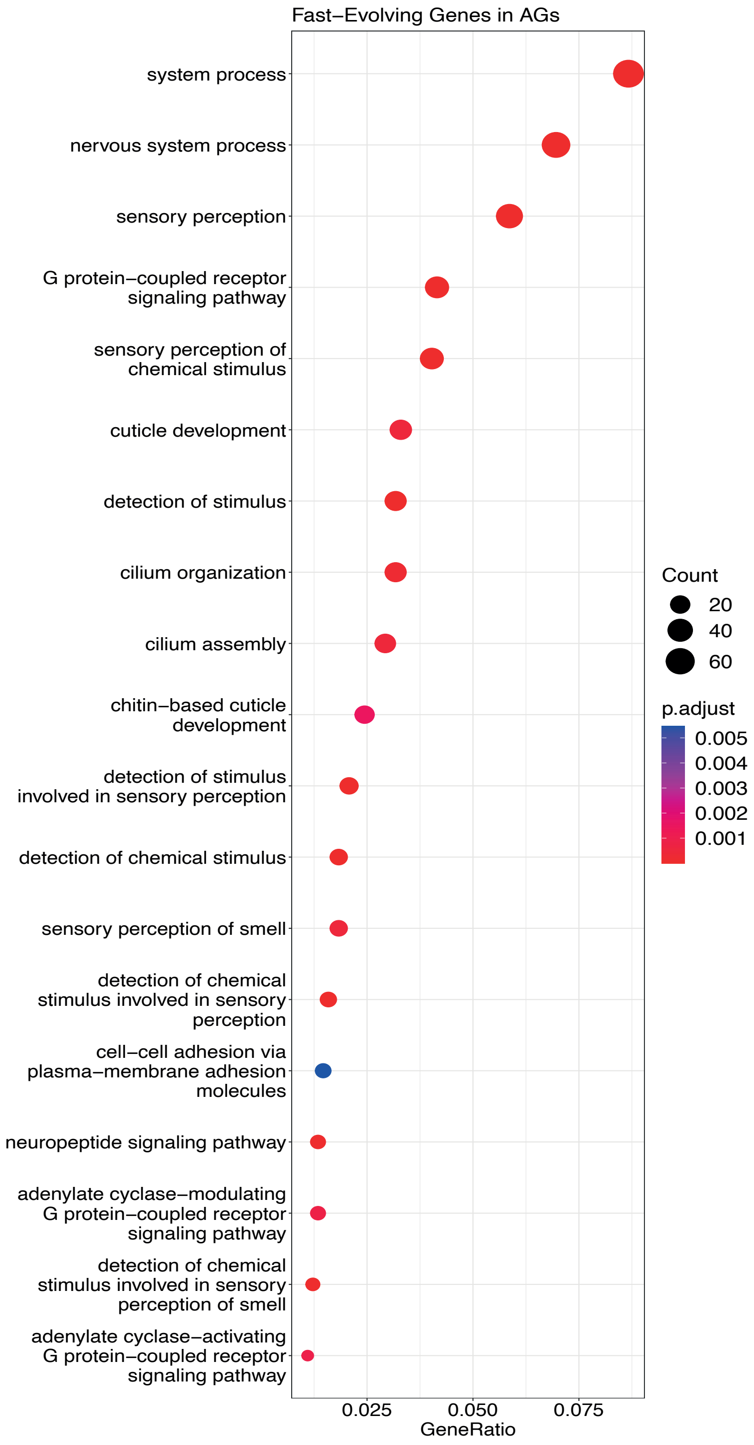


Figure S7: A dot plot from a gene ontology (GO) analysis showing all significantly enriched (q-value $< 0.05$) biological process terms among the genes showing rapid turnover in the AGs. Genes are designated as showing rapid turnover if they have rates at least two-fold higher than the mean. The list of all singleton genes expressed in the AGs is used as the background.

### References

A. Thompson, M. R. May, B. R. Moore, and A. Kopp. A hierarchical Bayesian mixture model for inferring the expression state of genes in transcriptomes. *Proceedings of the National Academy of Sciences*, 117(32):19339–19346, Aug. 2020. ISSN 0027-8424, 1091-6490. doi: 10.1073/pnas.1919748117.
