## Supplementary material for "Quantifying transcriptome turnover on phylogenies by modeling gene expression as a binary trait": Data_file_S1

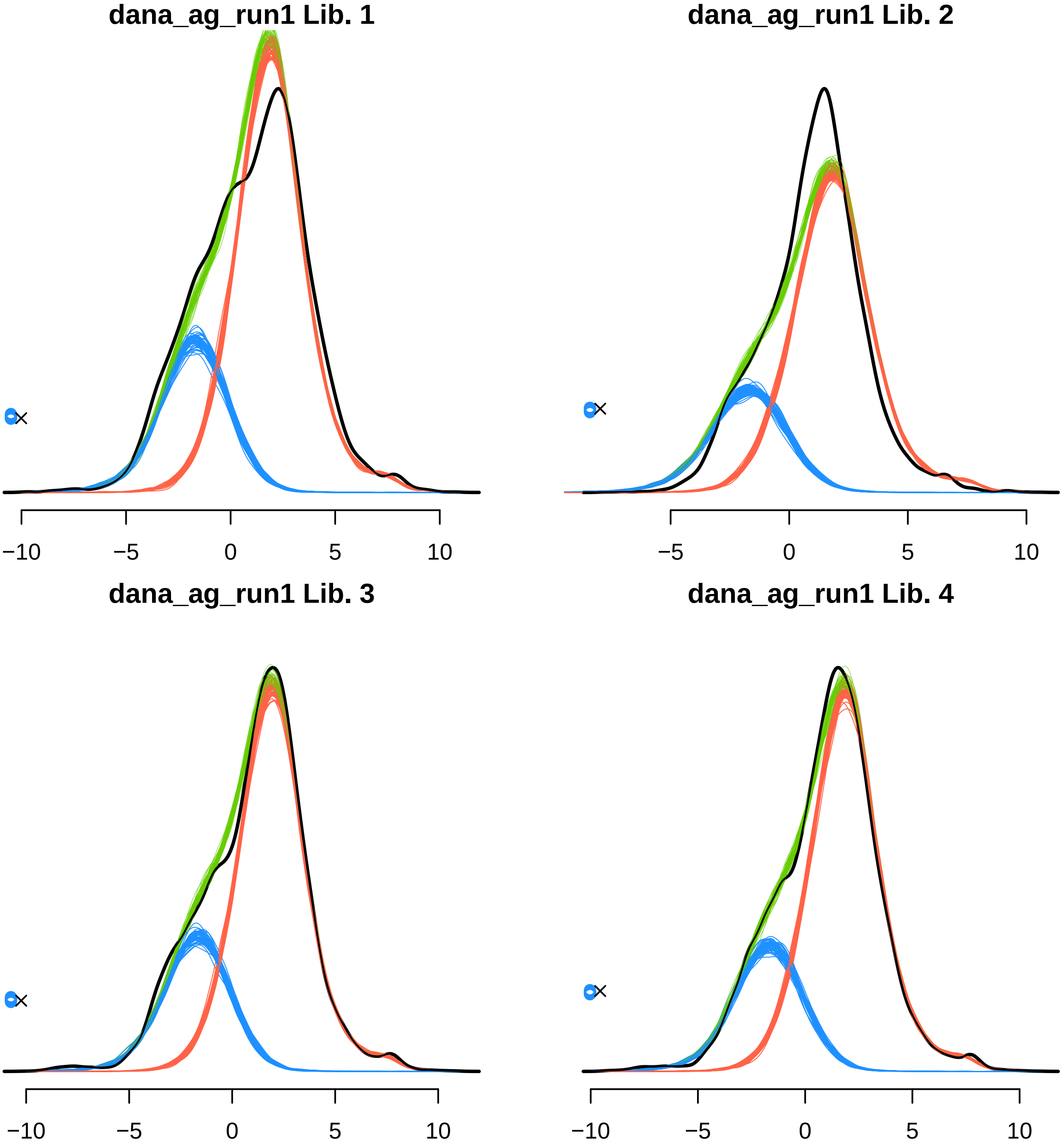

dana_ag_run1 Lib. 1
−10
−5
0
5
10
dana_ag_run1 Lib. 2
−5
0
5
10
dana_ag_run1 Lib. 3
−10
−5
0
5
10
dana_ag_run1 Lib. 4
−10
−5
0
5
10

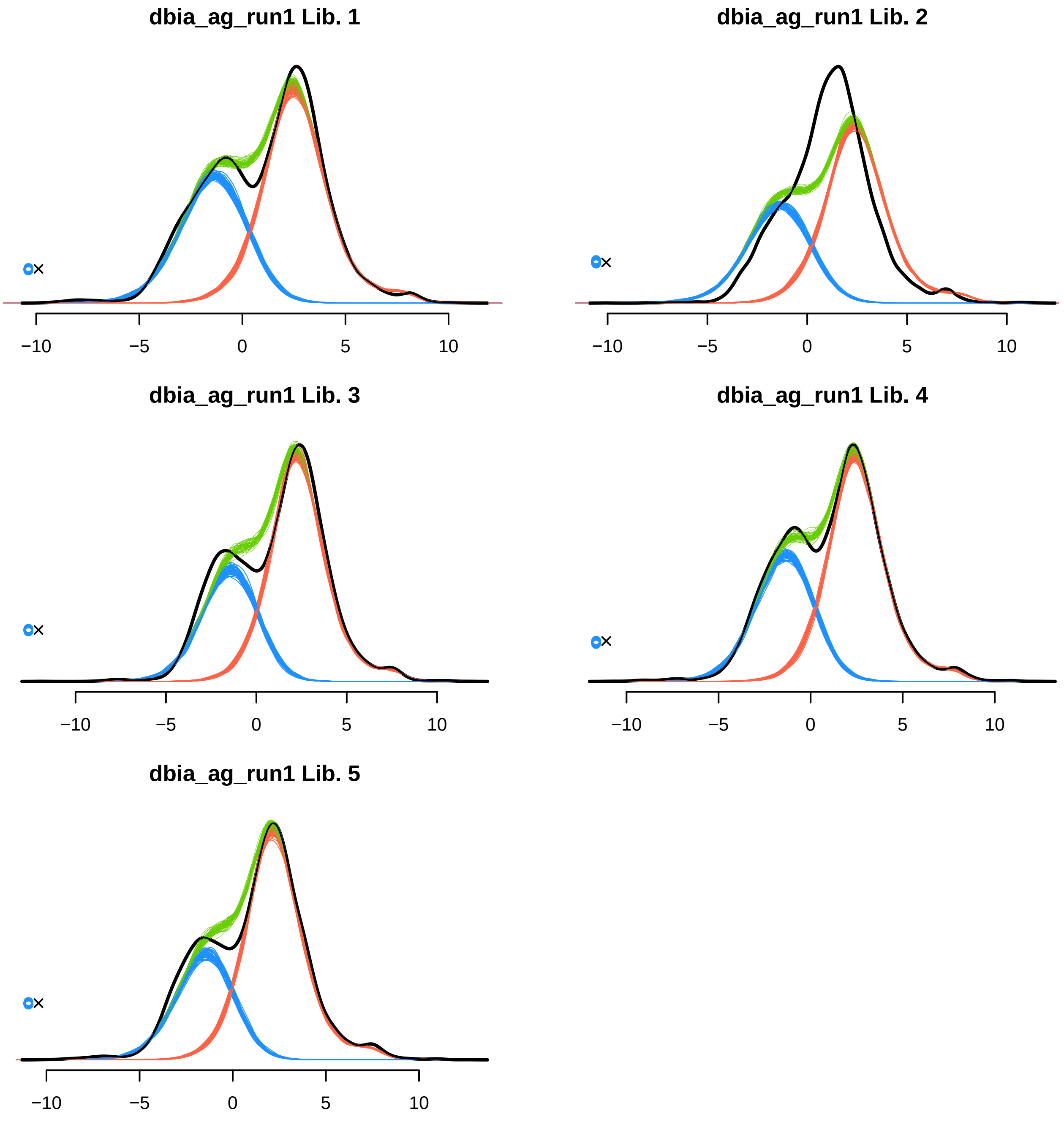
dbia_ag_run1 Lib. 1
−10
−5
0
5
10
dbia_ag_run1 Lib. 2
−10
−5
0
5
10
dbia_ag_run1 Lib. 3
−10
−5
0
5
10
dbia_ag_run1 Lib. 4
−10
−5
0
5
10
dbia_ag_run1 Lib. 5
−10
−5
0
5
10

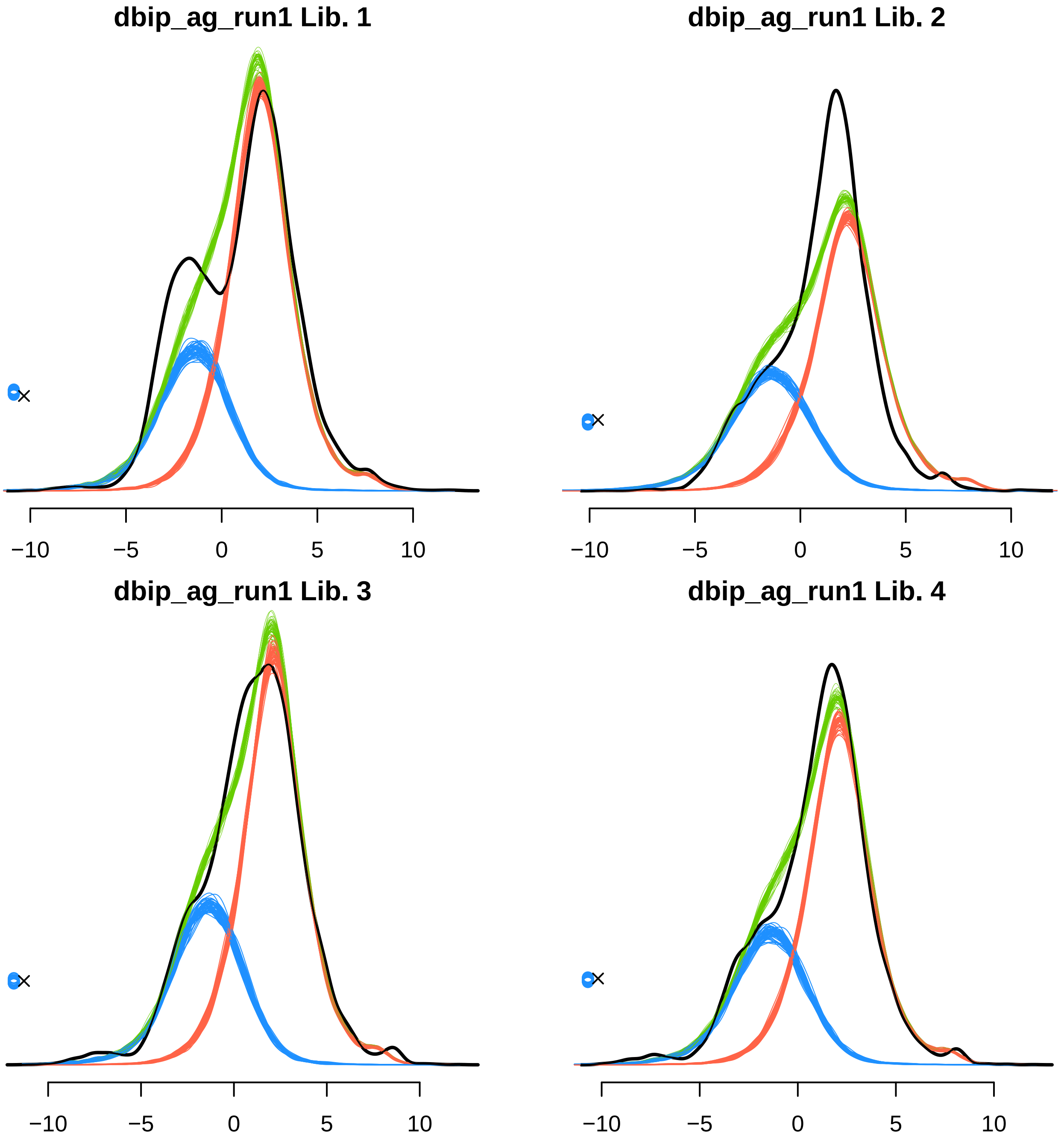
dbip_ag_run1 Lib. 1
−10
−5
0
5
10
dbip_ag_run1 Lib. 2
−10
−5
0
5
10
dbip_ag_run1 Lib. 3
−10
−5
0
5
10
dbip_ag_run1 Lib. 4
−10
−5
0
5
10

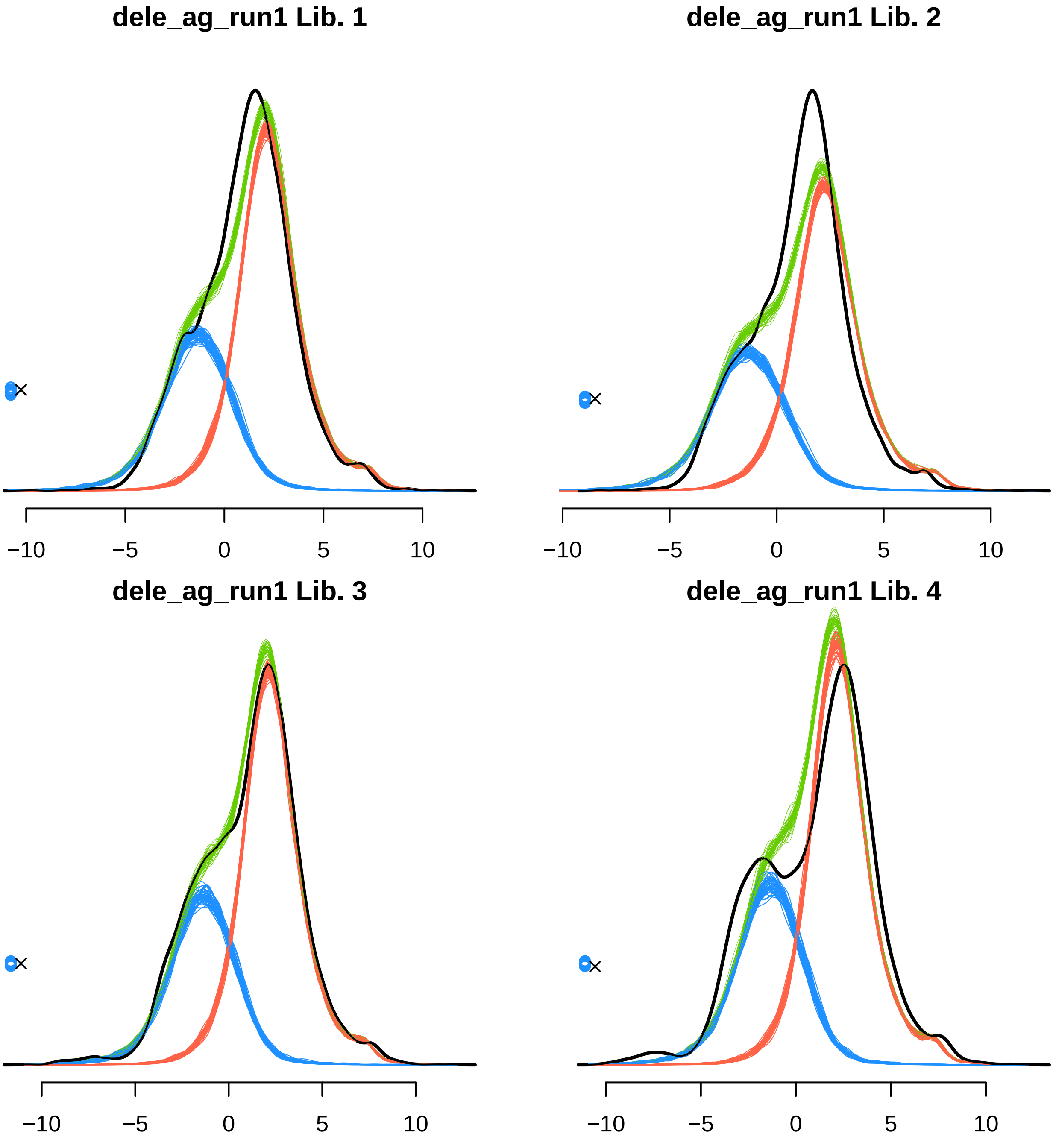

dele_ag_run1 Lib. 1
dele_ag_run1 Lib. 2
dele_ag_run1 Lib. 3
dele_ag_run1 Lib. 4
−10
−5
0
5
10

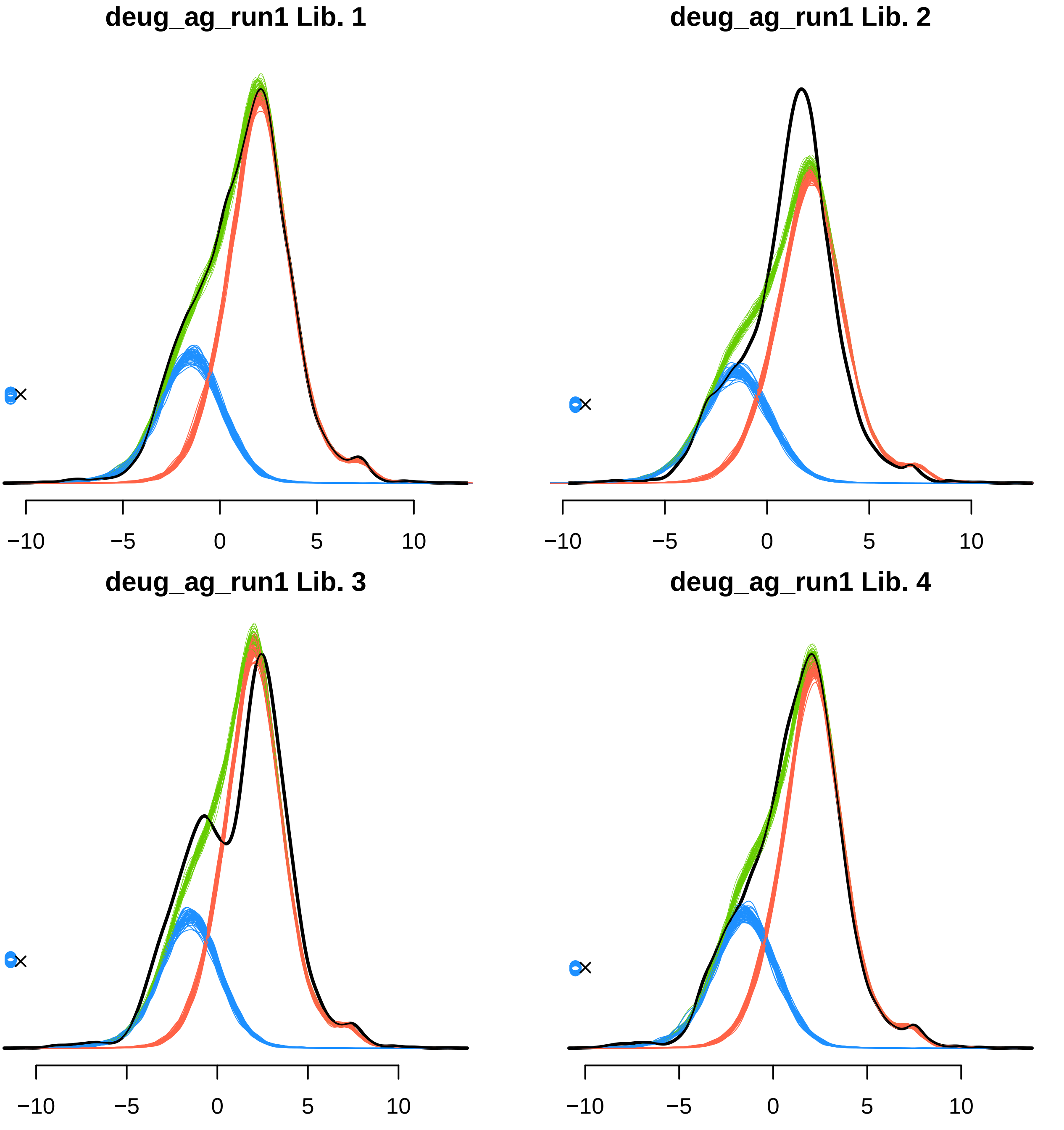
deug_ag_run1 Lib. 1
deug_ag_run1 Lib. 2
deug_ag_run1 Lib. 3
deug_ag_run1 Lib. 4
−10
−5
0
5
10

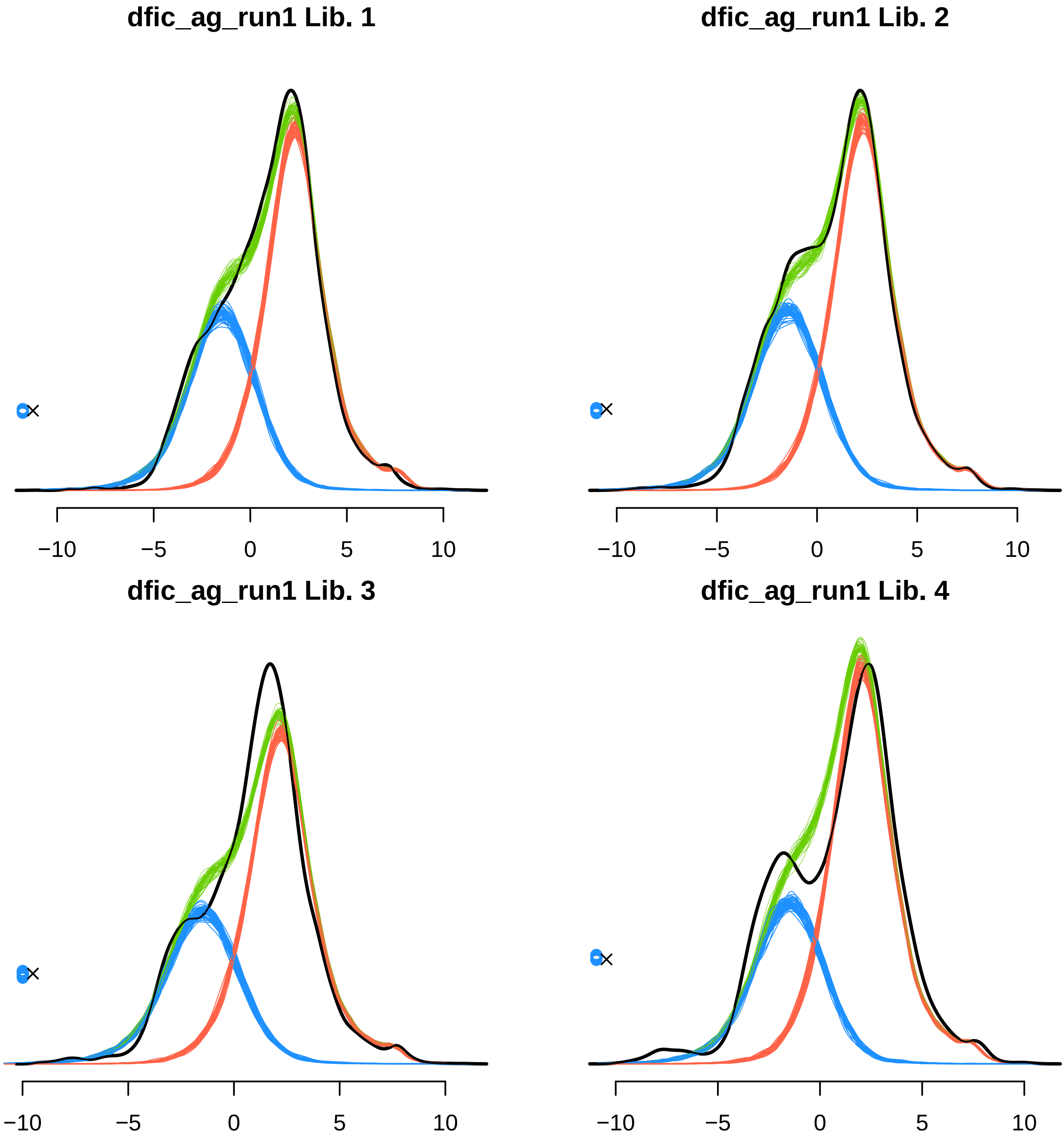

dfic_ag_run1 Lib. 1
−10
−5
0
5
10
dfic_ag_run1 Lib. 2
−10
−5
0
5
10
dfic_ag_run1 Lib. 3
−10
−5
0
5
10
dfic_ag_run1 Lib. 4
−10
−5
0
5
10

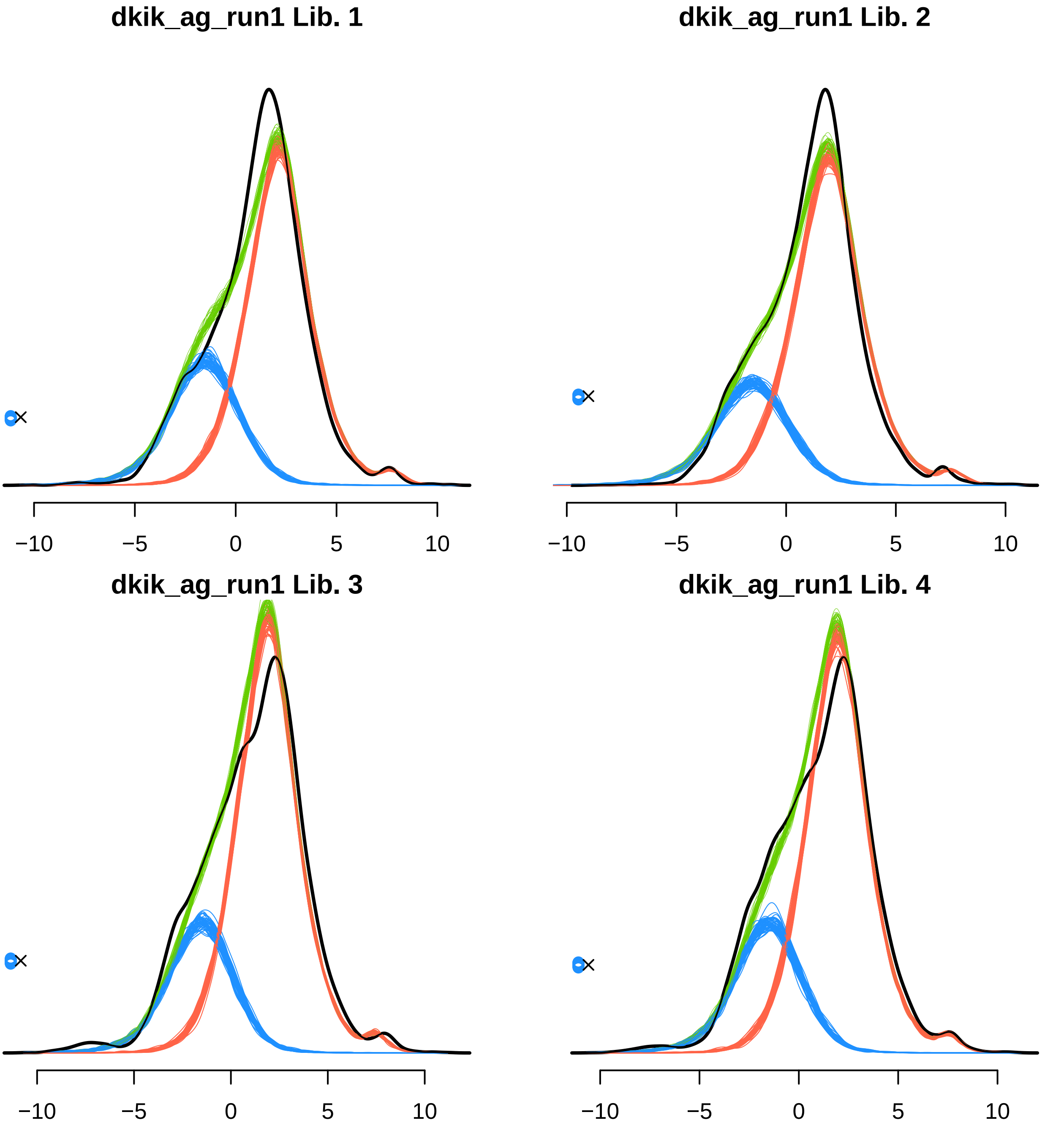
dkik_ag_run1 Lib. 1
dkik_ag_run1 Lib. 2
dkik_ag_run1 Lib. 3
dkik_ag_run1 Lib. 4
−10
−5
0
5
10

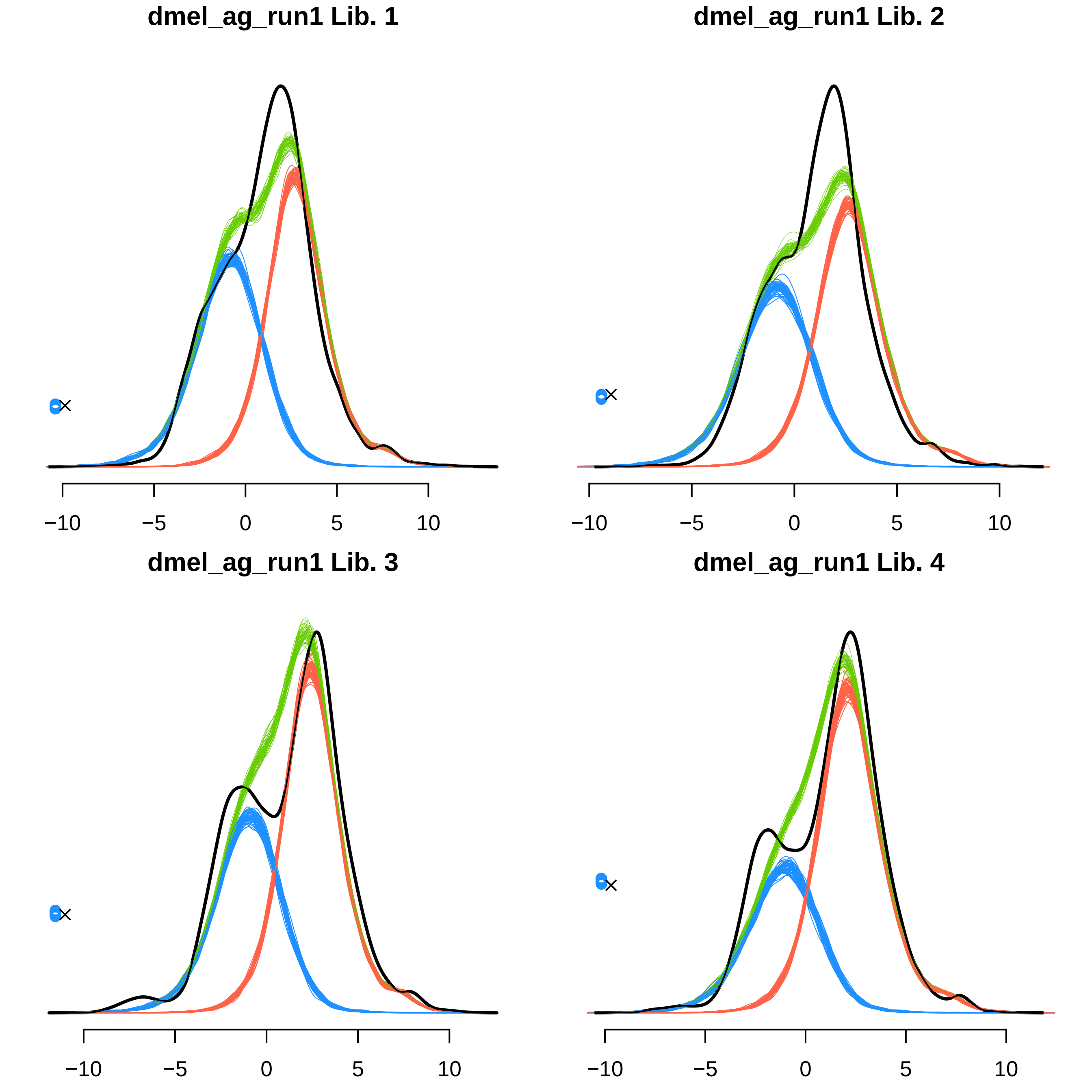
dmel_ag_run1 Lib. 1
−10
−5
0
5
10
dmel_ag_run1 Lib. 2
−10
−5
0
5
10
dmel_ag_run1 Lib. 3
−10
−5
0
5
10
dmel_ag_run1 Lib. 4
−10
−5
0
5
10

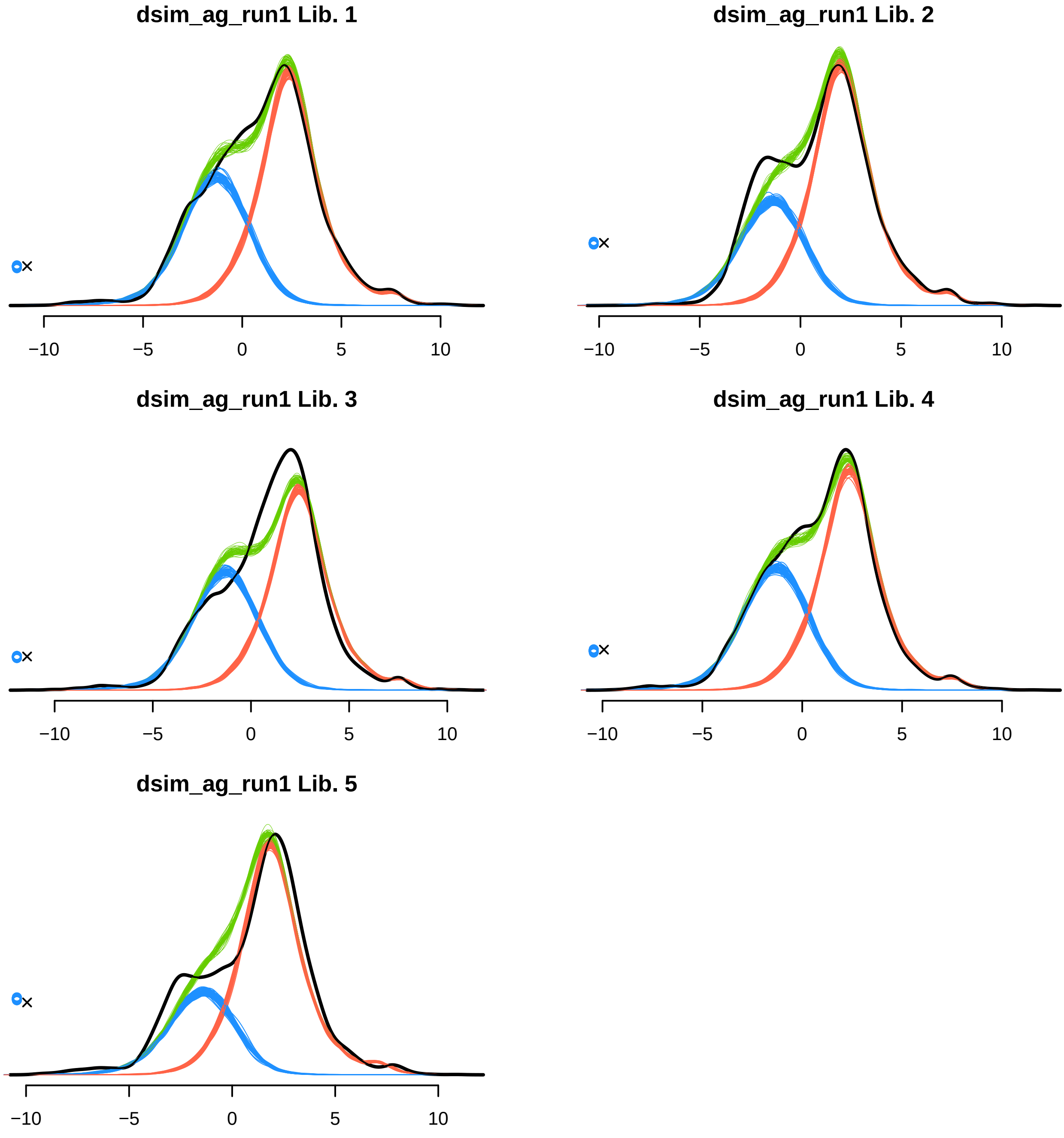
dsim_ag_run1 Lib. 1
−10
−5
0
5
10
dsim_ag_run1 Lib. 2
−10
−5
0
5
10
dsim_ag_run1 Lib. 3
−10
−5
0
5
10
dsim_ag_run1 Lib. 4
−10
−5
0
5
10
dsim_ag_run1 Lib. 5
−10
−5
0
5
10

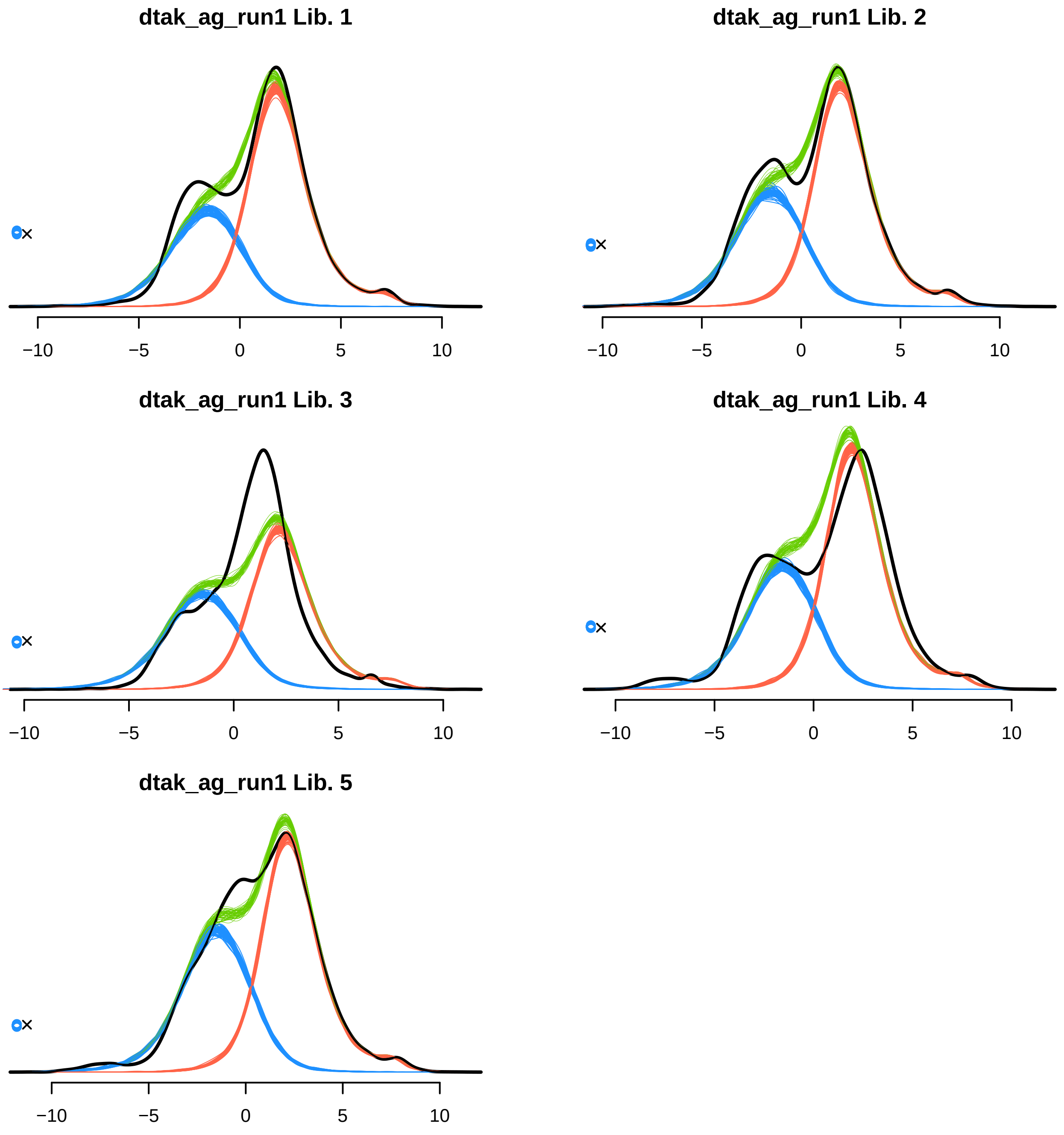

dtak_ag_run1 Lib. 1
−10
−5
0
5
10
dtak_ag_run1 Lib. 2
−10
−5
0
5
10
dtak_ag_run1 Lib. 3
−10
−5
0
5
10
dtak_ag_run1 Lib. 4
−10
−5
0
5
10
dtak_ag_run1 Lib. 5
−10
−5
0
5
10

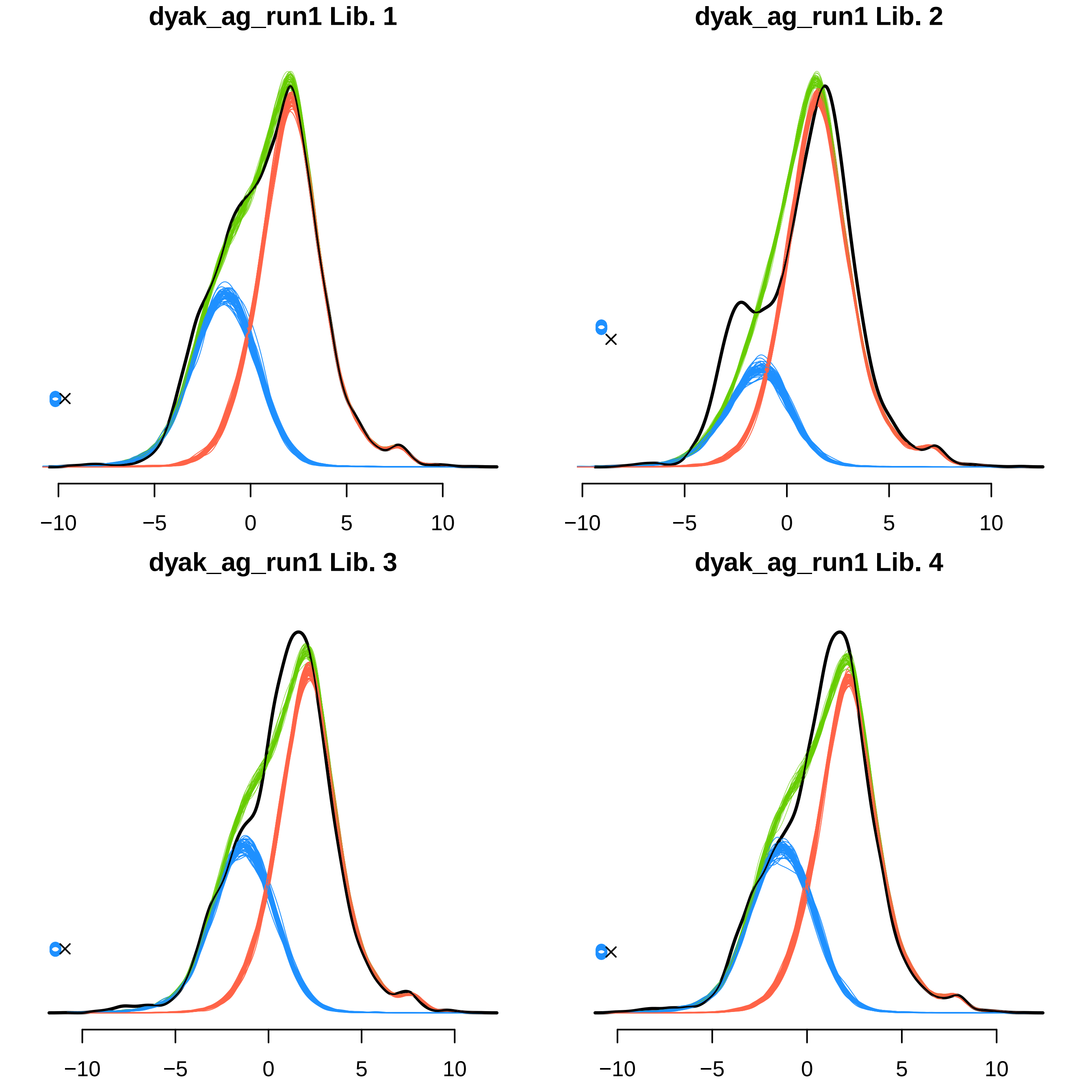

dyak_ag_run1 Lib. 1
−10
−5
0
5
10
dyak_ag_run1 Lib. 2
−10
−5
0
5
10
dyak_ag_run1 Lib. 3
−10
−5
0
5
10
dyak_ag_run1 Lib. 4
−10
−5
0
5
10

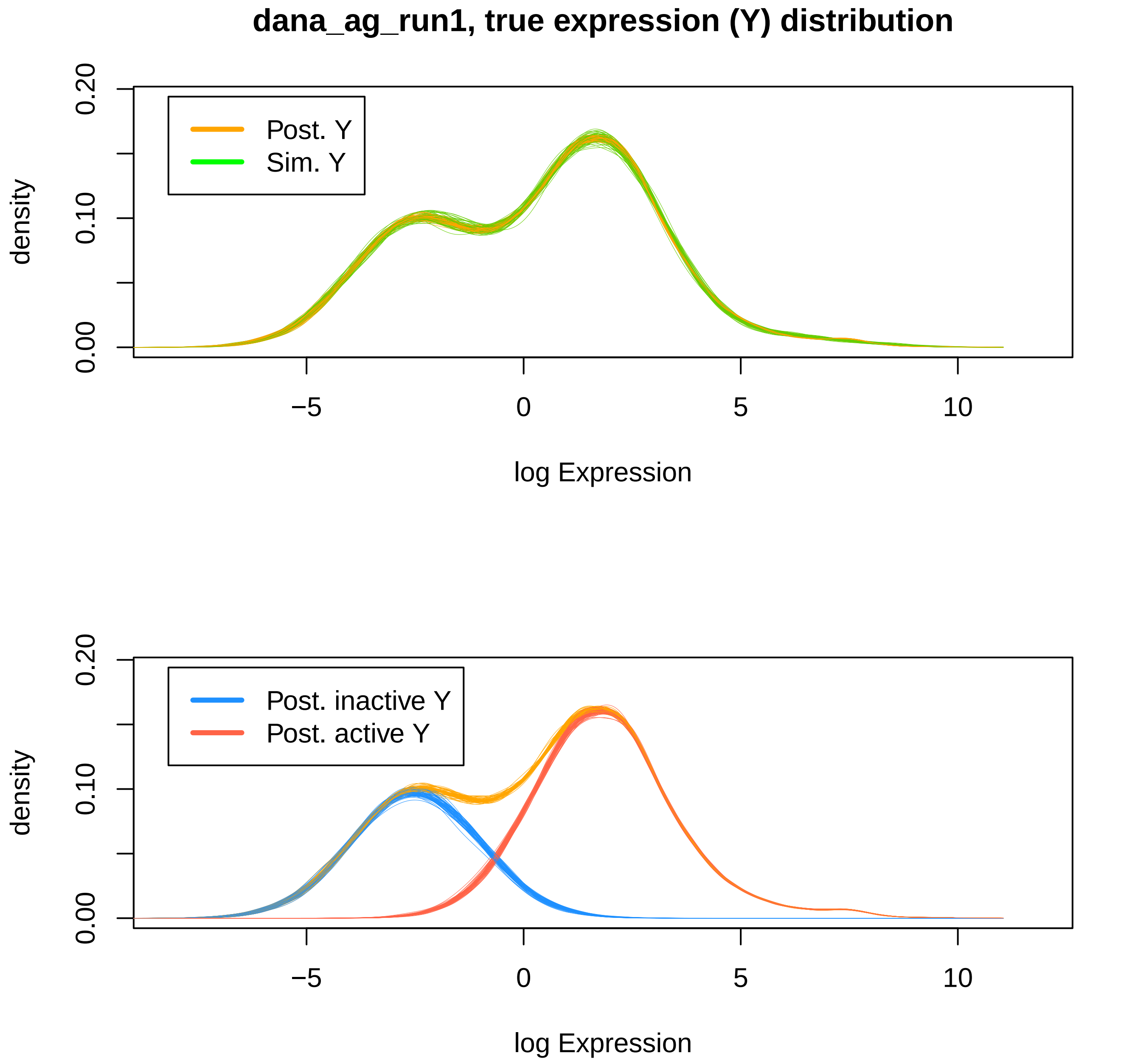

dana_ag_run1, true expression (Y) distribution
Post. Y
Sim. Y
density
0.00
0.10
0.20
−5
0
5
10
log Expression
Post. inactive Y
Post. active Y
density
0.00
0.10
0.20
−5
0
5
10
log Expression

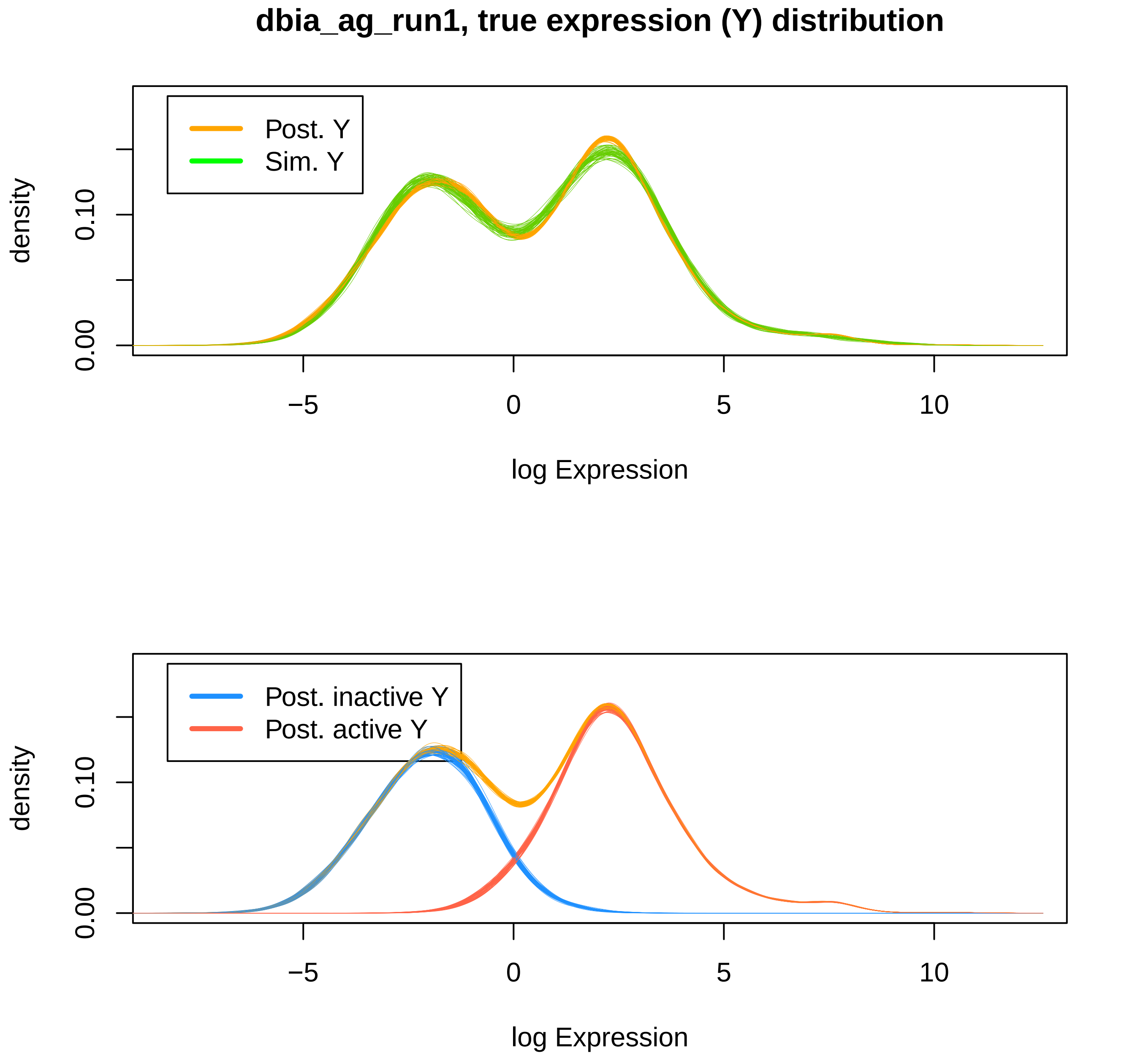
dbia_ag_run1, true expression (Y) distribution
Post. Y
Sim. Y
density
0.00
0.10
−5
0
5
10
log Expression
Post. inactive Y
Post. active Y
density
0.00
0.10
−5
0
5
10
log Expression

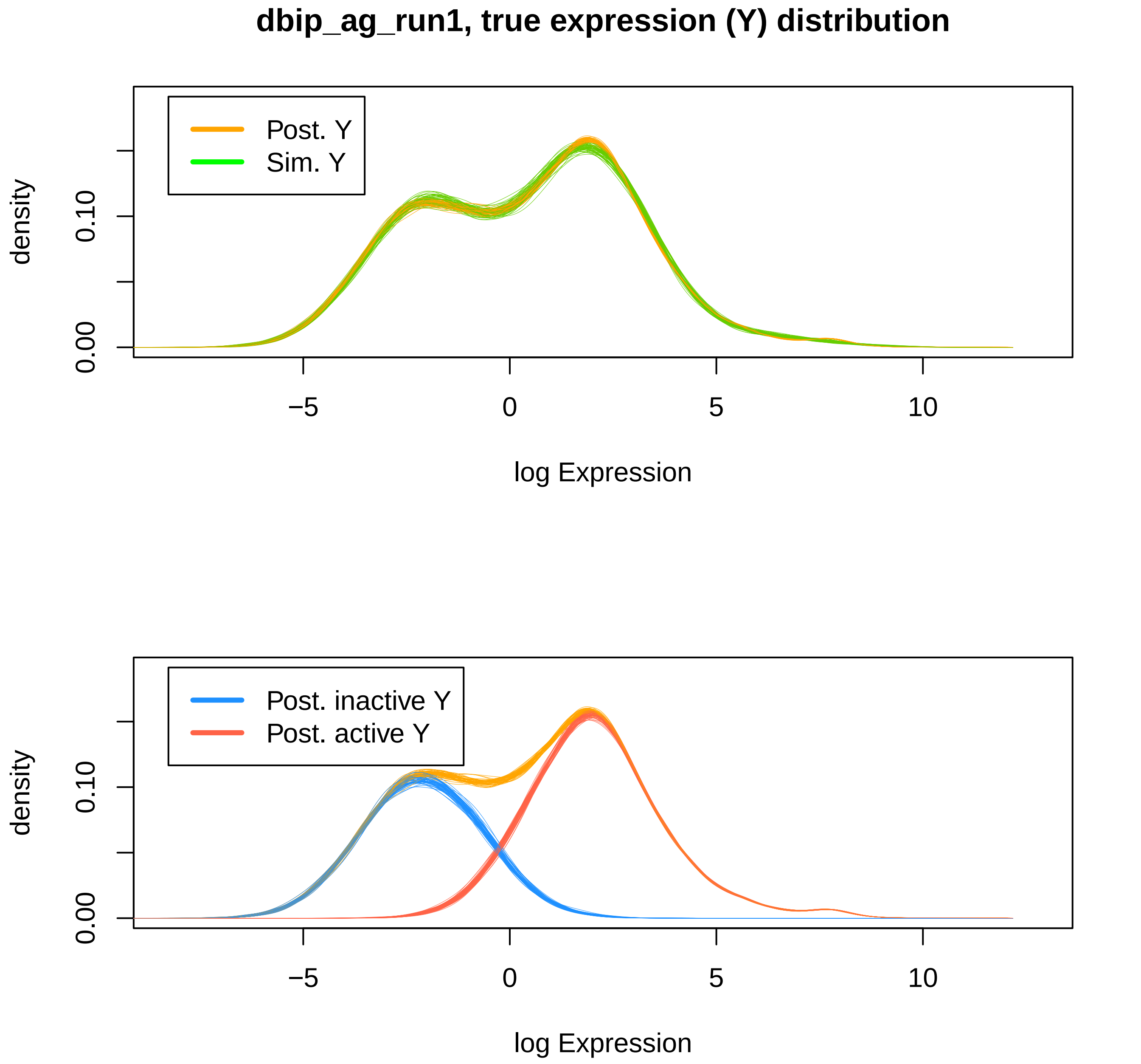
dbip_ag_run1, true expression (Y) distribution
Post. Y
Sim. Y
density
0.00
0.10
−5
0
5
10
log Expression
Post. inactive Y
Post. active Y
density
0.00
0.10
−5
0
5
10
log Expression

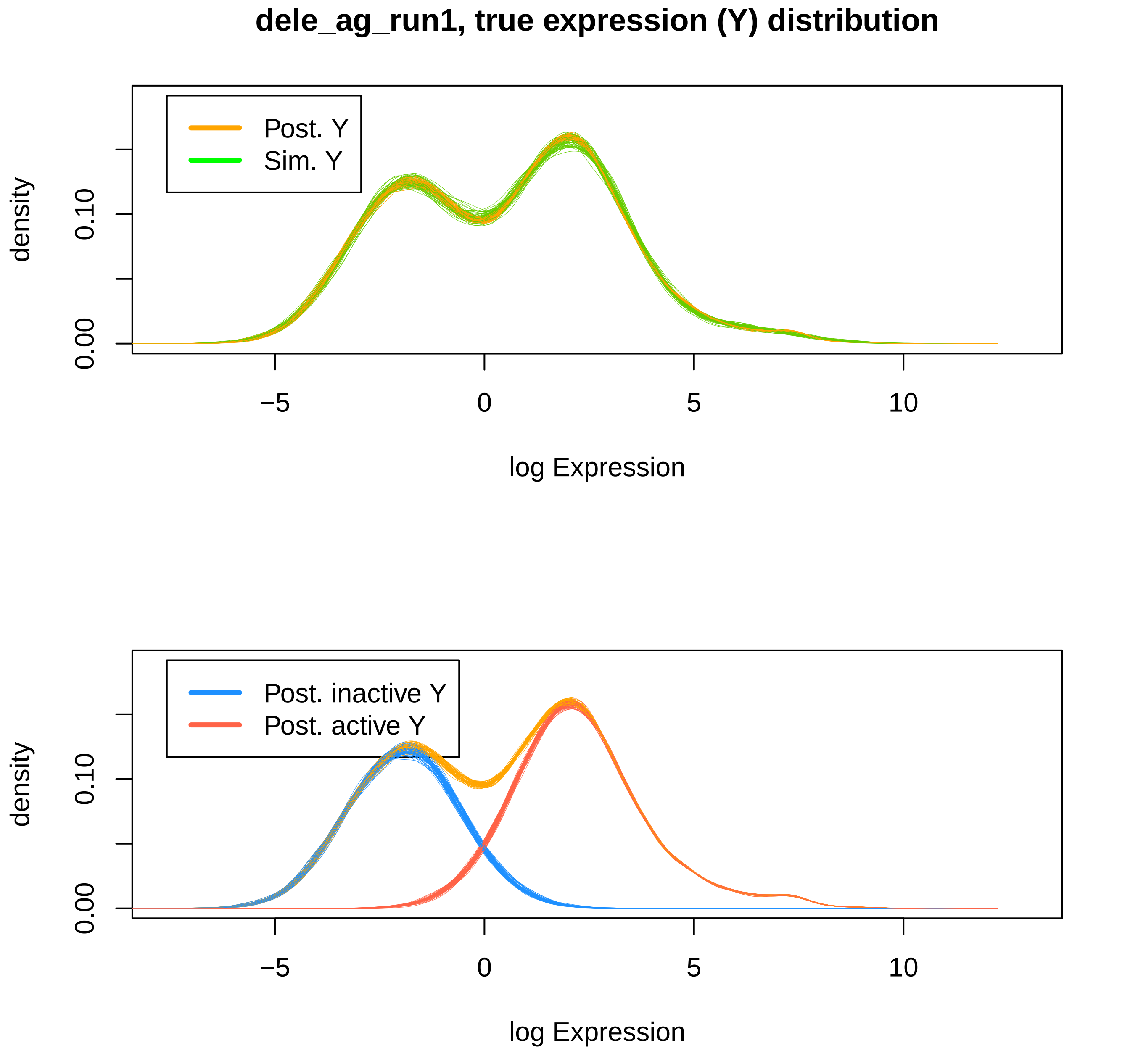
dele_ag_run1, true expression (Y) distribution
Post. Y
Sim. Y
density
0.00
0.10
−5
0
5
10
log Expression
Post. inactive Y
Post. active Y
density
0.00
0.10
−5
0
5
10
log Expression

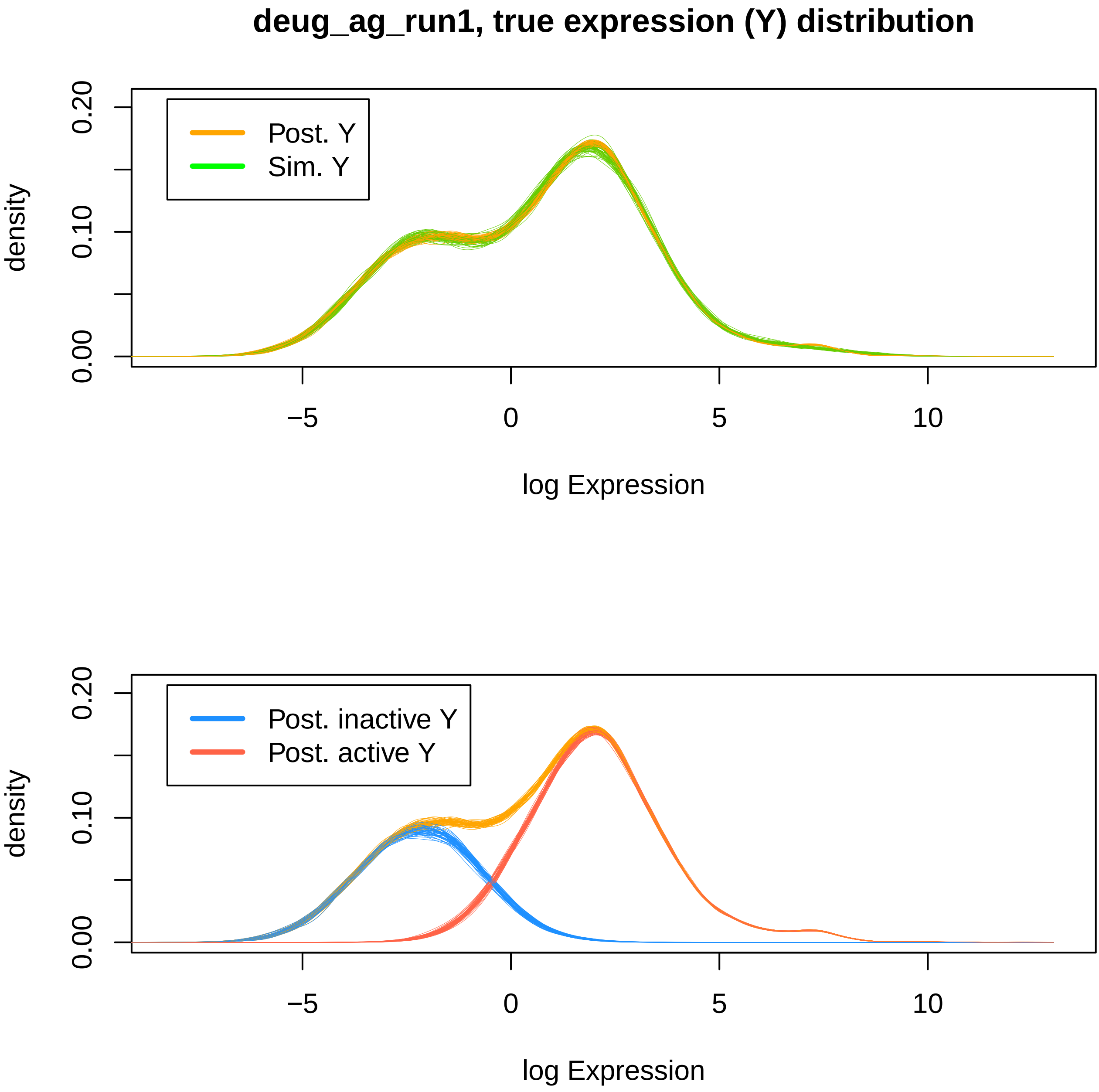

deug_ag_run1, true expression (Y) distribution
Post. Y
Sim. Y
density
0.00
0.10
0.20
−5
0
5
10
log Expression
Post. inactive Y
Post. active Y
density
0.00
0.10
0.20
−5
0
5
10
log Expression

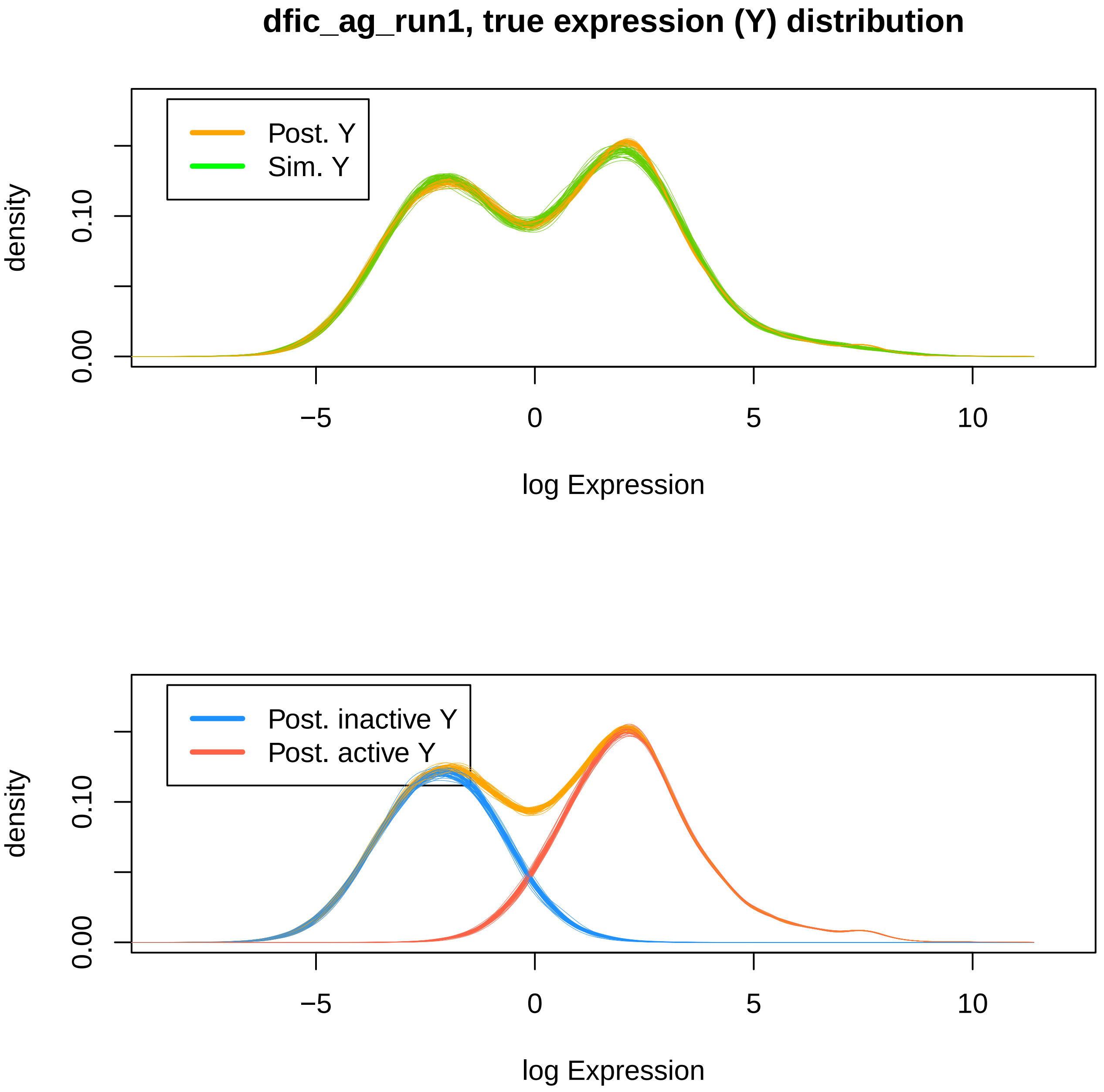
dfic_ag_run1, true expression (Y) distribution
Post. Y
Sim. Y
density
0.00
0.10
−5
0
5
10
log Expression
Post. inactive Y
Post. active Y
density
0.00
0.10
−5
0
5
10
log Expression

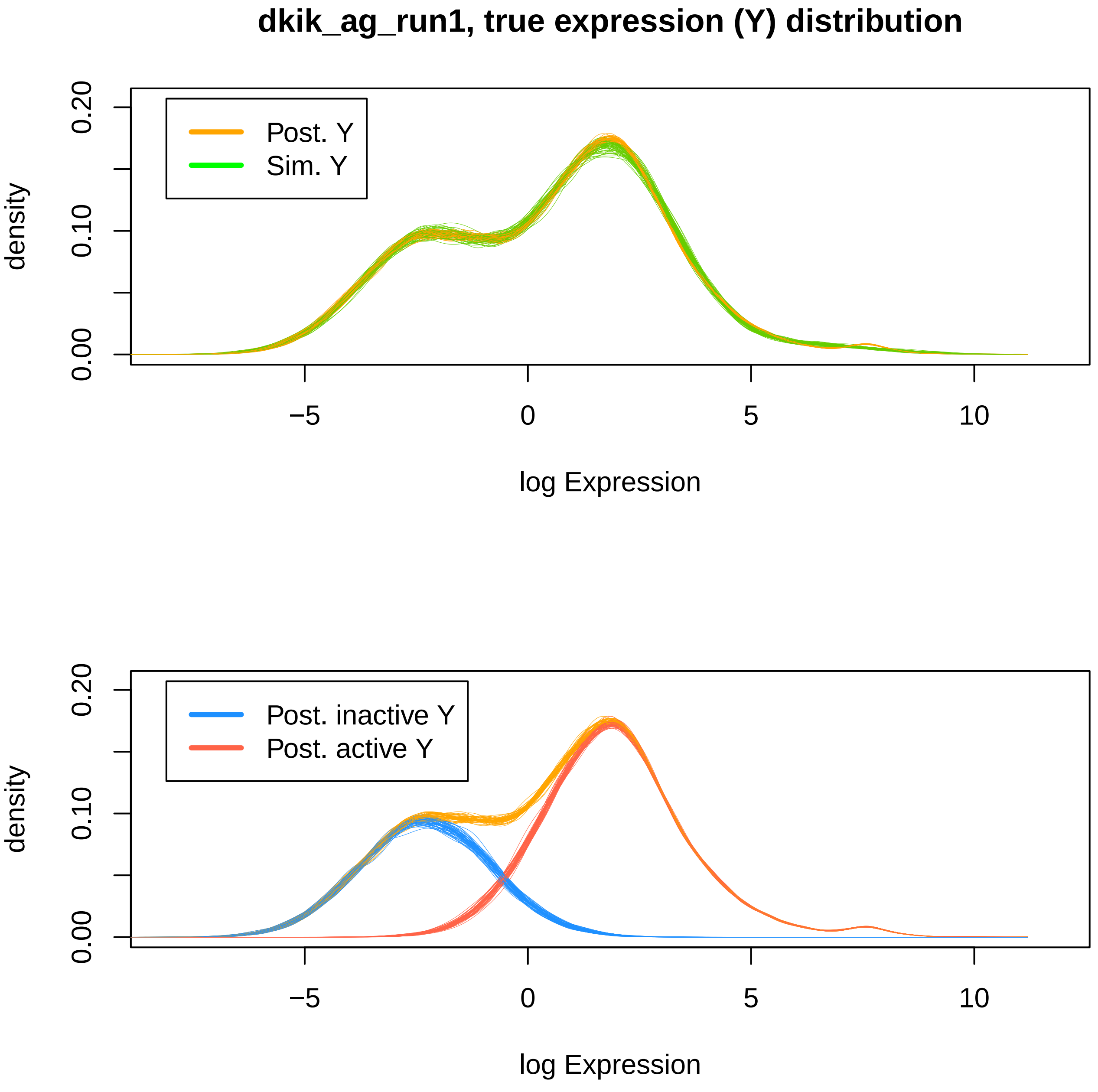
dkik_ag_run1, true expression (Y) distribution
Post. Y
Sim. Y
density
0.00
0.10
0.20
−5
0
5
10
log Expression
Post. inactive Y
Post. active Y
density
0.00
0.10
0.20
−5
0
5
10
log Expression

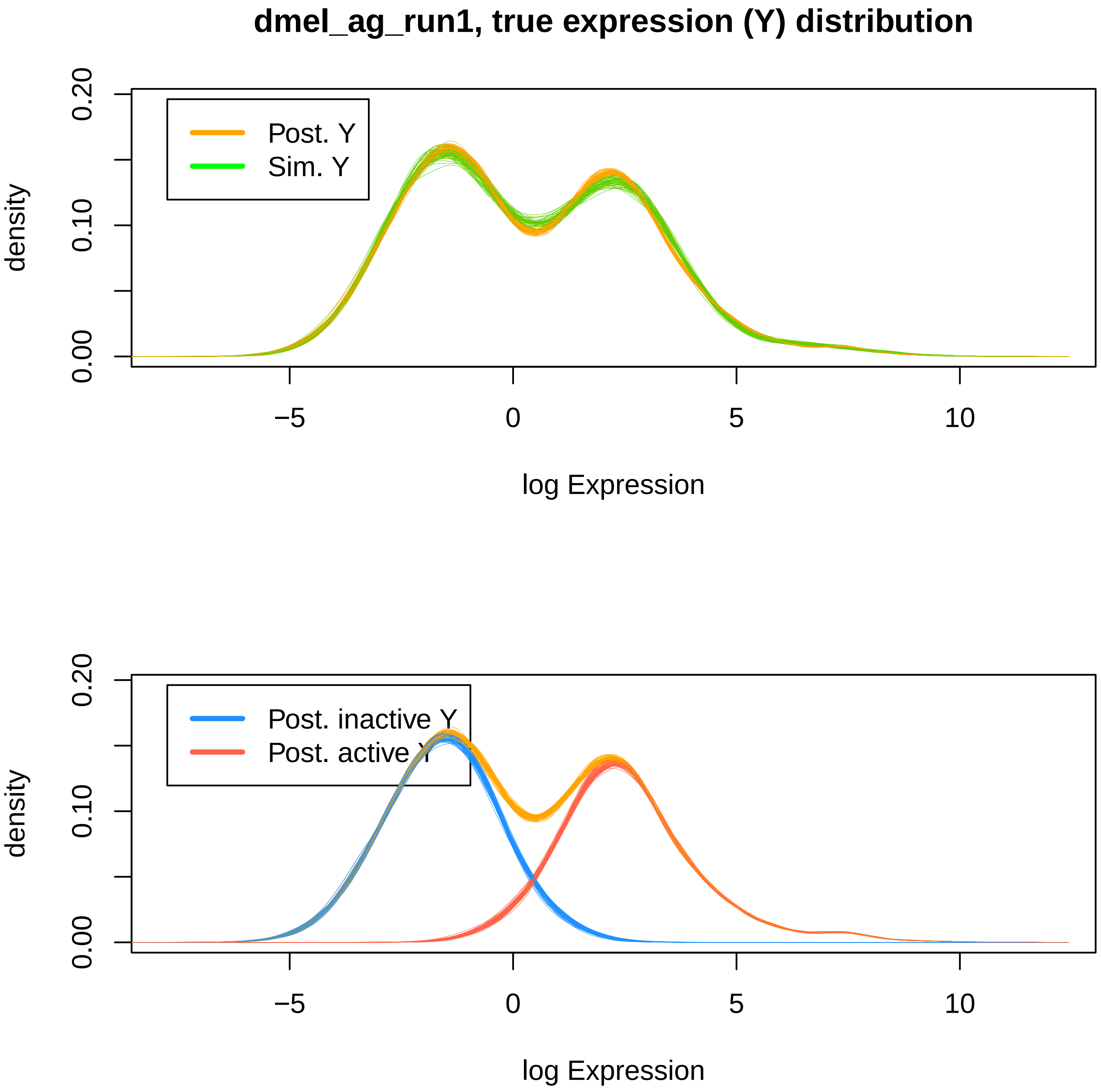
dmel_ag_run1, true expression (Y) distribution
Post. Y
Sim. Y
density
0.00
0.10
0.20
−5
0
5
10
log Expression
Post. inactive Y
Post. active Y
density
0.00
0.10
0.20
−5
0
5
10
log Expression

dsim_ag_run1, true expression (Y) distribution

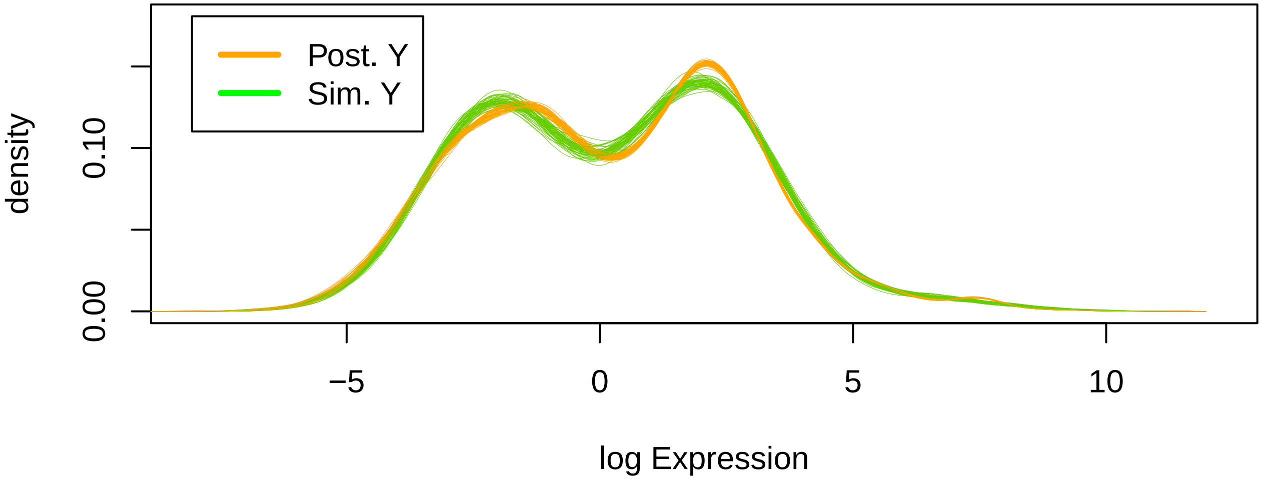


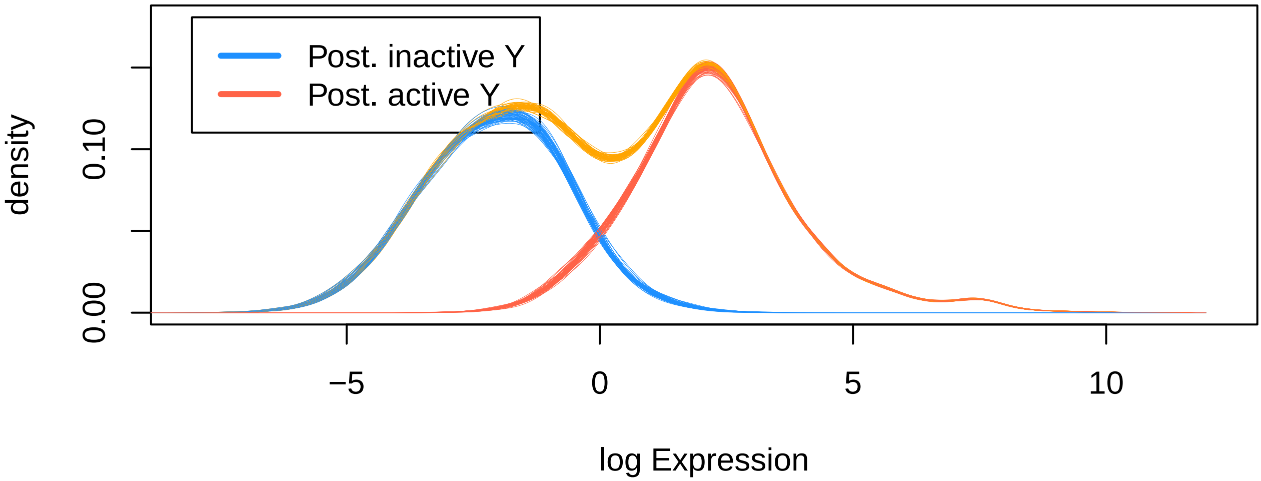

**dtak_ag_run1, true expression (Y) distribution**

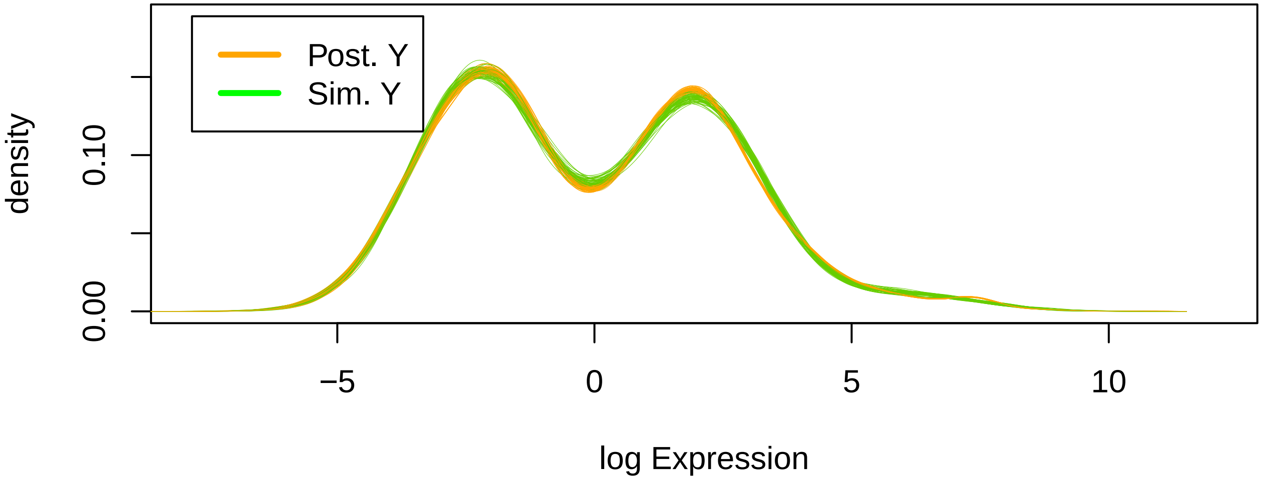


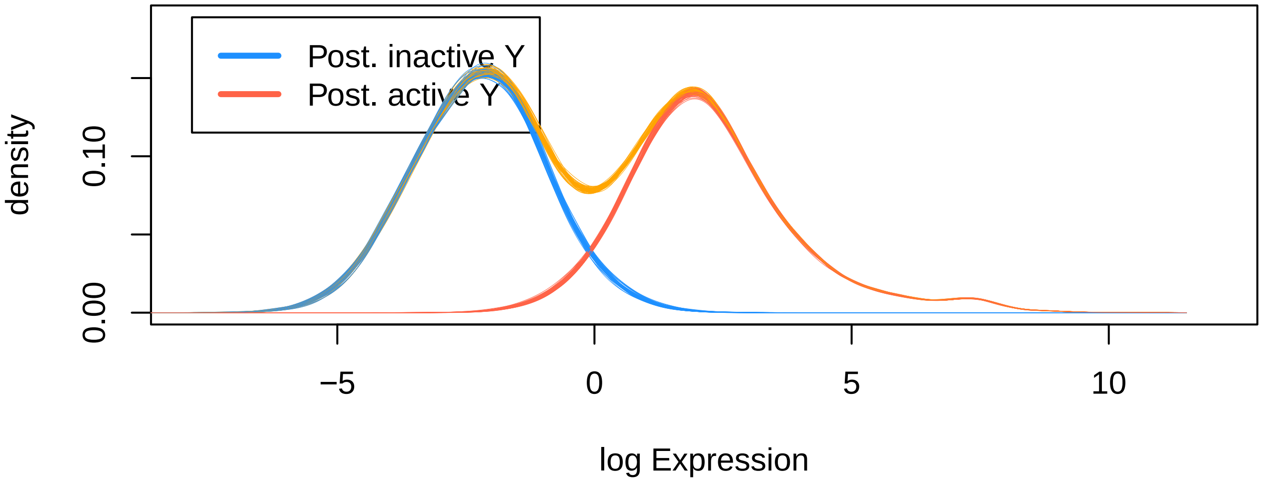

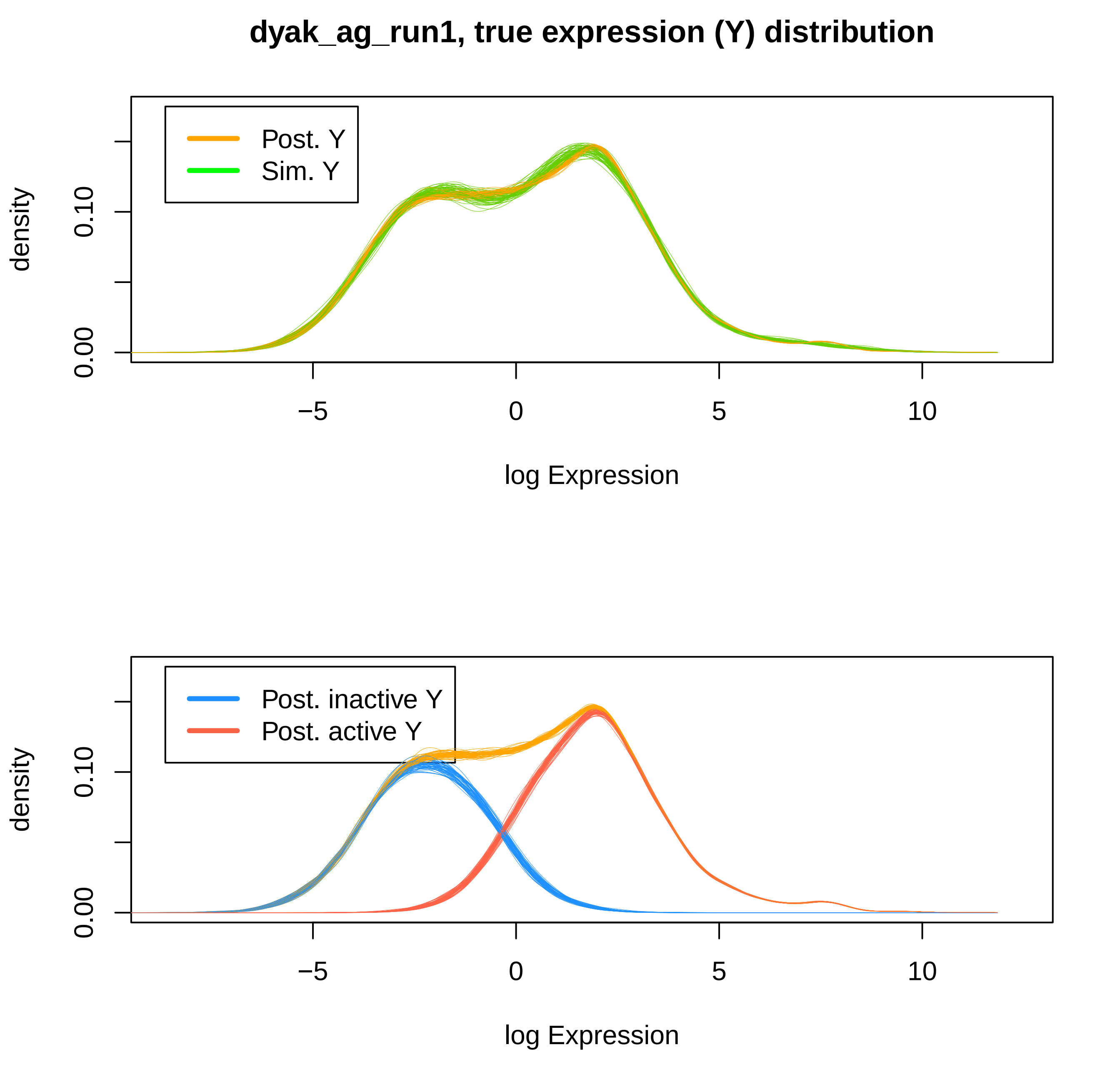

dyak_ag_run1, true expression (Y) distribution
Post. Y
Sim. Y
density
0.00
0.10
−5
0
5
10
log Expression
Post. inactive Y
Post. active Y
density
0.00
0.10
−5
0
5
10
log Expression

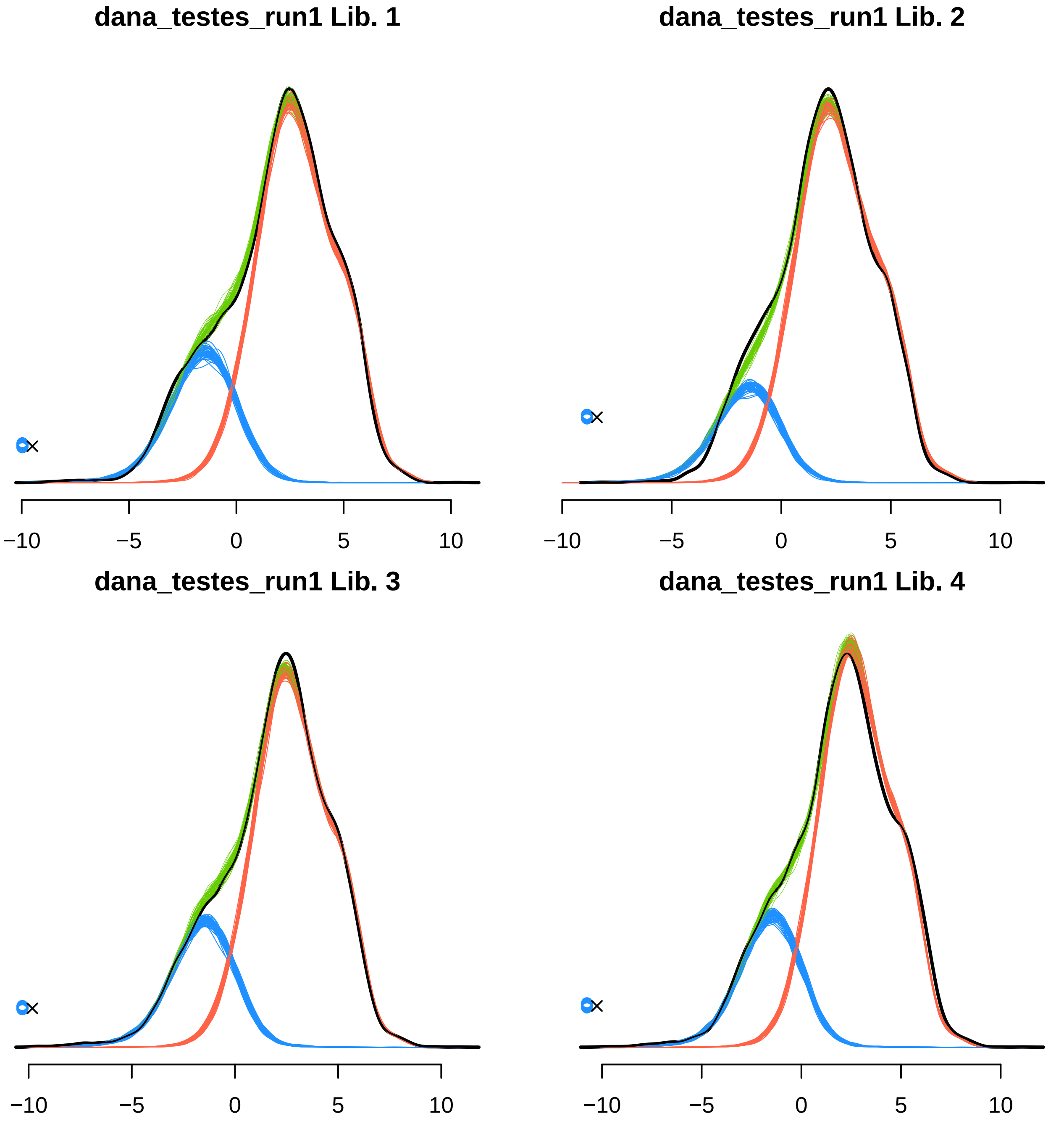

dana_testes_run1 Lib. 1
−10
−5
0
5
10
dana_testes_run1 Lib. 2
−10
−5
0
5
10
dana_testes_run1 Lib. 3
−10
−5
0
5
10
dana_testes_run1 Lib. 4
−10
−5
0
5
10

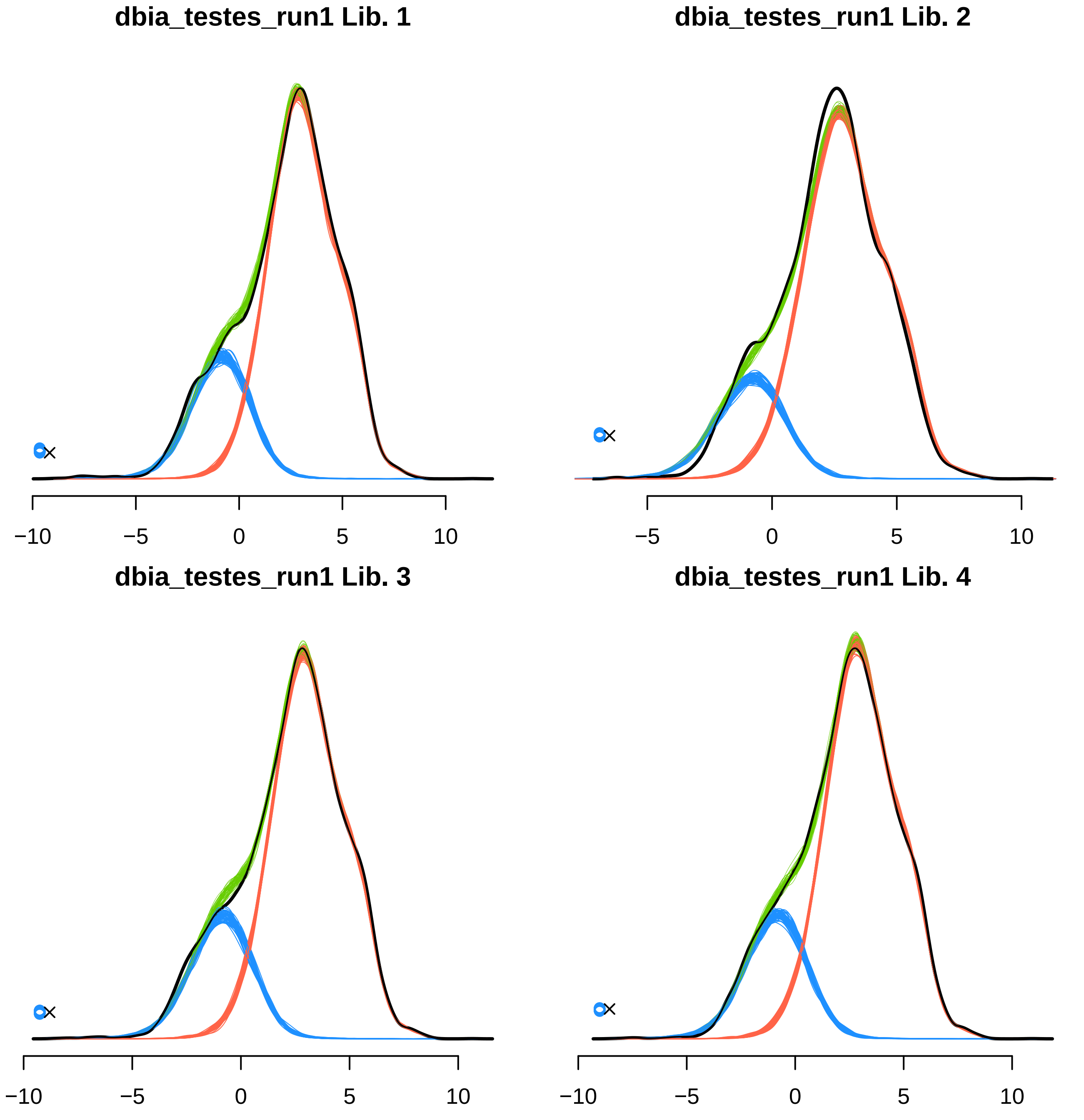
dbia_testes_run1 Lib. 1
−10
−5
0
5
10
dbia_testes_run1 Lib. 2
−5
0
5
10
dbia_testes_run1 Lib. 3
−10
−5
0
5
10
dbia_testes_run1 Lib. 4
−10
−5
0
5
10

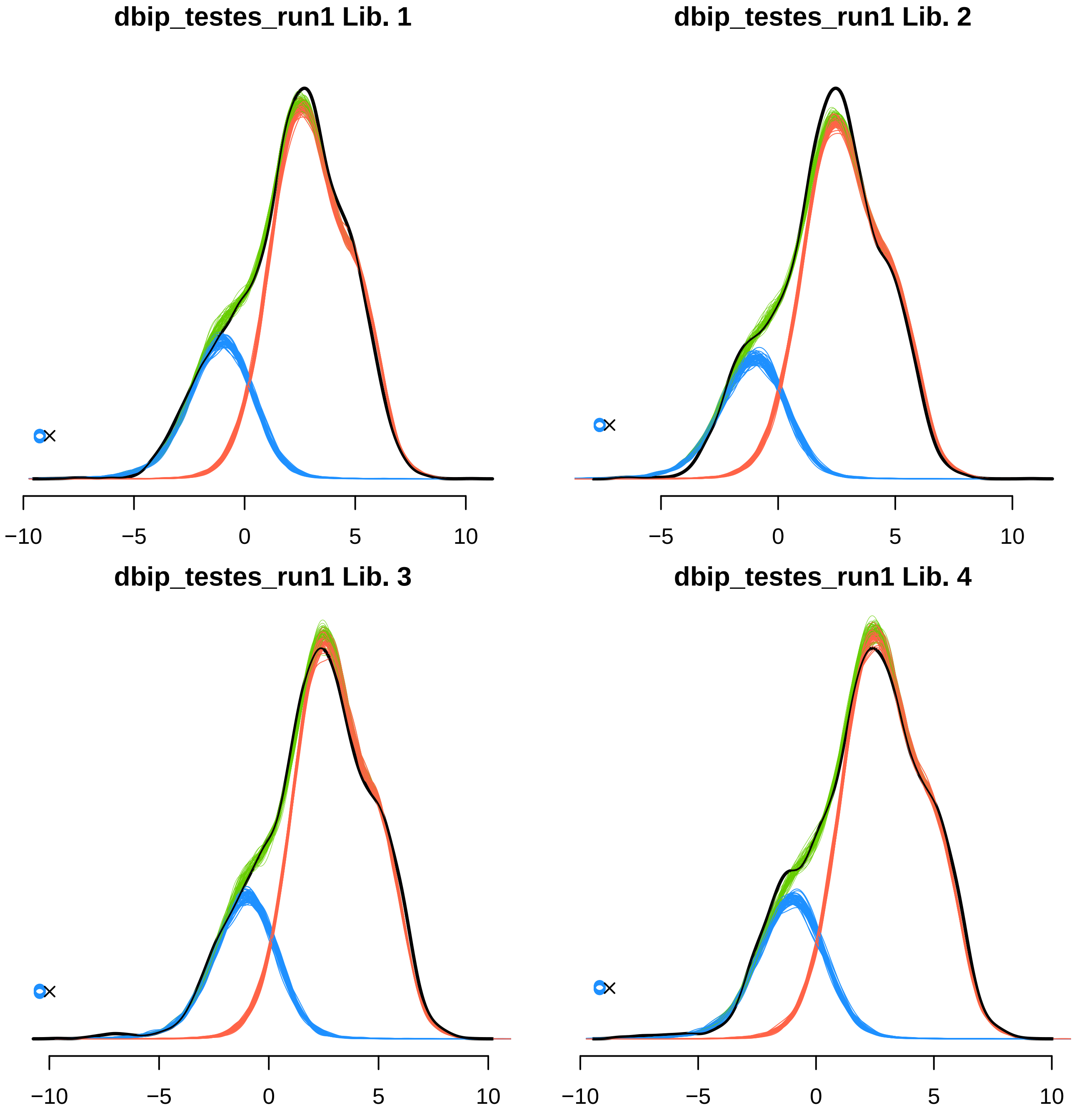
dbip_testes_run1 Lib. 1
−10
−5
0
5
10
dbip_testes_run1 Lib. 2
−5
0
5
10
dbip_testes_run1 Lib. 3
−10
−5
0
5
10
dbip_testes_run1 Lib. 4
−10
−5
0
5
10

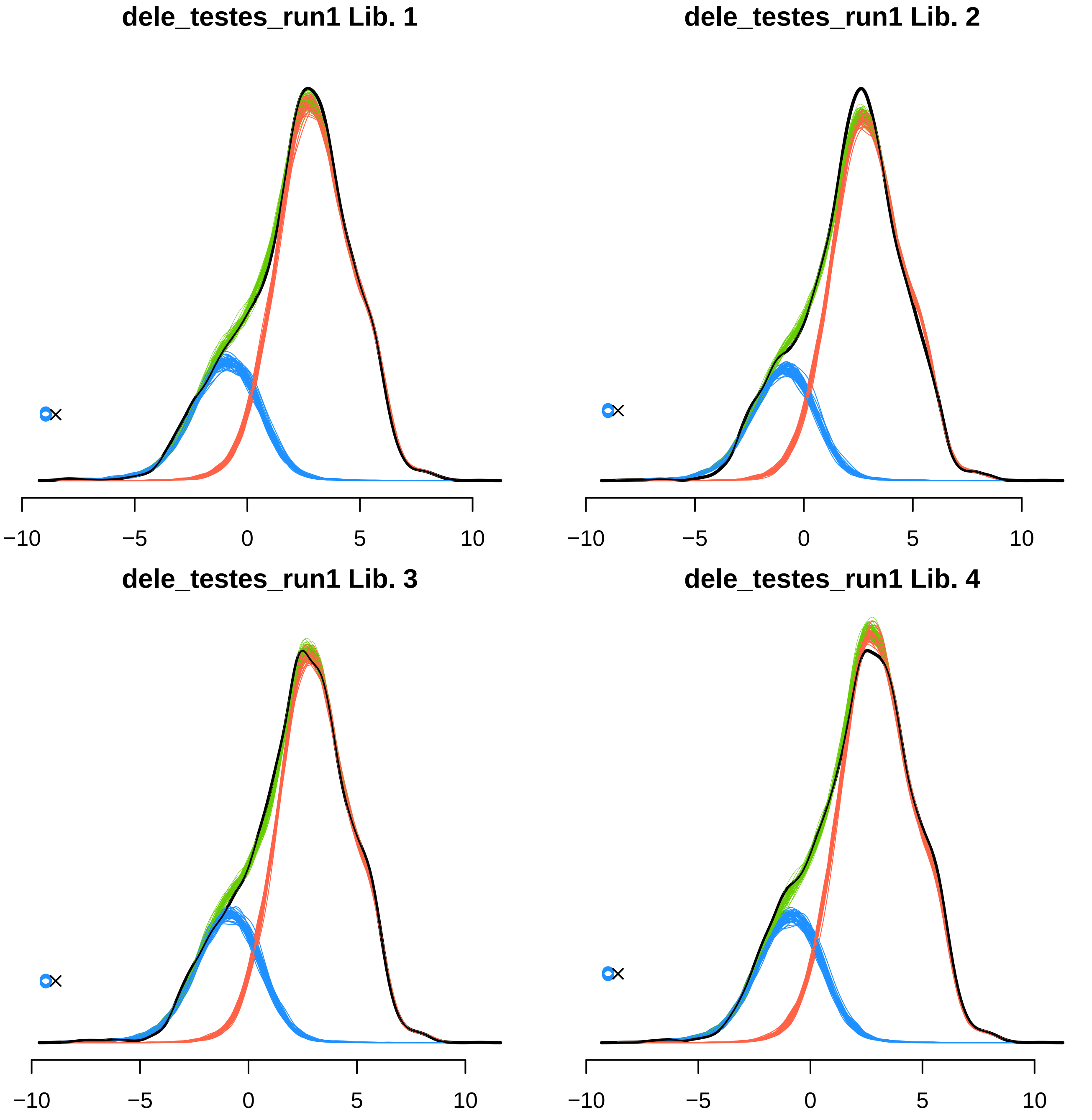
dele_testes_run1 Lib. 1
dele_testes_run1 Lib. 2
dele_testes_run1 Lib. 3
dele_testes_run1 Lib. 4
−10
−5
0
5
10

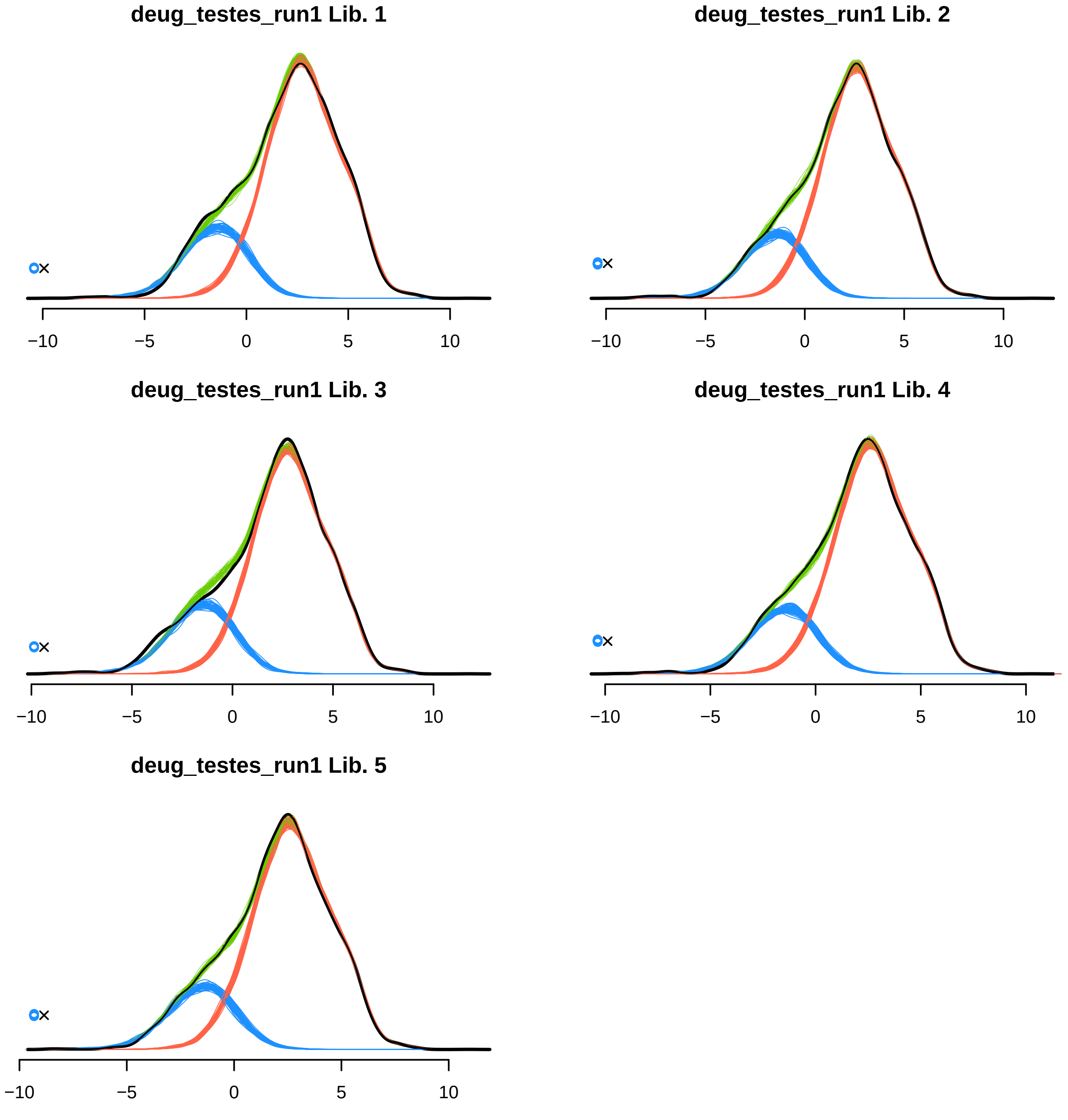
deug_testes_run1 Lib. 1
deug_testes_run1 Lib. 2
deug_testes_run1 Lib. 3
deug_testes_run1 Lib. 4
deug_testes_run1 Lib. 5
−10
−5
0
5
10

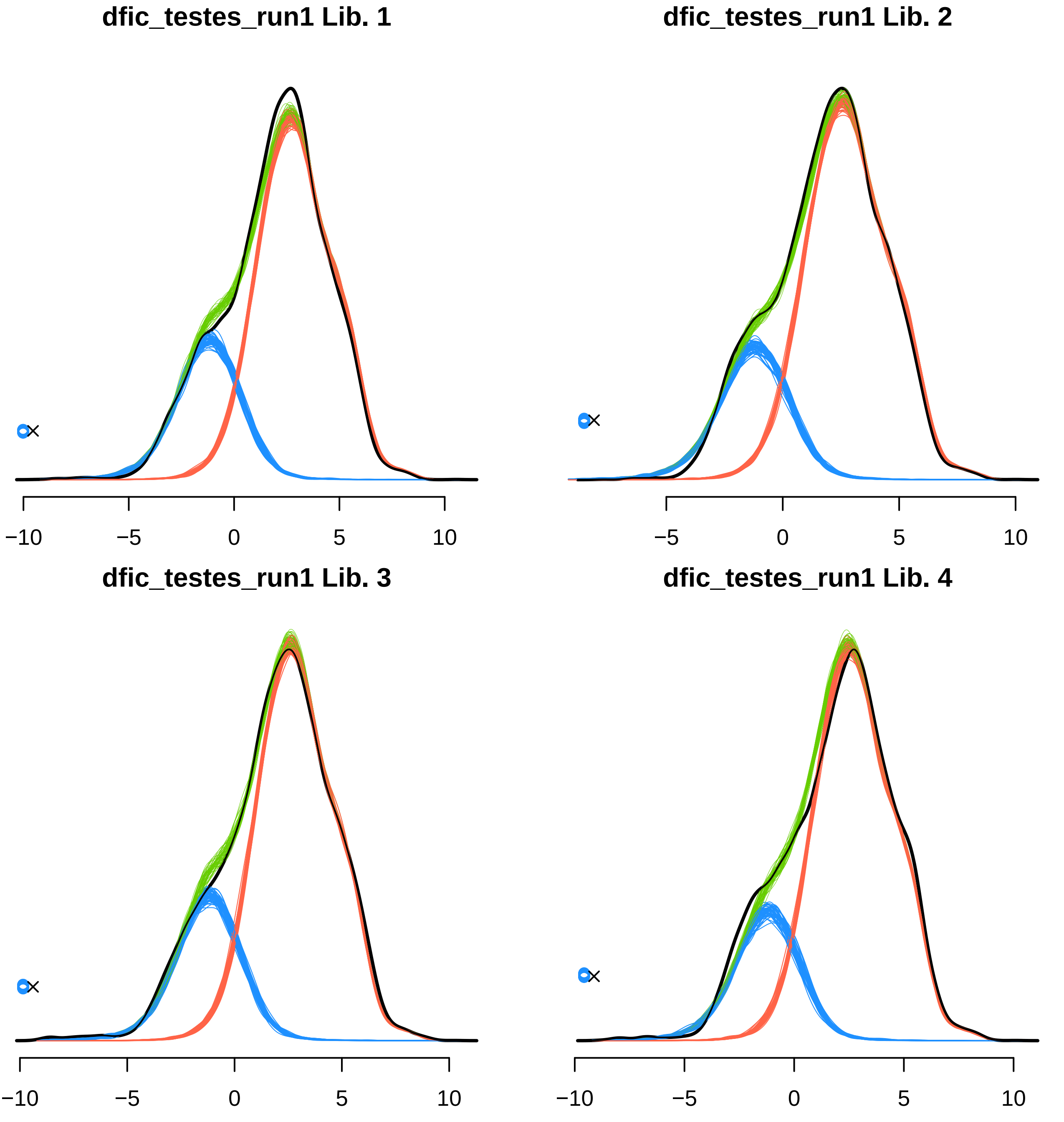
dfic_testes_run1 Lib. 1
−10
−5
0
5
10
dfic_testes_run1 Lib. 2
−5
0
5
10
dfic_testes_run1 Lib. 3
−10
−5
0
5
10
dfic_testes_run1 Lib. 4
−10
−5
0
5
10

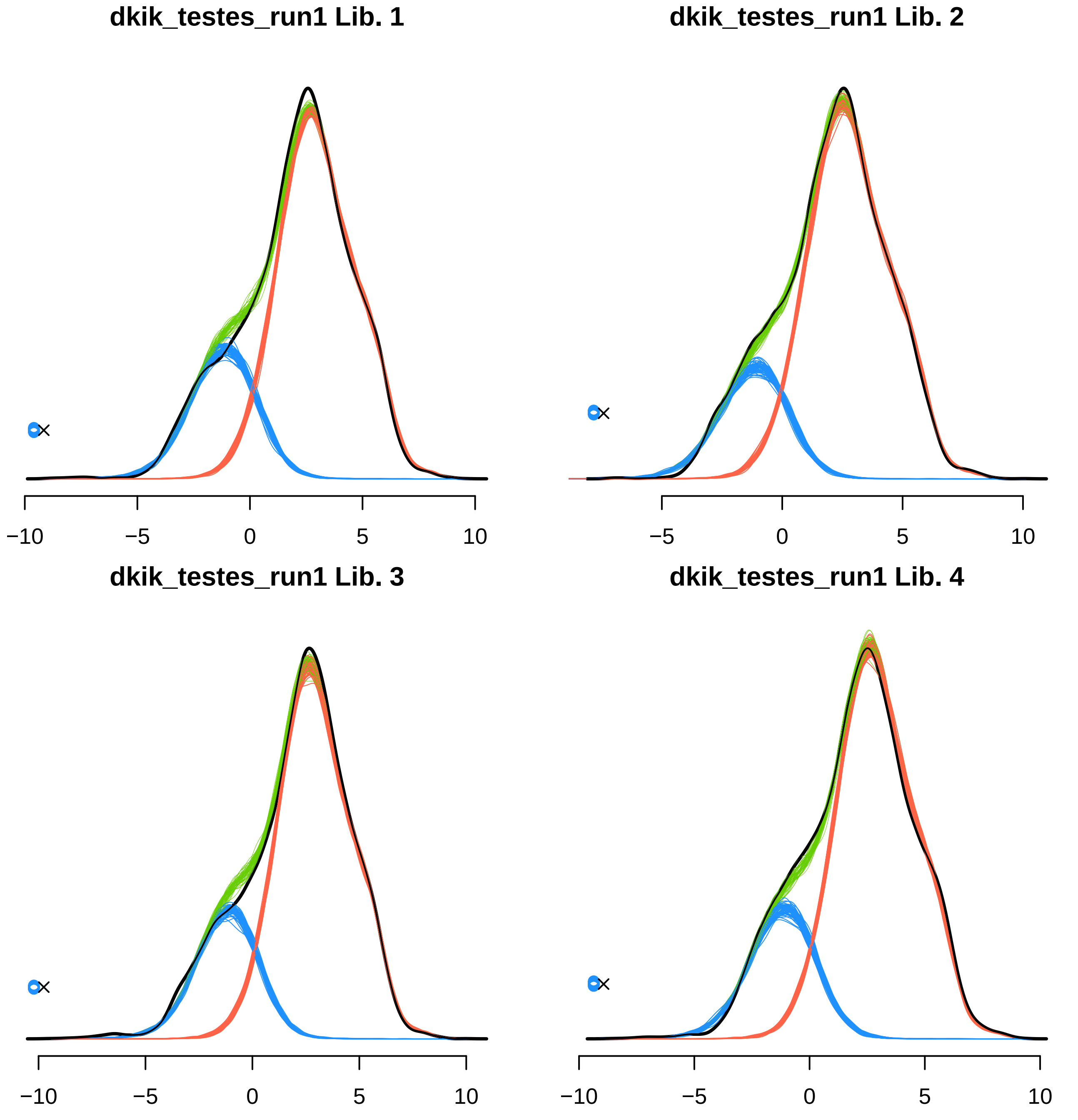

dkik_testes_run1 Lib. 1
−10
−5
0
5
10
dkik_testes_run1 Lib. 2
−5
0
5
10
dkik_testes_run1 Lib. 3
−10
−5
0
5
10
dkik_testes_run1 Lib. 4
−10
−5
0
5
10

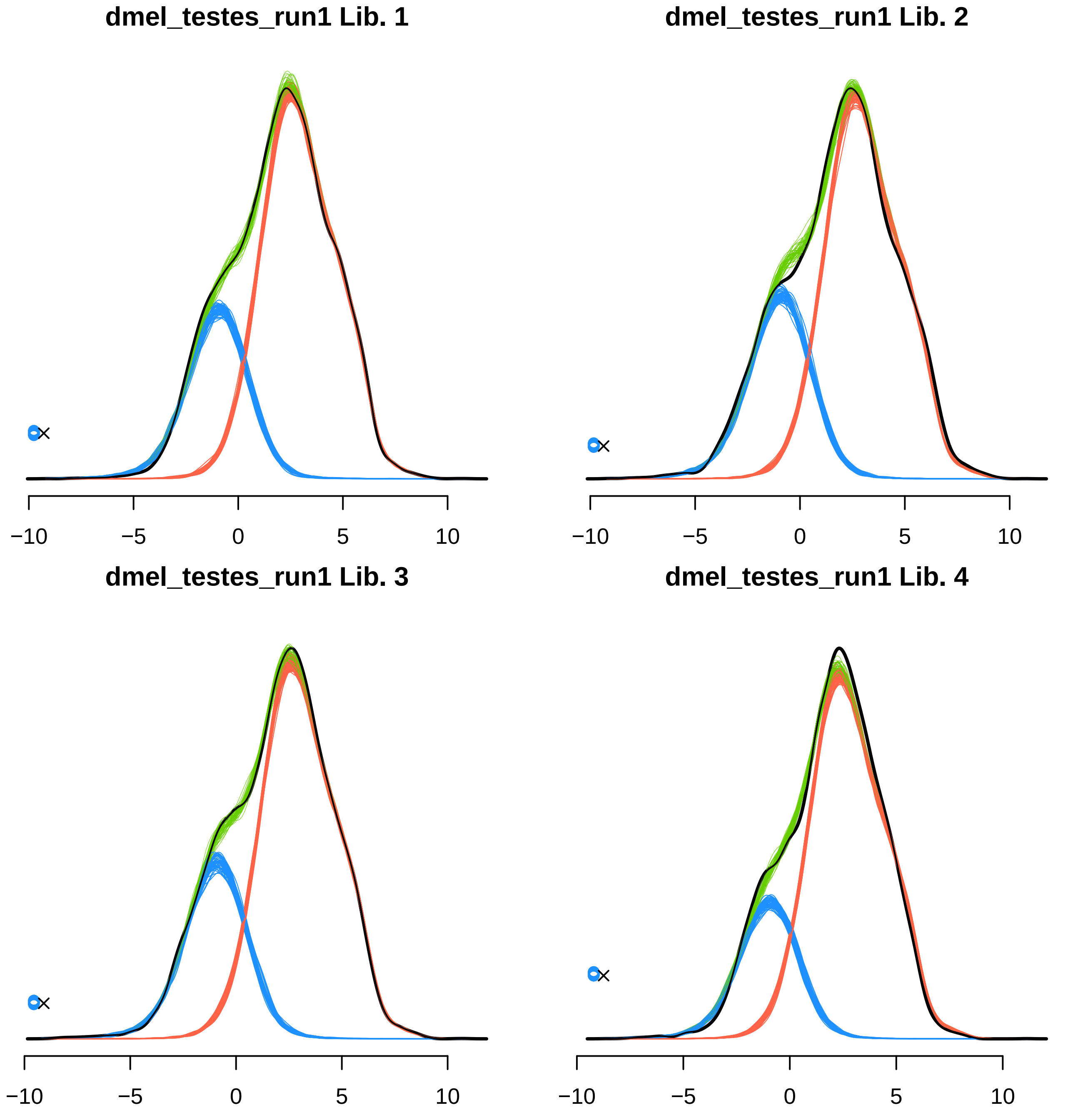

dmel_testes_run1 Lib. 1
dmel_testes_run1 Lib. 2
dmel_testes_run1 Lib. 3
dmel_testes_run1 Lib. 4
−10
−5
0
5
10

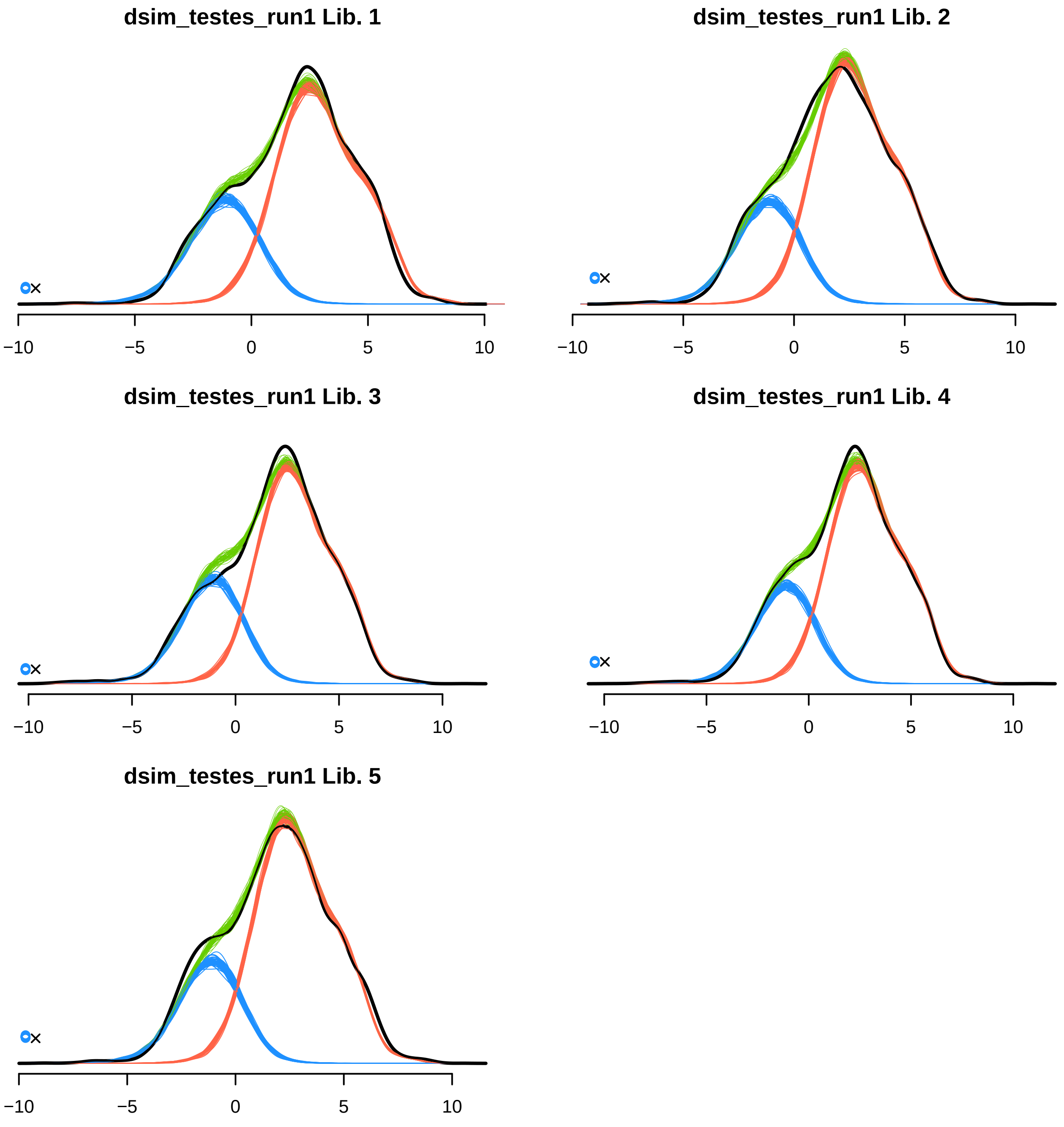
dsim_testes_run1 Lib. 1
−10
−5
0
5
10
dsim_testes_run1 Lib. 2
−10
−5
0
5
10
dsim_testes_run1 Lib. 3
−10
−5
0
5
10
dsim_testes_run1 Lib. 4
−10
−5
0
5
10
dsim_testes_run1 Lib. 5
−10
−5
0
5
10

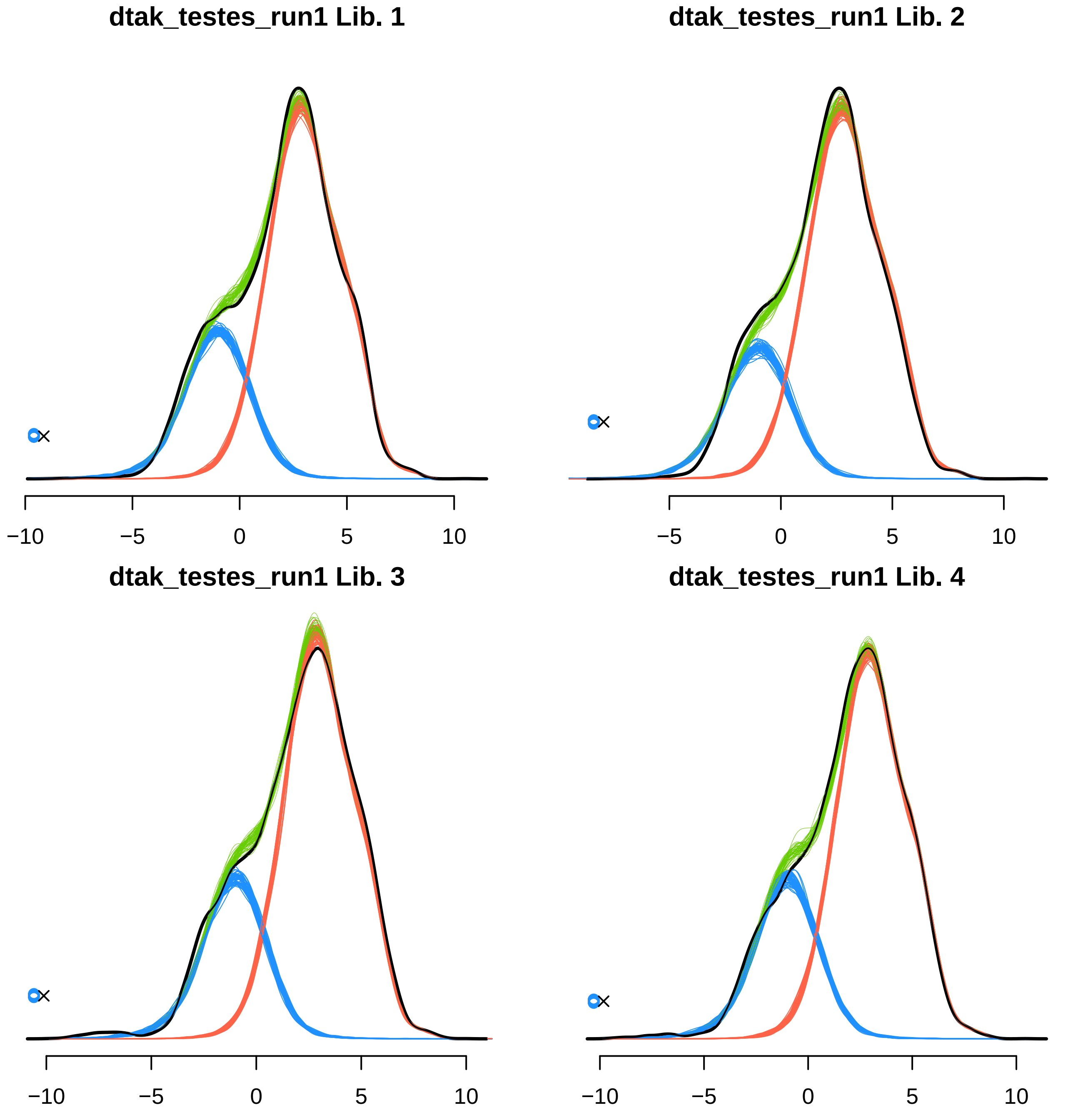

dtak_testes_run1 Lib. 1
−10
−5
0
5
10
dtak_testes_run1 Lib. 2
−5
0
5
10
dtak_testes_run1 Lib. 3
−10
−5
0
5
10
dtak_testes_run1 Lib. 4
−10
−5
0
5
10

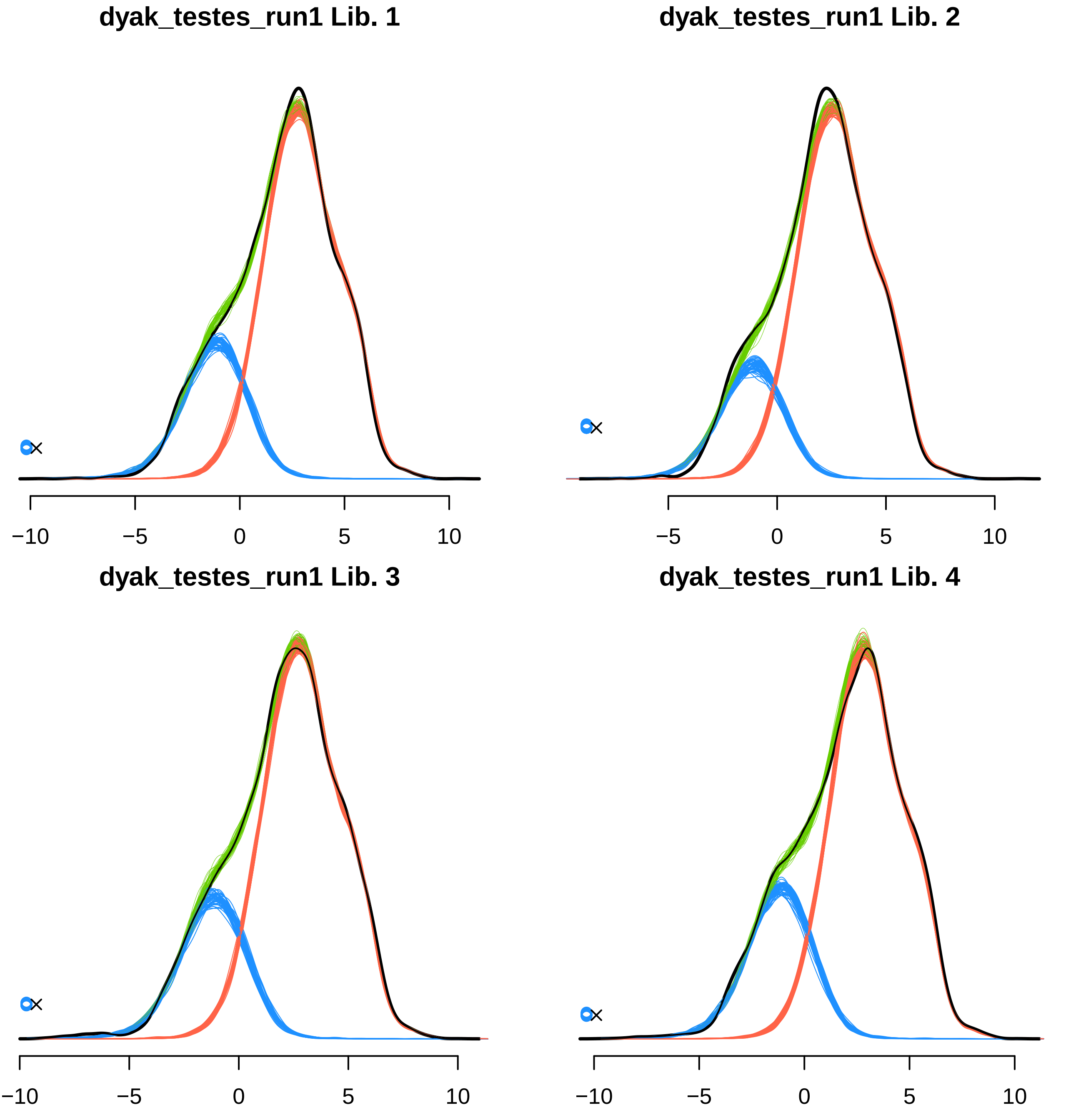
dyak_testes_run1 Lib. 1
−10
−5
0
5
10
dyak_testes_run1 Lib. 2
−5
0
5
10
dyak_testes_run1 Lib. 3
−10
−5
0
5
10
dyak_testes_run1 Lib. 4
−10
−5
0
5
10

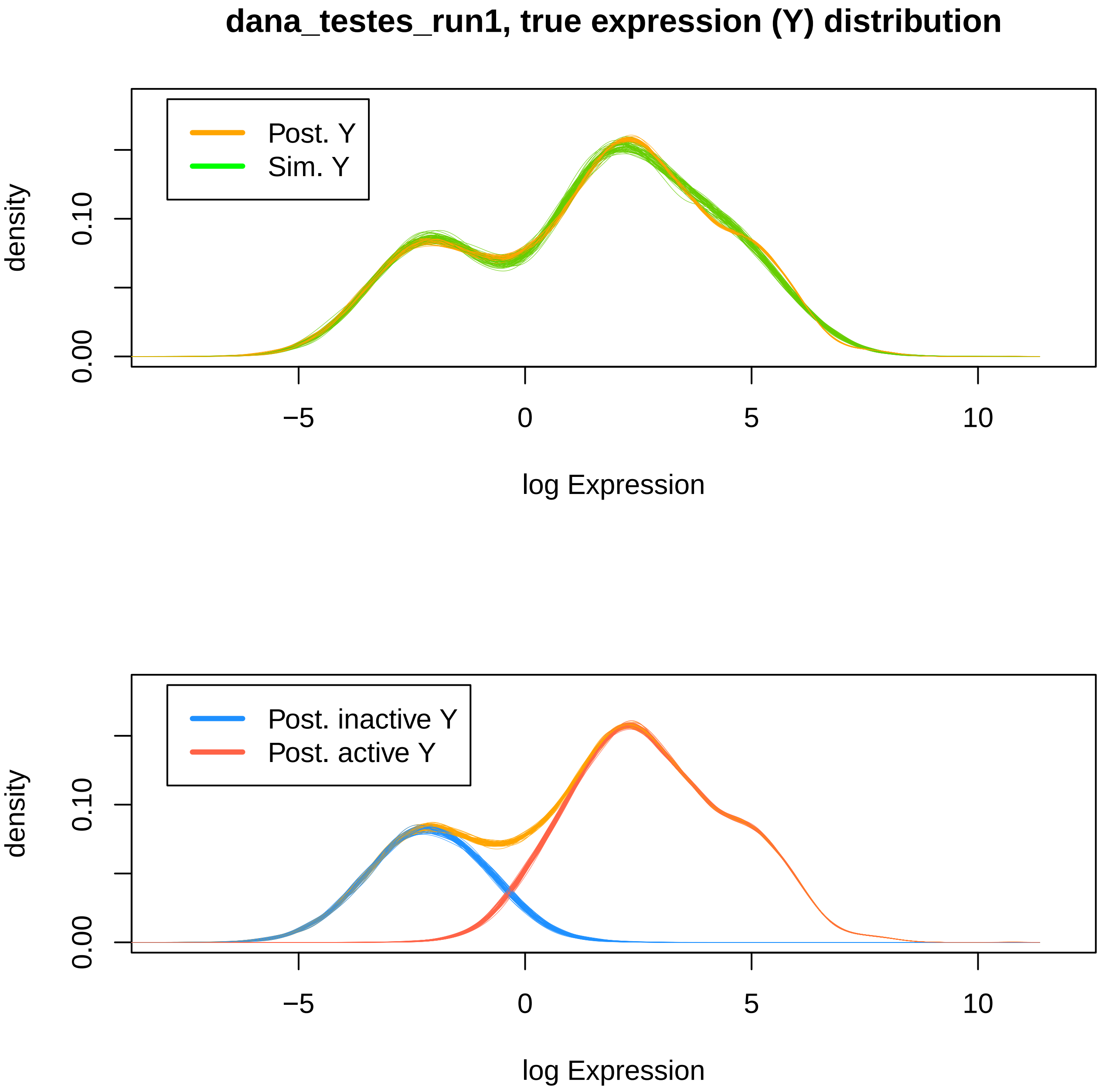
dana_testes_run1, true expression (Y) distribution
Post. Y
Sim. Y
density
0.00
0.10
−5
0
5
10
log Expression
Post. inactive Y
Post. active Y
density
0.00
0.10
−5
0
5
10
log Expression

**dbia_testes_run1, true expression (Y) distribution**

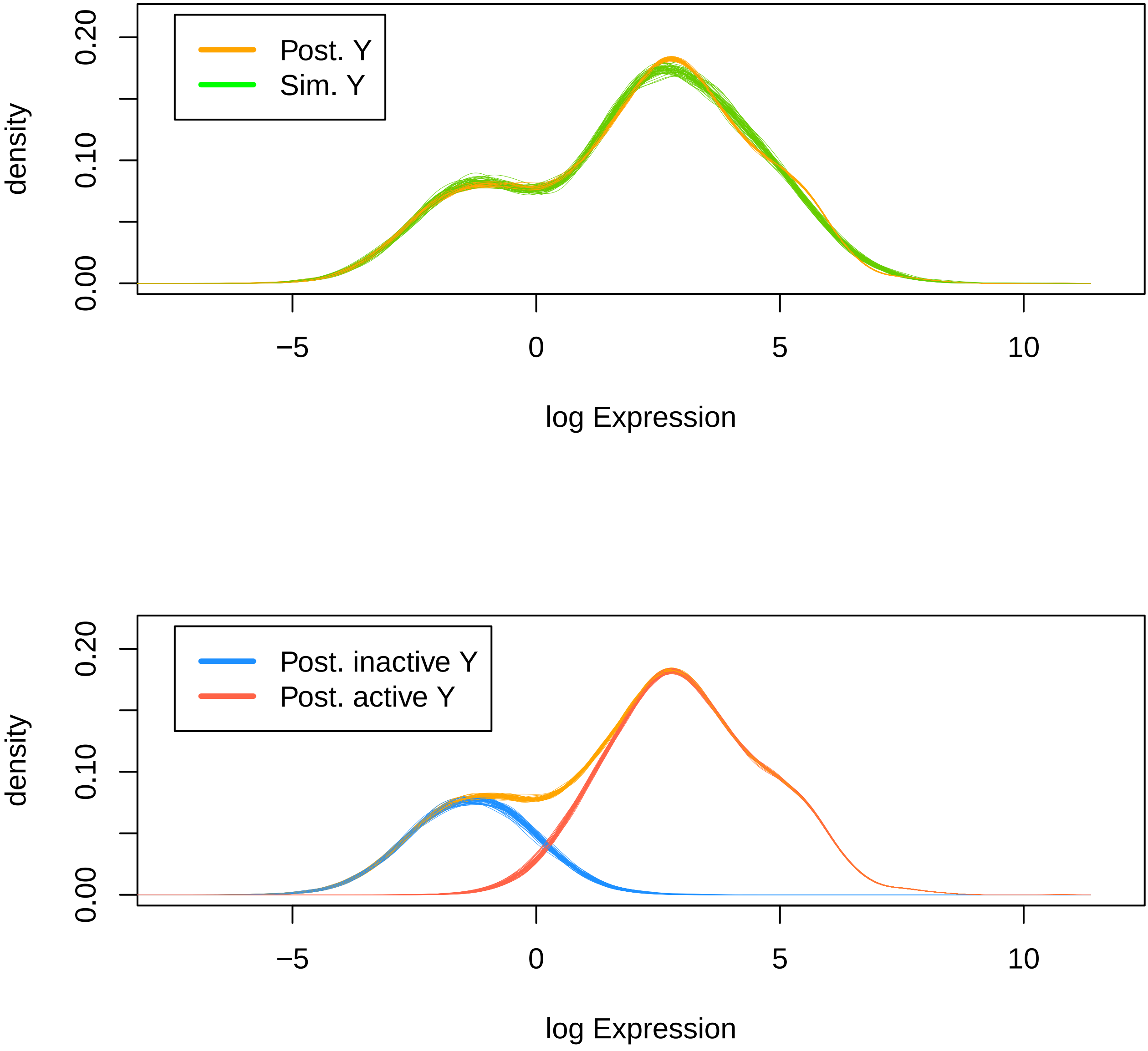

dbip_testes_run1, true expression (Y) distribution

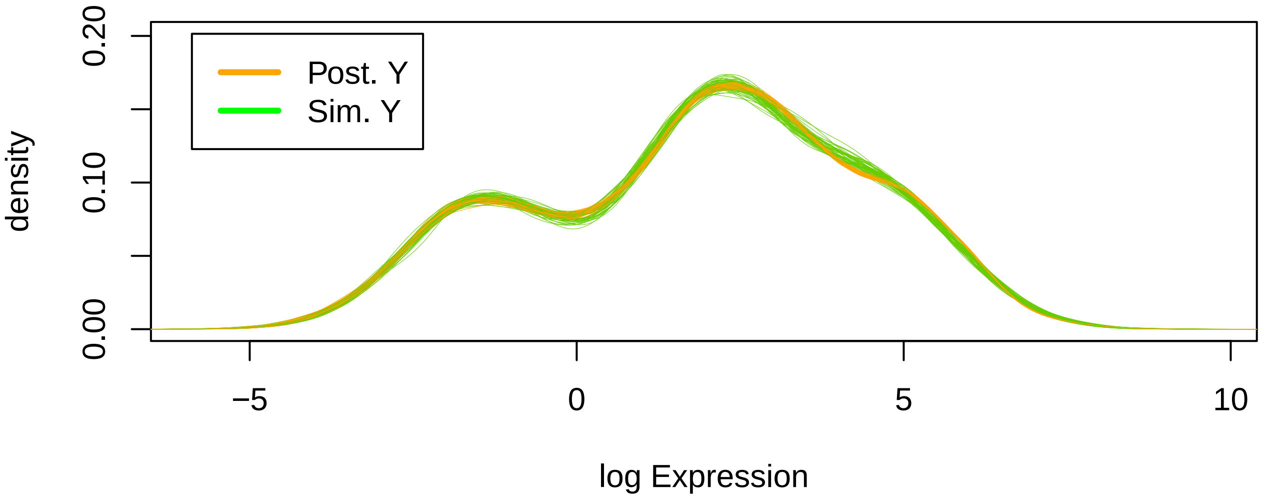


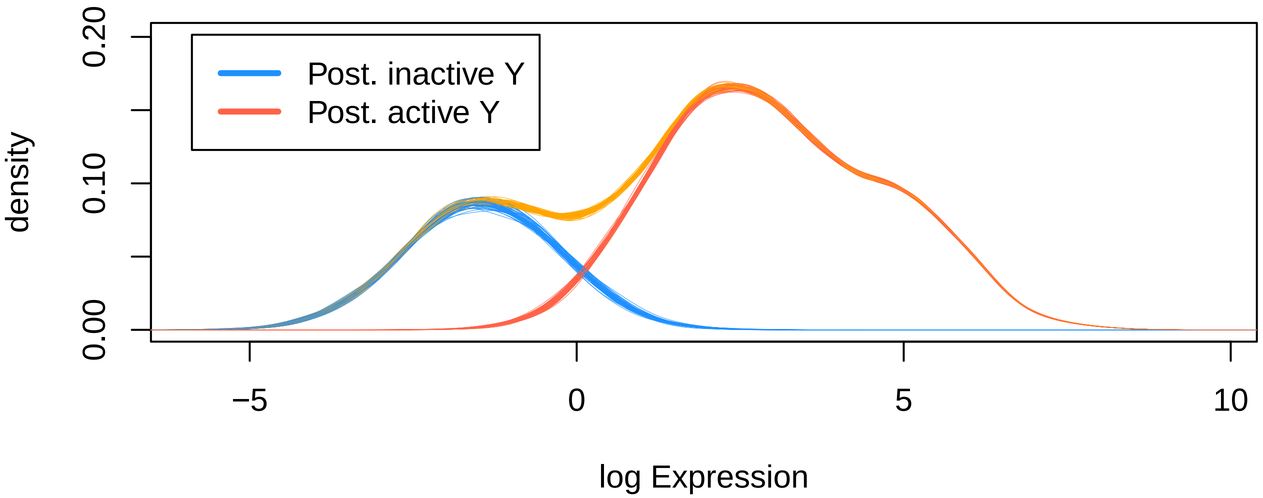

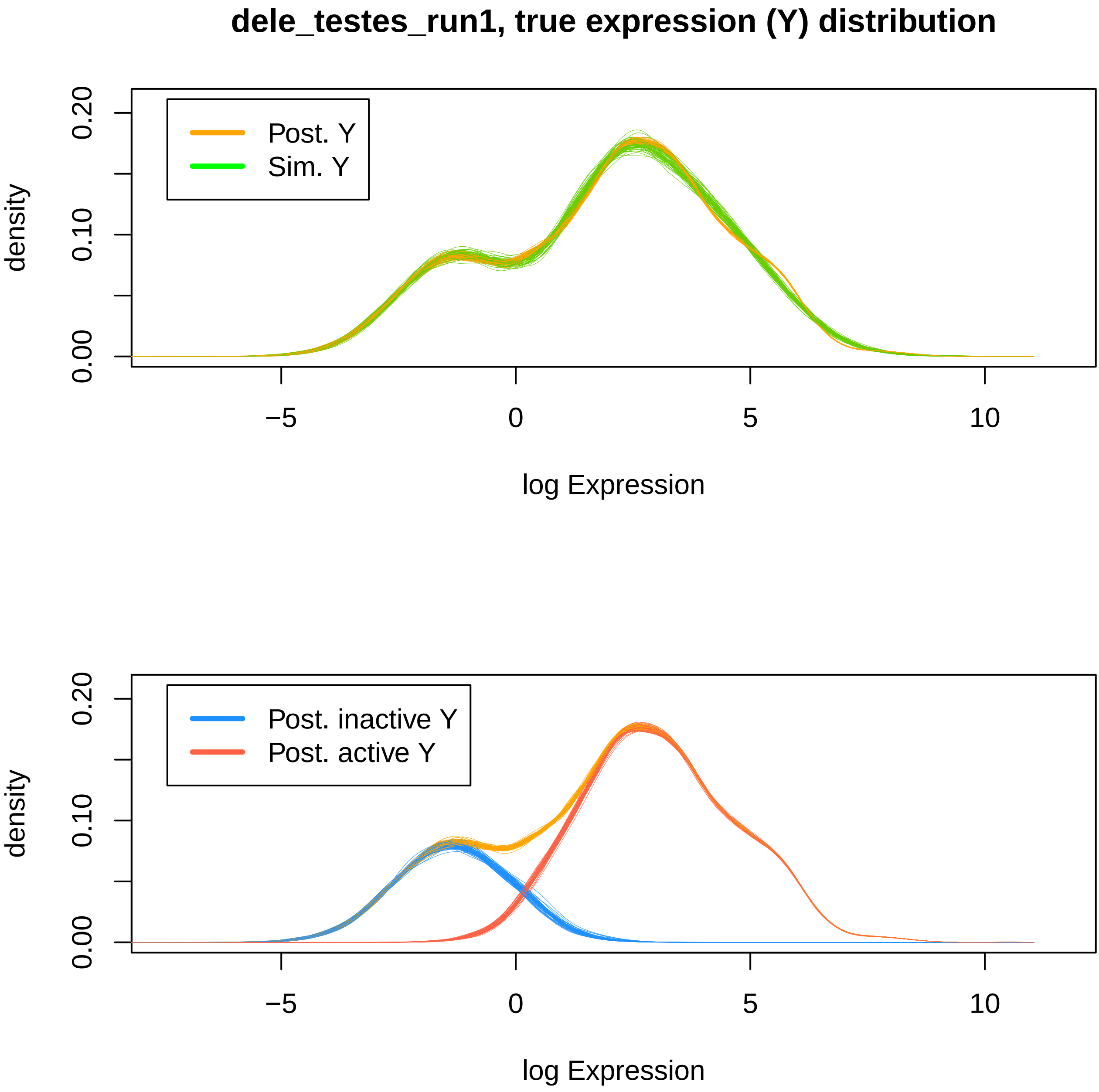
dele_testes_run1, true expression (Y) distribution
Post. Y
Sim. Y
density
0.00
0.10
0.20
−5
0
5
10
log Expression
Post. inactive Y
Post. active Y
density
0.00
0.10
0.20
−5
0
5
10
log Expression

### deug_testes_run1, true expression (Y) distribution

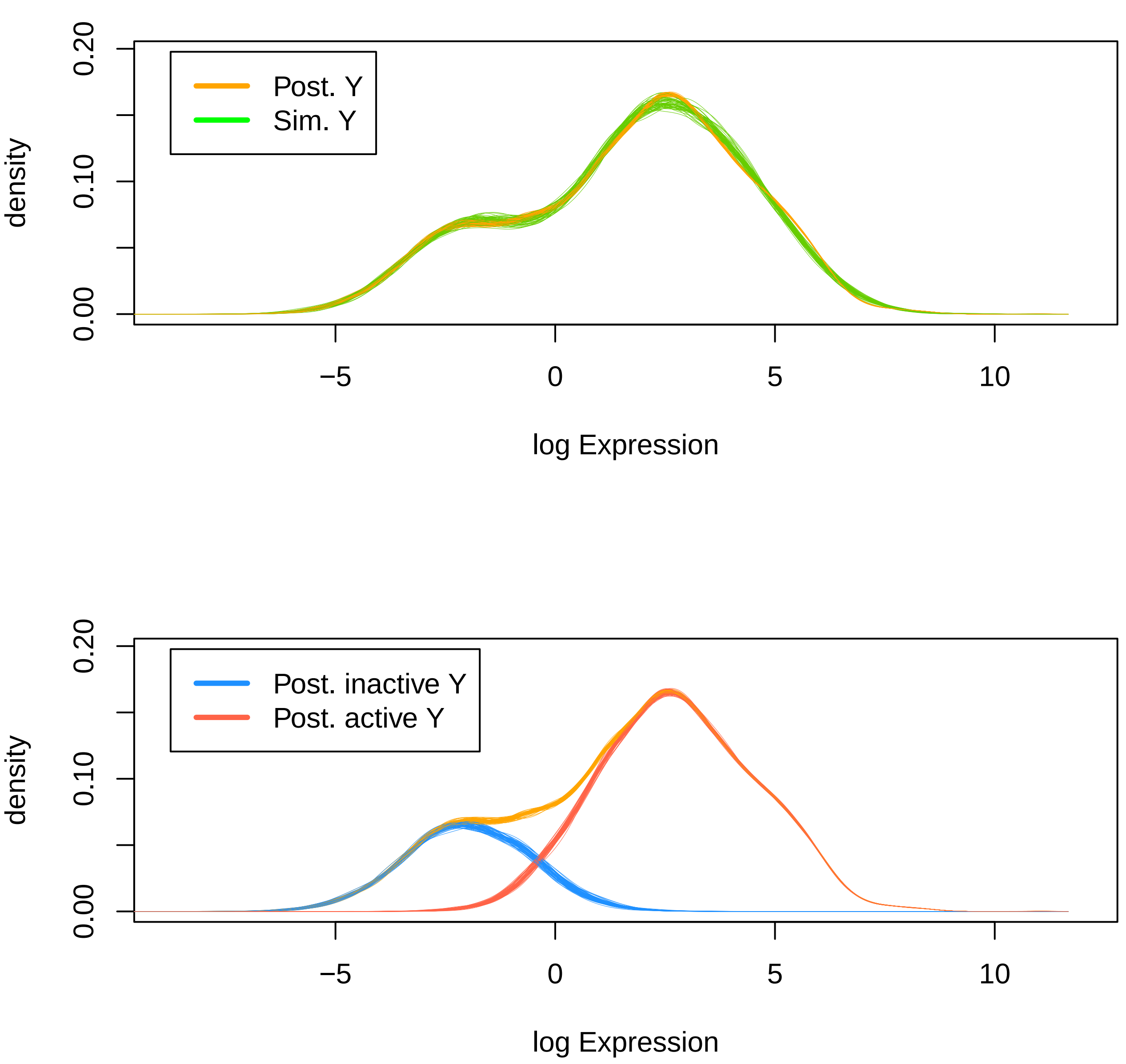

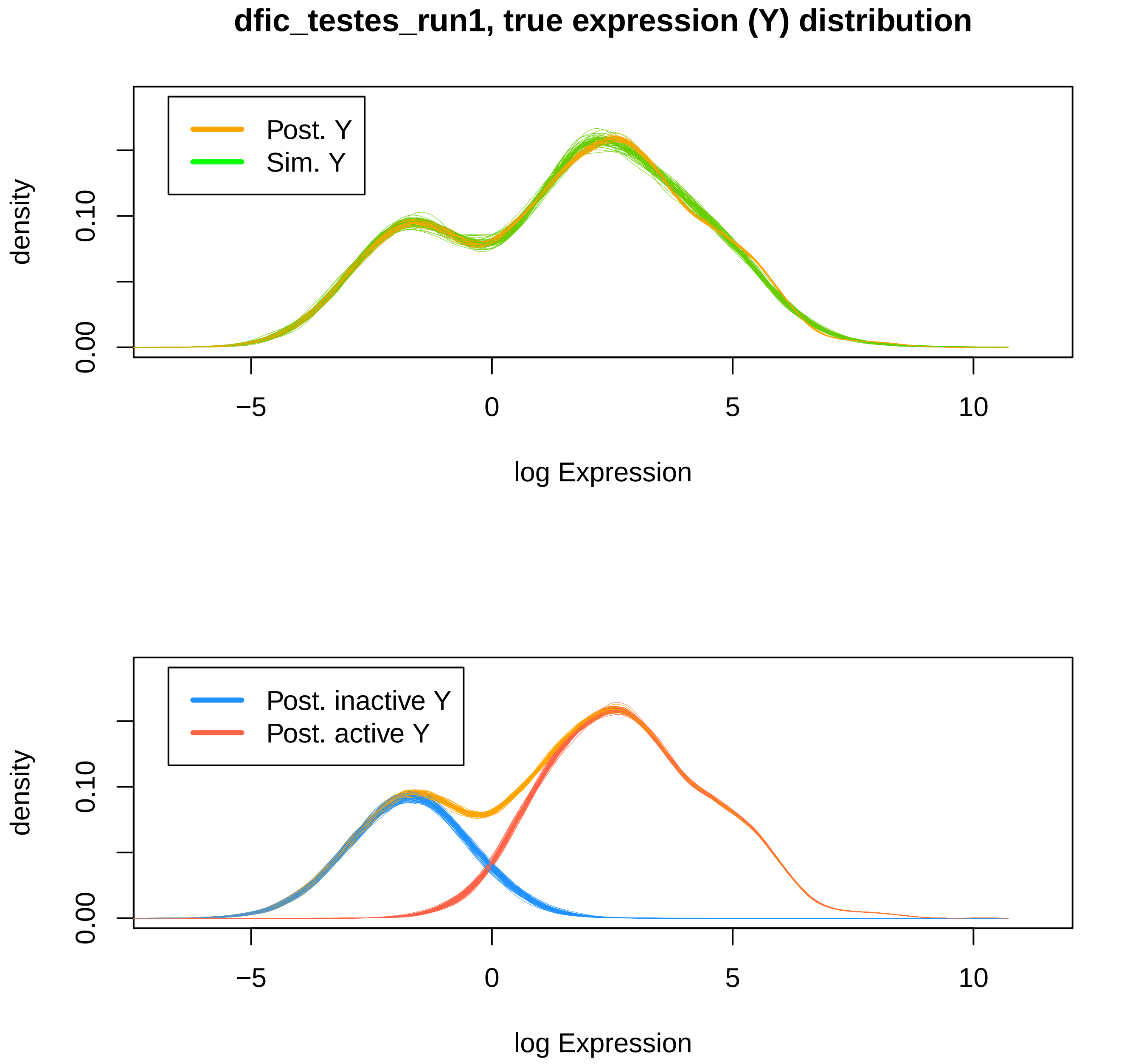
dfic_testes_run1, true expression (Y) distribution
Post. Y
Sim. Y
density
0.00
0.10
−5
0
5
10
log Expression
Post. inactive Y
Post. active Y
density
0.00
0.10
−5
0
5
10
log Expression

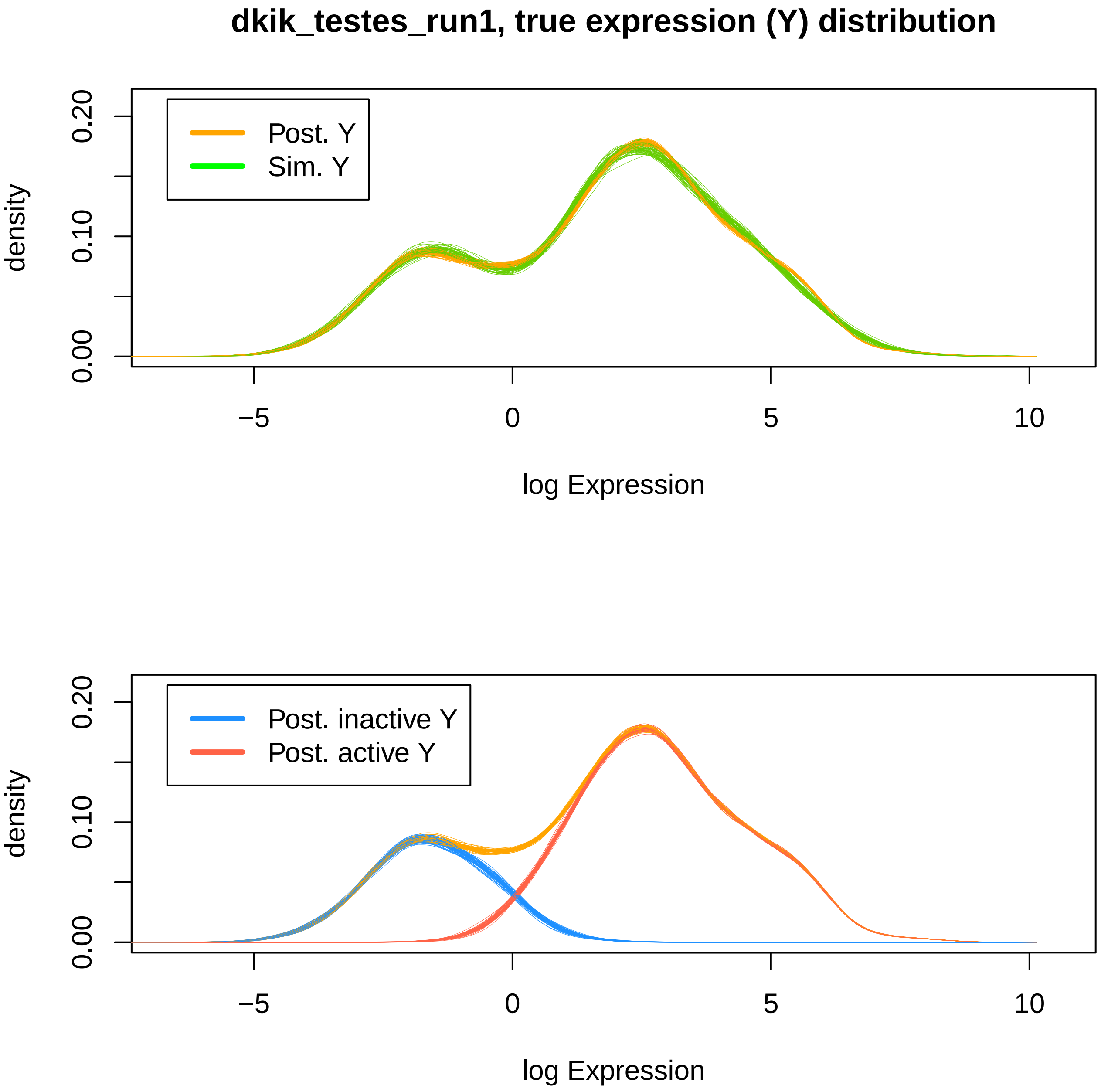
dkik_testes_run1, true expression (Y) distribution
Post. Y
Sim. Y
density
0.00
0.10
0.20
−5
0
5
10
log Expression
Post. inactive Y
Post. active Y
density
0.00
0.10
0.20
−5
0
5
10
log Expression

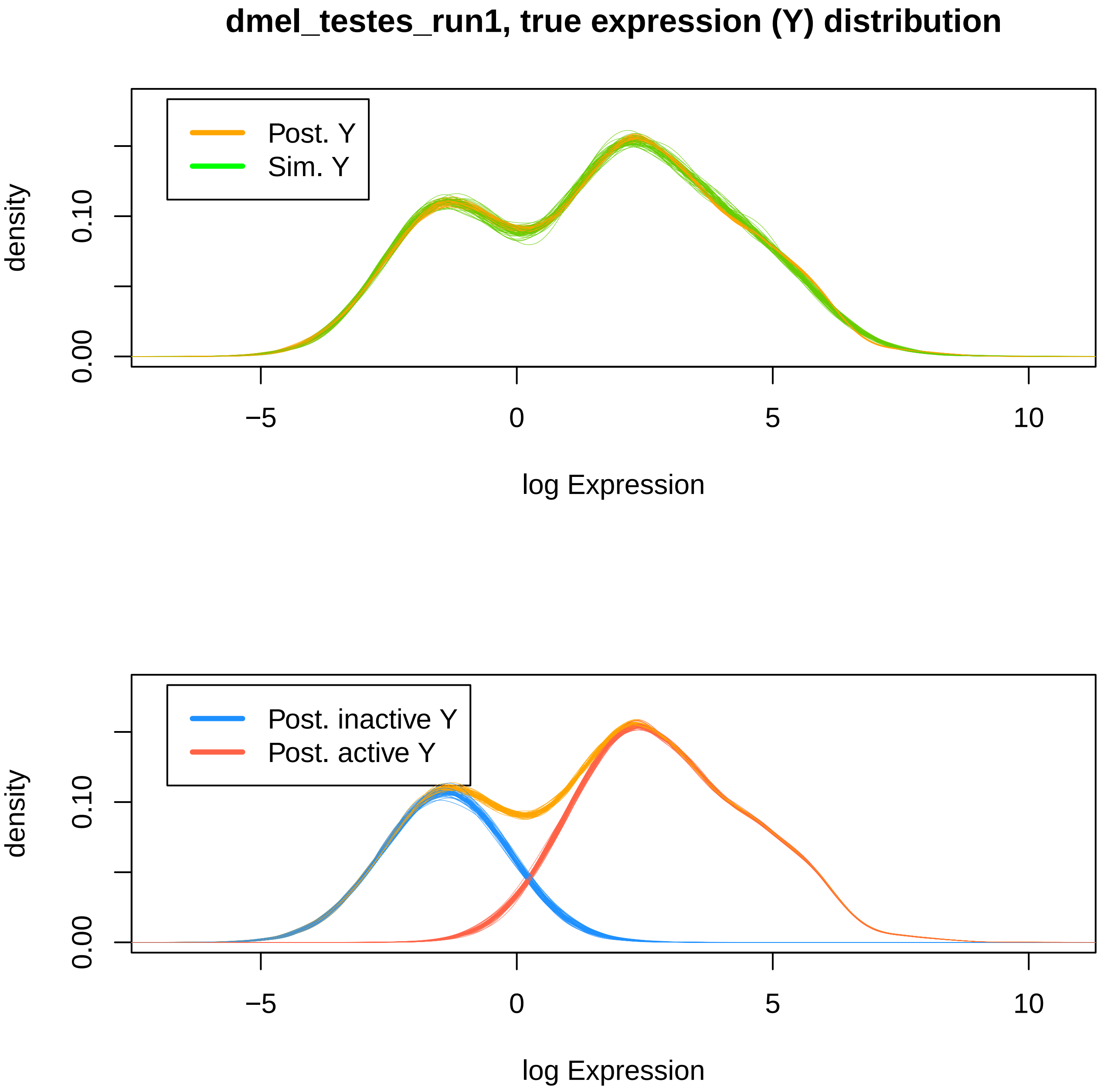
dmel_testes_run1, true expression (Y) distribution
Post. Y
Sim. Y
density
0.00
0.10
−5
0
5
10
log Expression
Post. inactive Y
Post. active Y
density
0.00
0.10
−5
0
5
10
log Expression

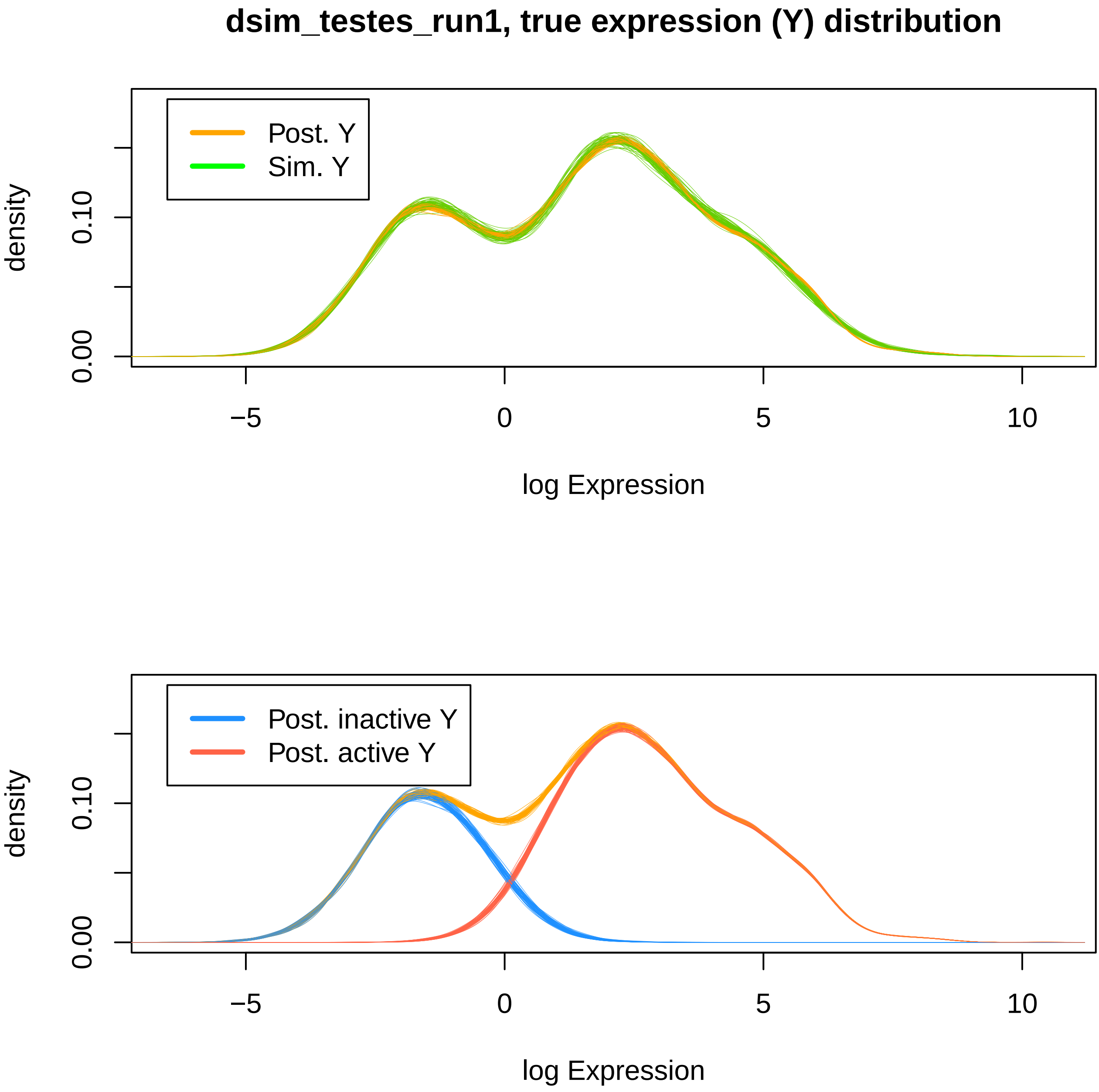
dsim_testes_run1, true expression (Y) distribution
Post. Y
Sim. Y
density
0.00
0.10
−5
0
5
10
log Expression
Post. inactive Y
Post. active Y
density
0.00
0.10
−5
0
5
10
log Expression

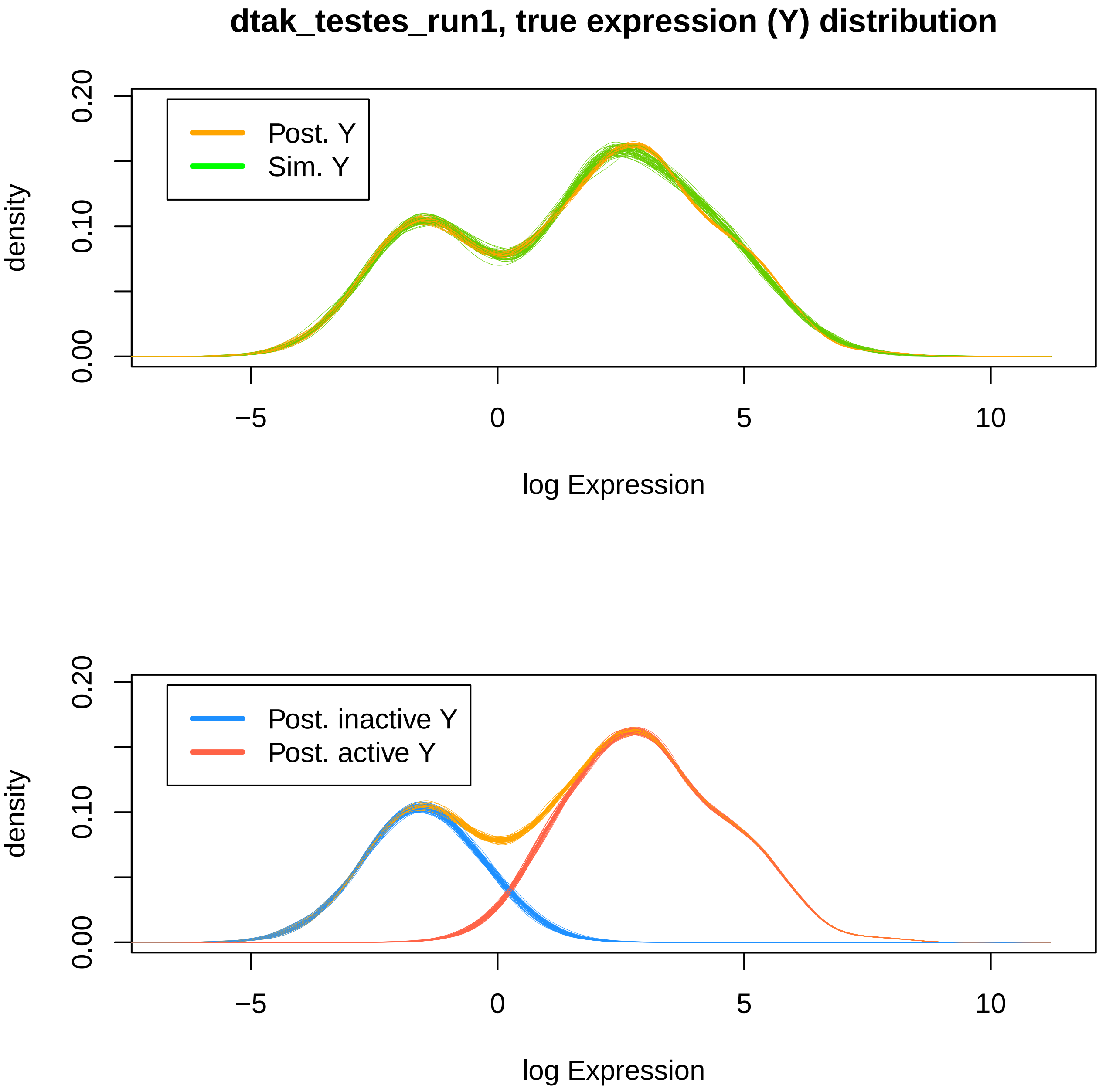
dtak_testes_run1, true expression (Y) distribution
Post. Y
Sim. Y
density
0.00
0.10
0.20
−5
0
5
10
log Expression
Post. inactive Y
Post. active Y
density
0.00
0.10
0.20
−5
0
5
10
log Expression

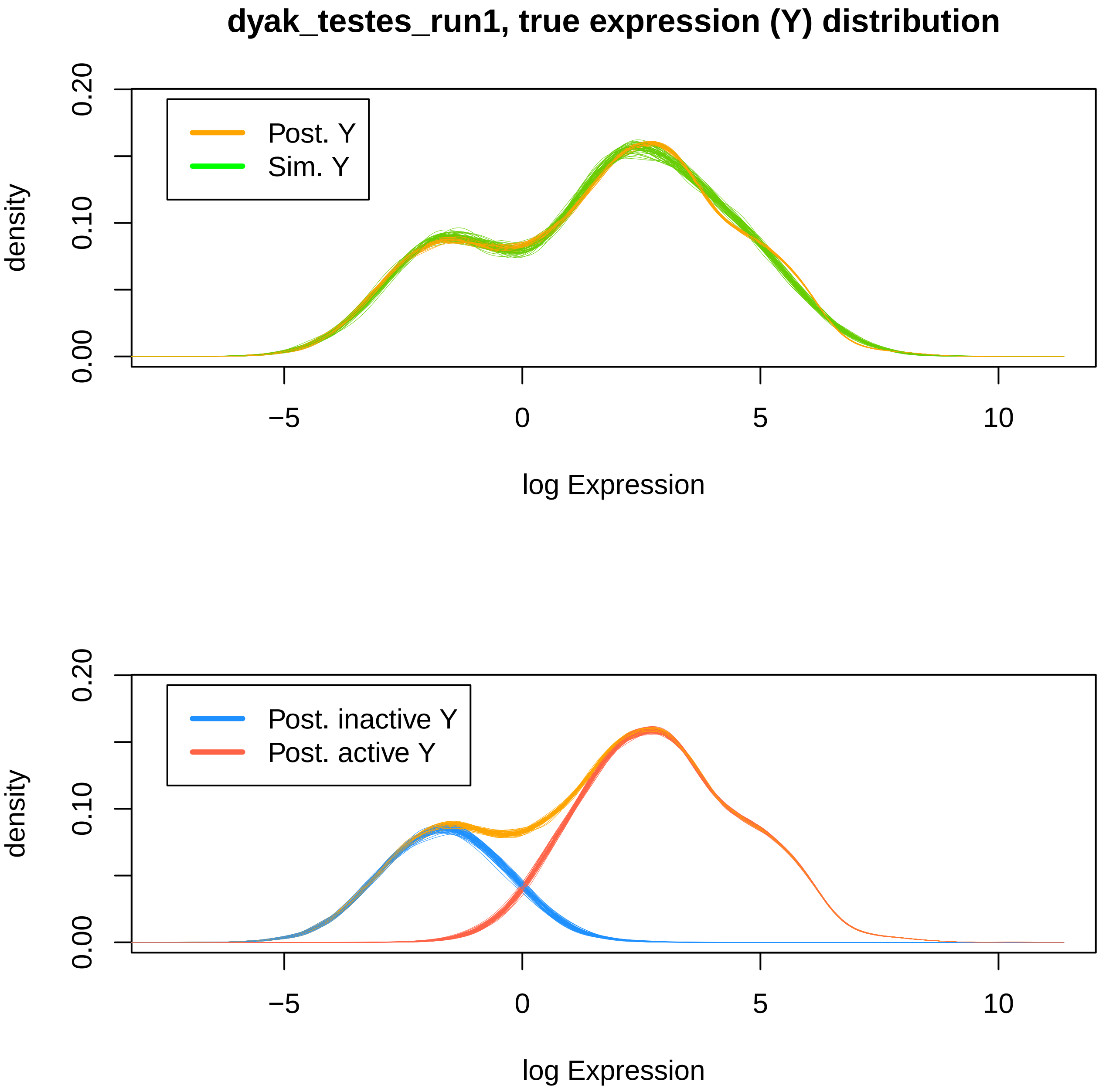
dyak_testes_run1, true expression (Y) distribution
Post. Y
Sim. Y
density
0.00
0.10
0.20
−5
0
5
10
log Expression
Post. inactive Y
Post. active Y
density
0.00
0.10
0.20
−5
0
5
10
log Expression
